# Epigenetic control of microglial developmental milestones from proliferative progenitors to efficient phagocytes

**DOI:** 10.1101/2025.11.03.685577

**Authors:** Marta Pereira-Iglesias, Duncan Martinson, Carles Falco, Joel Maldonado-Teixido, Marco González-Domínguez, Rodrigo Senovilla-Ganzo, Sol Beccari, Jorge Valero, Bella Mora-Romero, Ivan Ballasch, Sarah Viguier, Philipp Hane, Molly Boettiger, Julie A. Reisz, Alba Elías-Tersa, Yasmina Manso, Laura Parkkinen, Ana María Aransay, Federico N. Soria, Angelo D’Alessandro, Eduardo Soriano, Morgane S. Thion, Sonia Garel, Melanie Greter, Albert Giralt, Alberto Pascual, Fernando García-Moreno, David A. Menassa, Jose A. Carrillo, Amanda Sierra

**Author notes:** same contribution (Marta Pereira-Iglesias, Duncan Martinson, Carles Falco). **Correspondence Amanda Sierra, PhD** Achucarro Basque Center for Neuroscience, Parque Científico UPV/EHU, edificio sede, planta 3 Barrio Sarriena, s/n Leioa, 48940, Bizkaia, Spain; **José A. Carrillo, PhD** Mathematical Institute, University of Oxford, Andrew Wiles Building, Radcliffe Observatory Quarter, Woodstock Road, Oxford, OX2 6GG, UK.

## Abstract

Early immune perturbations increase the risk of neurodegenerative and neurodevelopmental disorders, yet the mechanisms underlying the maturation of microglia, the resident immune cells of the brain parenchyma, remain poorly defined. Specifically, how proliferation, morphological differentiation, and phagocytosis are coordinated among microglia progenitors as they colonize the embryonic brain remains unclear. Here, we combined mathematical modeling with spatiotemporal analyses of the murine hippocampus and cerebellum from postnatal day 2 (P2) to P60 to reconstruct the trajectory of microglial development. We identified a proliferative-to-quiescent (P/Q) switch around P3/P4 that preceded the acquisition of morphological complexity and efficient phagocytosis and was accompanied by coordinated shifts in cell-cycle dynamics and metabolic state. Strikingly, this P/Q switch was recapitulated in repopulation contexts in mice and in the human fetal brain, where later stages displayed enhanced phagocytic function coupled to reduced proliferation. Perturbing the proliferative phase through pharmacological or genetic disruption of CSF1R signaling impaired subsequent microglial complexity and phagocytosis efficiency, revealing an unexpected reliance of phagocytosis on proliferation-driven colonization. Finally, we show that microglia stepwise maturation during development is associated with chromatin remodeling and driven by the epigenetic regulator Ikaros. Together, these findings uncover the sequential milestones of microglial development, revealing a potential period of early vulnerability and establishing an unexpected linkage between proliferation and phagocytosis essential to understanding how these processes are coordinated in neurodegenerative disorders.

## Introduction

Microglia, the resident immune cells of the central nervous system parenchyma, were discovered in 1919 by Pio del Rio Hortega but their developmental origin evaded identification for nearly a century, with researchers supporting either a mesodermal or ectodermal lineage^1^. The controversy was only settled in the past decade, when it was demonstrated that the cells arise from primitive macrophage progenitors in the yolk sac that infiltrate the developing brain during early embryogenesis^2,3^. Interest in microglial development has since continued to rise, due to increasing evidence that has linked early immune perturbations to neurodegenerative^4^ and neurodevelopmental disorders^5^, suggesting that susceptibility to brain diseases during adulthood may be related to alterations in microglial development.

Uncovering the pathways governing microglia development remains challenging, however. Lineage–specific transcriptional regulators of early development and colonization, such as colony stimulating factor 1 receptor (CSF1R), RUNX, IRF8, TGFβ and Pu.1^3,6–12^, have been identified through fate-mapping, transcriptomic, and epigenetic studies, although the precise mechanism by which they operate on microglial progenitors remains unclear. Additionally, most studies lack the spatiotemporal resolution needed to dissect region-specific developmental trajectories and often rely on broad developmental stages or whole-brain samples^2,13^. While immunohistochemical^14^ and transcriptomic analyses^15–17^ have highlighted the striking proliferative capacity of microglia during early postnatal stages, it is still unclear how microglia transition from early amoeboid, sparsely distributed progenitors in the embryonic brain to highly ramified cells with a tessellated distribution and a homeostatic transcriptional profile in the adult brain.

More specifically, a critical yet understudied aspect of microglial maturation is how their proliferation, differentiation, and functional specialization are coordinated across their developmental trajectory. For example, the homeostatic profile of developmental microglia differs from that of adult microglia, which includes a mature sensome^15,18^ that recognizes cell debris^19^. Yet this finding conflicts with the assumption that microglia are intrinsically efficient phagocytes, an idea likely shaped by their classification within the macrophage lineage^20^. While some developing microglia have been shown engaged in phagocytosis in zebrafish, mice, and humans^21–23^, phagocytosis efficiency (i.e., the ability to rapidly and completely clear cell debris) has not been addressed. Thus, the questions of when and how microglia acquire the ability to efficiently engulf and clear debris, including cells undergoing apoptosis or programmed cell death, remain unresolved. Defining both the timing and regulatory mechanisms leading to phagocytosis maturation is crucial, as these are essential for maintaining brain homeostasis and supporting neurogenic niches^24,25^. Furthermore, failure to clear apoptotic cells leads to the release of intracellular contents, triggering local inflammation and tissue damage^26^.

Here, we addressed microglial colonization and functional maturation with a combined theoretical and experimental methodology. Specifically, we developed and calibrated a mathematical model of microglia proliferation and colonization using data from the murine hippocampus and cerebellum. Through a comprehensive spatiotemporal analysis between postnatal day 2 (P2) and P28, we discovered that microglia undergo a conserved sequence of developmental milestones, which consisted of an early phase of rapid proliferation, followed by a switch at P3-P4 to a mitotically quiescent phenotype that then reached morphological complexity and phagocytosis efficiency by P14. This proliferative-to-quiescent (P/Q) switch coincided with coordinated changes in cell-cycle dynamics and metabolic state. Disrupting the proliferative phase by pharmacologically or genetically altering the CSF1R pathway impaired the subsequent acquisition of microglial morphological complexity and phagocytosis efficiency, uncovering an unexpected reliance of phagocytosis on colonization. Finally, we demonstrated that microglial maturation is accompanied by chromatin remodeling and driven by the epigenetic regulator Ikaros. Together, our data reveals microglia developmental milestones as they colonize the brain, exposing a potential early period of vulnerability and a previously unidentified inverse relationship between proliferation and phagocytosis, which will be crucial to further understanding the role of microglia in neurodegenerative disorders.

## RESULTS

### Data-driven modeling suggests microglial maturation involves a switch from proliferation to mitotic quiescence

We first determined how microglial colonization of the brain parenchyma was orchestrated using mathematical modeling of microglial density and proliferation along a time course from postnatal day (P) 2 to P28 in fms-EGFP mice, in which microglia express the green fluorescent reporter^27^ (**Figure 1**). We focused on the hippocampus and cerebellum because these two regions have protracted postnatal development, allowing us to disregard cells incoming from the yolk sac^28,29^; and because they are well-defined anatomically, allowing us to quantify their respective volumes over time. In the hippocampus, microglial density increased steadily from P2 to P14 (**Figure 1A, B**), after which it declined due to continued hippocampal expansion (**Figure 1C**). Total microglia numbers were unaltered from P21 onwards (**Figure 1D**), indicating that colonization was completed at this stage. Notably, microglial expansion occurred earlier in the Cornu Ammonis (CA) region compared to the dentate gyrus (DG) (**Supp. Figure 1A-F**), suggesting region-specific timing in colonization dynamics. In the cerebellum, microglial density was consistently lower than in the hippocampus, in agreement with previous reports^30^. Microglia density increased until P10, followed by a decline as cerebellar volume expanded through P60 (**Supp. Figure 1G-J**). Final microglial numbers were reached earlier in the white matter (WM) compared to the granular and molecular layers, indicating a spatial gradient in cerebellar microglial colonization (**Supp. Figure 1K-S**). To determine the role of proliferation in microglial expansion in the hippocampus and cerebellum, we analyzed the expression of the proliferation marker Ki67^31^. In the hippocampus, we observed a high proportion of proliferating microglia at the earliest time points, with 41.3L±L2.6% microglia expressing Ki67 at P2. This proportion declined sharply by P7 (6.8L±L0.3%) and remained low thereafter (**Figure 1E, F**). Despite this decline in the proliferation rate, the small percentage of proliferative microglia found between P7-P14, applied to an expanding microglial population, led to relative constant numbers of proliferative microglia up until P14 (**Figure 1G; Supp.** Figure 2A-D). A similar proliferative trajectory was observed in the cerebellum (**Supp. Figure 2E-M**), with early postnatal peaks in microglial proliferation followed by a progressive decline, revealing a conserved temporal profile of microglial proliferation across brain regions.

**Figure 1.**
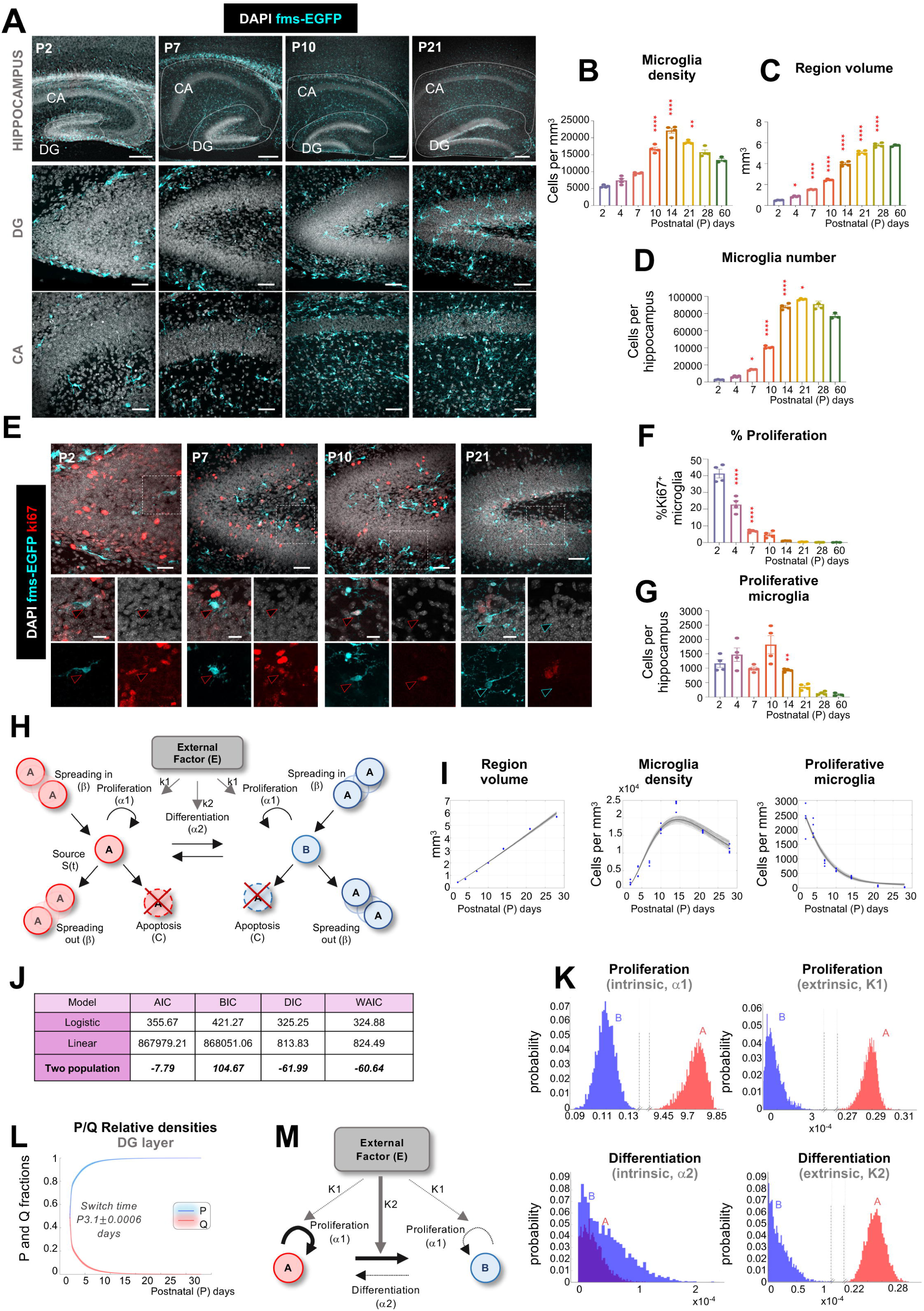
Mathematical modeling of hippocampus and cerebellum development identifies a coordinated switch from proliferative to mitotically quiescent microglia. (**A**) Representative confocal images showing the hippocampus at P2, P7, P10, and P21 in fms-EGFP mice. DAPI (white) staining shows nuclei and GFP (cyan) microglia. (**B**) Density of microglia in the hippocampus. (**C**) Hippocampal region volume. (**D**) Total microglia number per hippocampal volume. (**E**) Representative confocal images showing the hippocampus at P2, P7, P10, and P21 in fms-EGFP mice. Proliferation was identified by Ki67 (red), microglia as GFP positive cells (cyan), and nuclei with DAPI (white). High magnification images show examples of proliferative (red arrows) and non-proliferative (cyan arrows) microglia. (**F**) Percentage of proliferative microglia in the hippocampus. (**G**) Total number of proliferative microglia in the hippocampus. (**H**) The two-population model using differential equations allowed microglia to distribute into two different mathematical populations, A and B, and accounts for four possible interactions: proliferation (α1), differentiation (α2), spreading between regions (β), and apoptosis (**C**). Proliferation and differentiation can occur at intrinsic (K_1_) or extrinsic rates (K_2_) in the model, the latter of which is modulated by an environmental factor that is assumed to correlate with the sub-region volume (E). (**I**) Structure volume, microglial density and proliferation in the hippocampus. Dots represent the raw data and the blue line and grey shaded area the model estimation. (**J**) Mathematical criteria to determine model optimal fitness: Akaike information criterion (AIC), Bayesian information criterion (BIC), and Watanabe-Akaike information criterion (WAIC). Lower scores indicate better fits to the model. (**K**) Estimated marginal posterior distributions of the intrinsic and environmental dependent proliferation and differentiation parameters by Markov Chain Monte Carlo (MCMC) in the hippocampus. (**L**) Relative density of the proliferative and quiescent populations in the DG. (**M**) Schematic summary of the main results of the two-population model, showing that population A is highly proliferative compared to population B, and it’s a proliferation rate largely driven by intrinsic cues. Population A differentiates into population B driven by extrinsic cues, whereas population B does not differentiate back into A. Bars show mean±SEM of n=4 mice (B-D, F, G). Data were analyzed by one-way ANOVA followed by Tukey post hoc test, *p<0.05, **p<0.01, ***p<0.001. Only significant differences between consecutive ages are shown. Scale bars (**A**): upper row, 100µm; middle and bottom row, 50µm. Thickness, left to right, z= 10.5µm, 13.3µm, 17.5µm, 11.2µm (upper row); z=21.7µm, 21µm, 16.8µm, 16.1µm (middle row); z=21.7µm, 18.9µm, 21µm, 16.8µm (bottom row). Scale bars: (**E**) upper row, 50µm; bottom row, 20µm. Thickness, left to right, z=21µm, 21.7µm, 19.6µm, 18.9µm; bottom row z=2.1µm, 2.1µm, 2.1µm, 2.1µm.

To gain greater insight into how microglia coordinate proliferation, apoptosis, and migration to colonize these brain regions, we developed a series of increasingly complex mathematical models using ordinary differential equations and calibrated them to the above data (density of microglia, percentage of proliferative microglia, brain region volume) using Markov chain Monte Carlo (MCMC) parameter estimation (**Supp. Figure 3A-D; Supp. Text**).

Focusing first on the cerebellum because of its paradigmatic non-monotonic (increasing-then decreasing) pattern in microglial density across its layers, we tested a minimal logistic growth model in which the density of microglia depended on the balance between net proliferation/apoptosis and migration out of the region. This model accurately estimated the volume and total microglial density, but not the density of proliferative (Ki67^+^) microglia (**Supp. Figure 3A, B**). Therefore, we next developed a more complex linear model that accounted for an incoming source of cells that could further contribute to the increasing microglia cell density over time, but this model also failed to describe microglial proliferation (**Supp. Figure 3C, D**). Hence, we increased again the complexity of the model using data from all hippocampal and cerebellar sub-regions and allowed microglia to distribute into two mathematical populations that could differ in their respective rates of proliferation, differentiation, apoptosis, and spreading between sub-regions. In each sub-population, proliferation and differentiation could respectively occur at a baseline, “intrinsic” rate in which cells spontaneously proliferated or transformed into the other type. Proliferation and differentiation could also occur at separate rates that were controlled by an environmental factor that we assumed was coupled to the volume of the brain region. The two-population model, which provided the best fit based on three independent model selection criteria, robustly captured the observed dynamics of regional volume, microglial density, and proliferation across all layers of the hippocampus and cerebellum (**Figure 1H-J; Supp.** Figure 3E-I).

To identify the biological relevance of the two mathematical populations identified by the mathematical model, we estimated the underlying model parameters governing proliferation, differentiation, apoptosis, and spreading using a Markov chain Monte Carlo (MCMC) sampling algorithm. In both the cerebellum and the hippocampus, population A was highly proliferative compared to population B and its proliferation rate was largely driven by intrinsic cues. Additionally, extrinsic cues drove population A to differentiate into population B, but population B did not differentiate back into A (**Figure 1K; Supp.** Figure 4A**, B**). Hence, the model suggested that microglia behaved as two different subpopulations: one consisting of proliferative progenitors (P) that expand cell-autonomously, and a quiescent (Q) one into which progenitors transform according to signals from the expanding brain structure.

We next dissected the temporal dynamics underlying the switch between P and Q microglia by quantifying their relative abundance over time. The model predicted that these two populations did not coexist: microglia remained highly proliferative until P3– P4, after which they synchronously exited the cell cycle and switched into a mitotically quiescent state (**Figure 1L**). Notably, the times at which microglia were predicted to switch from a majority-P state to a majority-Q state were similar regardless of whether the mathematical model was calibrated to data either from hippocampal layers or from cerebellar layers (**Supp. Figure 4B**). The earliest P/Q switched occurred in the DG region of the hippocampus (3.1L±L0.0009 days) and the latest in the granular layer of the cerebellum (4.2L±L0.001 days). It is important to note that the same mathematical model provided near-identical trends despite having different region-specific (i.e., hippocampus and cerebellum) parameter values dictating the respective time scales of proliferation, differentiation, and apoptosis. Thus, the two-population model identified a milestone of microglial development that was not evident from the observation of their population dynamics (**Figure 1M**): specifically, a synchronous switch from proliferative to mitotically quiescent cells (P/Q switch) that takes place during early development in the hippocampus and cerebellum, around P3-P4, long before they reach their final density around P30-60.

Finally, we assessed the remaining parameters using MCMC sampling. In both the cerebellum and hippocampus, predicted apoptosis rates were low compared to predicted proliferation rates, with only few cells undergoing apoptosis in the first few postnatal days (**Supp. Figure 4C, D**). This result is consistent with the coupling between microglial proliferation and apoptosis in the adult brain^32^ but suggests that microglial apoptosis does not play a major role in their colonization. We then analyzed the role of migration between sub-regions (spreading) by comparing models that permitted spreading only in specific directions. In the hippocampus, the best fit to the data occurred when microglia invaded from outside the hippocampus into the CA and then spread inward towards the DG (**Supp. Figure 4E, F**). In contrast, in the cerebellum the best fit occurred when microglia from the white matter spread outwardly towards the granule cell layer and then to the molecular layer (**Supp. Figure 4G, H**). These contrasting migration routes suggest that local microenvironmental cues orchestrate region-specific microglial positioning during development. To test this hypothesis, we analyzed microglial maturation in mice deficient for Reelin, a scaffold peptide for neuronal migration^33,34^. Mice with inducible ubiquitous reelin depletion (*UbiCre-ERT2-Reelin^Flox/Flox^* treated with tamoxifen at P1-P3) showed a reduction in the number of microglia in the cerebellum by P7 that recovered by P14, and that was not accompanied by changes in proliferation, suggesting a transient alteration in spreading that did not affect their final colonization (**Supp. Figure 5A-E**). However, no effects of reelin deficiency were found in the hippocampus (**Supp. Figure 5F-H**). These findings underscore that microglial colonization is not solely a function of cell-intrinsic programming but is dynamically tuned by the environment. In addition, they suggest that apoptosis and spreading between subregions have minor roles in colonization, and that proliferation is the major driver of the microglia population expansion.

### Changes in cell cycle dynamics and metabolism accompany the P/Q switch

To further validate the two-population mathematical model, we tested its prediction that the doubling time of the microglial population lengthened over time, implying an increase in cell cycle duration with age (**Figure 2A; Supp.** Figure 4I). To analyze cell cycle, we used a dual thymidine analog labeling approach with sequential injections of EdU and BrdU (5-ethynyl-2’-deoxyuridine and 5-bromodeoxyuridine, respectively, 50mg/kg each) in P2 and P7 fms-EGFP mice at different intervals. Microglia that re-entered the cell cycle were identified as EdU^⁺^/BrdU^⁺^ cells, and the peak of double-labeled cells indirectly indicated the cell cycle duration^35^. Consistent with model predictions, we observed a significant age-dependent decline in the proportion of double-labeled microglia in both the hippocampus (**Figure 2B, C**) and the cerebellum (**Supp. Figure 4 J, K**). At P2, the peak of double labeled microglia was found at 12h (19.1L±L0.3% EdU^⁺^/BrdU^⁺^ microglia in hippocampus; 34.1L±L2.7% in cerebellum), whereas by P7, the peak of EdU^⁺^/BrdU^⁺^ microglia decreased markedly and was found at 24h (6.7L±L0.6% EdU^⁺^/BrdU^⁺^ microglia in hippocampus; 5.3L±L1.0% in cerebellum). This increased duration of microglial cell cycle from 12h at P2 to 24h at P7 mirrors similar dynamics observed in neural progenitor cells, where cell cycle elongation accompanies the transition from expansion to differentiation^36^. Together, these data confirm the model’s prediction of increased doubling time and reveal a developmental switch in microglial proliferative behavior. Together, our model shows that brain colonization is driven by a small cohort of fast cycling microglia.

**Figure 2.**
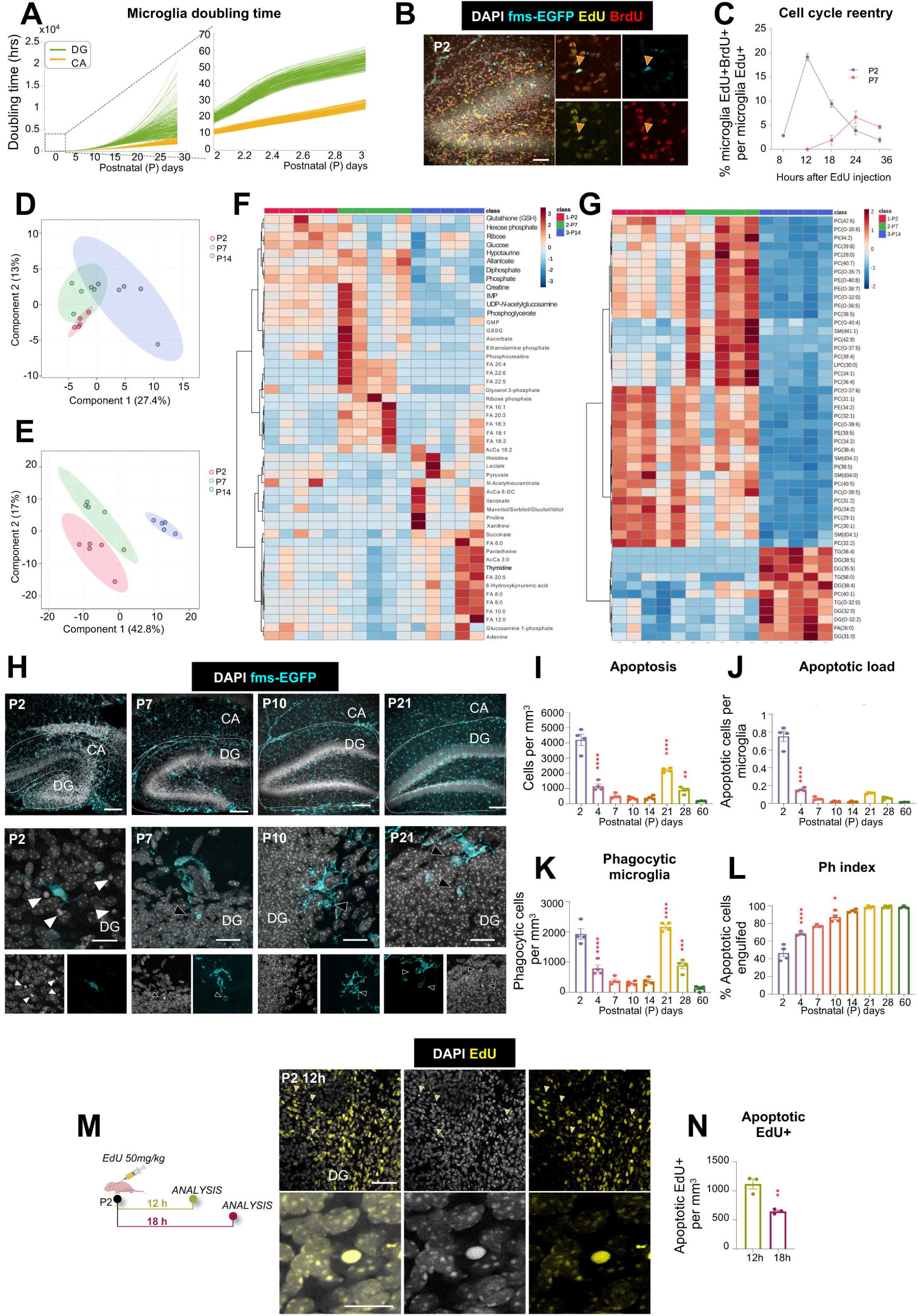
Microglia undergo changes in cell cycle dynamics and metabolism around the P/Q switch. **(A)** The two-population model predicts the doubling time of the microglia population increases with age in the hippocampus. (**B**) Representative confocal images of the hippocampus from P2 fms-EGFP mouse injected with EdU and BrdU (8h apart) DAPI (white) staining shows nuclei, EdU (yellow) cells at t=0 and BrdU (red) cells at t=8h. High magnification show EdU cells (yellow arrow) and EdU^+^BrdU^+^ cells (orange arrow). (**C**) Percentage of EdU^+^BrdU^+^ microglia at P2 and P7 in the hippocampus. (**D, E**) partial least squares-discriminant analysis (PLS-DA) of metabolomics (**D**) and lipidomics (**E**) composition FACS-sorted microglia at P2, P7 and P14 in fms-EGFP mice. (**F**) Heatmap of the top 50 metabolites by p-value (one-way ANOVA) in microglia at P2, P7 and P14. (**G**) Heatmap of the top 50 lipids by p-value (one-way ANOVA) in microglia at P2, P7 and P14. (**H**) Representative confocal images showing the hippocampus at P2, P7, P10, and P21 in fms-EGFP mice. Apoptotic cells were identified by their condensed nuclear morphology detected by DAPI (white) and microglia as GFP positive cells (cyan). High magnification images show non-engulfed (P2, white arrow) and engulfed (P7, P10, P21, black arrows). (**I**) Density of apoptotic cells in the hippocampus. (**J**) Clearance ratio (ratio of apoptotic cells per microglia in the hippocampus). (**K**) Total number of phagocytic cells per hippocampus. (**L**) Phagocytic index (% of apoptotic cells phagocytosed by microglia) in the hippocampus. (M) Representative confocal image of an EdU^+^ apoptotic cell. Experimental design drawing created with BioRender.com. (**N**) Density of EdU^+^ apoptotic cells 12 and 18 hours after EdU injection. Bars show mean±SEM of n=3 (C, N), n=4 (I-L) mice. Data were analyzed by one-way ANOVA followed by Tukey post hoc test, *p<0.05, **p<0.01, ***p<0.001. Only significant differences between consecutive ages are shown. Scale bar (**B**): 50µm and 20 µm (detail). Thickness z=10.5µm and 2.1µm. Scale bars (**H**): upper row, 100µm; bottom row 20µm. Thickness, left to right: z=12.25µm, 11.2µm, 10.15µm, 27.3µm (upper row); z=2.1µm, 2.1µm, 2.1µm, 2.1µm (bottom row).

We then confirmed the shift in proliferation dynamics by assessing changes in microglial metabolism using mass spectrometry from FACS-sorted microglia. For robustness, and due to the largely similar time course of microglial development between hippocampus and cerebellum, we pooled microglia from the two regions. As they switched from highly proliferative progenitors (at P2) to mitotically quiescent cells (at P14), microglia displayed changes in their metabolic composition (**Figure 2D; Supp.** Fig. 6), and more robust changes in their lipidomic profile (**Figure 2E; Supp.** Fig. 6). Compared to P2 microglia, P7 microglia were characterized by lower levels of glucose and hexokinase product hexose phosphate; elevated levels of later glycolytic intermediates (e.g. fructose bisphosphate, phosphoglycerate) and pentose phosphate pathway product ribose phosphate, suggestive of glucose carbon rerouting to favor the PPP over glycolysis. Additionally, reduced glutathione (GSH) was decreased at P7 while Krebs cycle metabolites were elevated, though not significantly. Of the 11 unsaturated fatty acids measured, 8 were significantly increased in microglia at P7 compared to P2. Interestingly, at P14, microglial free fatty acid composition changed entirely, favoring the accumulation of saturated medium chain fatty acids (e.g., C6-C10) while unsaturated fatty acids were restored to the P2 levels. Also of note were observed changes at the level of nucleotides: GMP, IMP, allantoate, and urate increased from P2 to P7, then decreased below P2 levels. (**Figure 2F; Supp.** Fig. 6). A class-level analysis of the top 50 lipids suggests a clear shift from phospholipids at P2 and P7 to di- and triglycerides at P14 (**Figure 2G**). Together, these results suggest a complex remodeling of microglial metabolism progressively around the P/Q switch as they transition from highly proliferative cells rich in multiple metabolic pathways and signaling/structural lipids, to mitotically quiescent cells that accumulate energy lipids.

### Microglia inefficiently phagocytose apoptotic cells during early postnatal development

We next tested the functional implications of the metabolic changes associated with the P/Q switch by assessing the ability of microglia to efficiently phagocytose apoptotic cells in the hippocampus and cerebellum. In the hippocampus, apoptotic cells, defined as cells with pyknotic/karyorrhectic nuclear morphology (a gold-standard in apoptosis assessment), peaked at P2 and P21 (**Figure 2H, I**). The P21 peak was specific to the dentate gyrus (**Supp. Figure 7A-D**), likely related to the establishment of the adult neurogenic cascade. We next evaluated the apoptotic load ratio (number of apoptotic cells per microglia), which remained below one through the time course (**Figure 2J**), in sharp contrast to the apoptotic load ratio that adult microglia can effectively clear (up to 4 apoptotic cells per microglia)^37^. However, the density of microglia engaged in phagocytosis (i.e., displaying phagocytic pouches containing apoptotic cells) did not match the apoptotic cell density up until P21 (**Figure 2K**). As a result, microglial phagocytosis efficiency, determined by the Ph index (percentage of apoptotic cells engulfed by microglia), was lower during early developmental stages and only reached adult levels by P14 (**Figure 2L**), suggesting an insufficient recruitment of phagocytic microglia during early postnatal development.

For example, at P2 each hippocampal microglia faced 0.72 ± 0.04 apoptotic cells (**Figure 2J**) but only 33.63 ± 3.02% of microglia were engaged in phagocytosis (**Supp. Figure 7E**), resulting in over 50% of apoptotic cells remaining un-engulfed (**Figure 2L)**. We observed a similar increase in phagocytosis efficiency in the cerebellum across postnatal development, although the cerebellum had a single peak of apoptosis at P2, and the maximum phagocytosis efficiency was reached at P21 (**Supp. Figure 7F-O**). The comparison of hippocampus and cerebellum points to a conserved developmental trajectory of functional maturation across brain regions with region-specific temporal milestones.

We then tested whether astrocytes compensated for the limited microglial phagocytic efficiency during early development, as they do in the absence of functional microglia in the adult brain^38^. We found no evidence of compensatory astrocytic phagocytosis in the hippocampus or cerebellum during the early postnatal period (**Supp. Figure 8**), underscoring microglia as the main phagocytes in the early postnatal brain. Together, these findings indicate that while some individual microglia engage in phagocytosis at early developmental stages, the microglial population does not respond efficiently to apoptotic cells before P14 in the hippocampus and P21 in the cerebellum, suggesting a delayed maturation of microglial phagocytosis efficiency.

We reasoned that low phagocytosis efficiency during early development would affect the clearance time of apoptotic cells. To experimentally assess the kinetics of microglial clearance, we resorted to analyzing apoptosis in the newborn cell population based on our previous findings that most newborn neurons in the adult hippocampal neurogenic niche undergo apoptosis^24^. We labeled proliferative cells at P2 with the thymidine analog EdU and quantified EdU^+^ cells undergoing apoptosis at 12 and 18 hours post-injection in the hippocampus (**Figure 2N, M**). The disappearance of newborn, EdU^+^ apoptotic cells in the 6-hour period allowed us to calculate their clearance time, which was 3.2L±L0.3 hours, significantly slower than the 1.2–1.5 hours reported in adults^24^. These findings confirm that reduced microglial phagocytosis efficiency translates into delayed apoptotic cell clearance.

To further confirm that early microglia were not efficient phagocytes, we challenged them with an increased apoptotic load by injecting mice with low (2.5 mg/kg) or high-dose (6 mg/kg) ethanol at P2, P7, P9, and P14 (**Figure 3A**). Both doses of ethanol induced a consistent rise in hippocampal apoptosis at all ages (**Figure 3B**), increasing the apoptotic load per microglia (**Figure 3C**) without affecting microglial numbers (**Supp. Figure 9A).** However, it was only by P14 that enough microglia engaged in phagocytosis (**Figure 3D**) to maintain their efficiency during the ethanol challenge compared to untreated microglia (**Figure 3E**). P14 microglia used the same strategy as adult cells^39^ to cope with the apoptotic challenge by expanding their phagocytic capacity, i.e., the percentage of microglia with one or more phagocytic pouches (**Figure 3F; Supp.** Figure 9B-H), thus maintaining tight coupling between apoptosis and phagocytosis (**Figure 3G**). At earlier ages, microglia exhibited an insufficient increase in their phagocytic capacity and failed to preserve the Ph index, indicating that the ability to proportionally scale phagocytosis to demand is acquired only at P14 (**Supp. Figure 9B-H**). Similar findings in the cerebellum (**Supp. Figure 9I-W**), with insufficient engagement of microglia in phagocytosis prior to P14, point to a shared maturation program. These data identify P14-P21 as a critical time window for functional maturation, in which microglia become dynamically responsive to apoptotic challenges after the P/Q switch has occurred.

**Figure 3.**
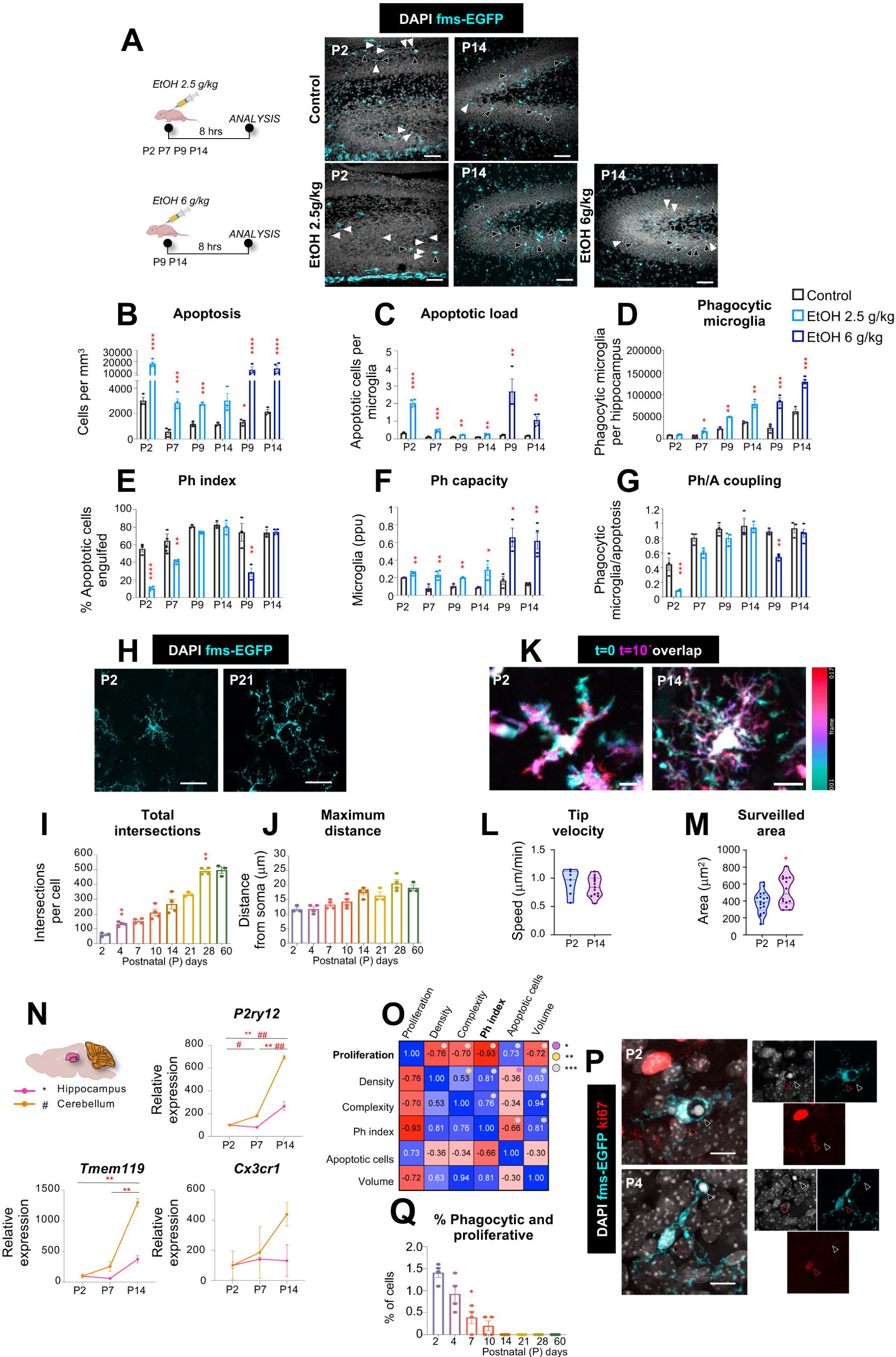
Microglial phagocytosis efficiency increases over time, as microglia matures morphologically and transcriptionally. (**A**) Representative confocal images showing the hippocampus at P2 and P14 in fms-EGFP mice in control conditions and after IP ethanol injection of 2.5g/kg or 6g/kg. Apoptotic cells were identified by their condensed nuclear morphology detected by DAPI (white) and microglia as GFP positive cells (cyan). Black arrows indicate microglia phagocytosis and white arrows indicate apoptotic cells not engulfed. (**B**) Density of apoptotic cells in the hippocampus, (**C**) Clearance ratio (ratio of apoptotic cells per microglia in the hippocampus). (**D**) Total numbers of phagocytic microglia in the hippocampus. (**E**) Phagocytic index in the hippocampus (% of apoptotic cells phagocytosed by microglia). (**F**) Phagocytic capacity in the hippocampus (proportion of microglia with one or more phagocytic pouches). (**G**) Coupling between phagocytosis and apoptosis in the hippocampus. Bars show mean±SEM of n=4 mice. Data were analyzed by multiple t-test, *p<0.05, **p<0.01, ***p<0.001 comparing vehicle (black) vs. ethanol injected (blue) mice. Experimental design drawing created with BioRender.com. (**H**) Representative confocal images of microglia (cyan) in the hippocampus at P2 and P21 in fms-EGFP mice. (**I**) Total number of intersections (reflecting the number of processes) (**J**) and maximum distance of the processes in hippocampal microglia. (**K**) Representative 2-photon time series (10 min, color-coded) of microglia in the hippocampus at P2 and P14. (**L**) Velocity of microglial process tips. (**M**) Cumulative area surveilled in the 10 min interval. (N) Expression levels from Pu.1^+^ sorted microglia at P2, P7, and P14 in the hippocampus and cerebellum of fms-EGFP mice Relative expression of P2ry12, Tmem119, and Cx3cr1. (**O**) Multiple correlations in the hippocampus showing the correlation coefficient R between proliferation, density, complexity (number of processes x distance), phagocytic index, density of apoptotic cells, and volume. (**P**) Representative confocal images of proliferative and phagocytic microglia in the hippocampus at P2 and P4 in fms-EGFP mice. Proliferative microglia were identified by Ki67 expression (red) and microglia as GFP positive cells (cyan). DAPI (white) staining shows nuclei. White arrows indicate an apoptotic cell engulfed and red arrows indicate proliferative microglia. **(Q)** Percentage of proliferative and phagocytic microglia over total microglia in the hippocampus. Bars show mean±SEM of n=3 control, n=4 ethanol injected (B-G), n=4 (I-J, R), n=3 (O) mice. in L-N, each dot represents an individual cell, with n= 17 cells from 9 animals at P2, and n=14 cells from 7 animals at P14. Data were analyzed by multiple t-test, *p<0.05, **p<0.01, ***p<0.001 comparing vehicle (black) vs. ethanol injected (blue) mice (**B-G**), or by one-way ANOVA followed by Tukey post hoc test, *p<0.05, **p<0.01, ***p<0.001 (**I, J, O, R**). Significant correlations are shown by violet dots (p<0.05), tangerine dots (p<0.019, and gray dots p<0.001). Only significant differences between consecutive ages are shown. Experimental design drawing created with BioRender.com. Scale bars (**A**): 50µm. Thickness, left to right z=10.5µm, 12.5µm; z=13.3µm, 11.2µm, 18.2µm. Scale bars (**H**): 20µm. Thickness, left to right z=10.5µm, 12.5µm. Scale bars (K): 10 µm. Scale bars (**R**): 50µm. Thickness, up to down z=2.8µm, 2.8µm.

### The developmental increase in phagocytosis efficiency is matched by increased microglia morphological complexity and surveillance

We then explored cellular mechanisms underlying the progressive increase in phagocytosis efficiency across the time course. Given that microglial phagocytosis is executed by *en passant* branches of their motile processes^24,40^, we assessed process morphology and dynamics across development. Sholl analysis of morphology from P2 to P60 revealed a progressive increase in process complexity, peaking at P28 in both hippocampus and cerebellum (**Figure 3H-J; Supp.** Figure 10A-C). To determine whether this structural maturation enhanced functional surveillance, we performed two-photon time-lapse imaging on acute hippocampal slices at P2 and P14 (**Figure 3K-M)**. While tip process velocity remained constant (**Figure 3L**), the cumulative area monitored by microglial processes increased significantly with age (**Figure 3M**). These results indicate that enhanced morphological complexity, rather than faster process dynamics, drives the developmental increase in surveillance, which likely contributes to more efficient apoptotic cell phagocytosis. The morpho-functional maturation of microglia was further supported by the upregulated expression of some homeostatic genes involved in environmental sensing, such as *P2ry12* and *Tmem119* (**Figure 3N**), in agreement with previous results^16,41,42^. Their increased expression suggests a progressive functional maturation of microglia toward active tissue surveillance.

Together, these findings indicate that microglial phagocytosis efficiency emerges gradually during postnatal development, in parallel with increasing transcriptional maturation and morphological complexity.

Finally, we analyzed which variables had a stronger effect on the phagocytosis efficiency by performing multiple correlations across the time course in the hippocampus (**Figure 3O**) and the cerebellum (**Supp. Figure 10D**). The Ph index correlated positively with the microglial density and morphological complexity (taking into account both process ramification and length), confirming the expected reliance of phagocytosis efficiency on the number of microglia and their surveillance capacity; and with the hippocampal volume, further supporting the maturation of microglial phagocytosis over postnatal development. However, the Ph index correlated negatively with the number of apoptotic cells, again suggesting that phagocytosis was not sufficiently engaged to cope with apoptosis through postnatal development. In addition, we observed a very strong negative correlation between microglial phagocytosis and proliferation (R^2^=0.87, *p*<0.0001) through the time course. In agreement, we found a very small percentage of microglia that were engaged both in proliferation (labeled with Ki67) and phagocytosis (displaying a pouch containing an apoptotic cell; **Figure 3P, Q; Supp.** Figure 10D-F), suggesting that microglia could not perform phagocytosis and proliferation simultaneously.

### Restricted overlap between proliferation and phagocytosis is conserved across different regions, conditions and species

Our findings support a sequential model of microglial maturation, in which proliferation is switched off and subsequently followed by morpho-functional specialization, including the acquisition of efficient phagocytosis. To further test whether proliferation and phagocytosis also occurred sequentially in other murine brain regions, we examined the somatosensory cortex, a region with perinatal development, from embryonic day E18 to postnatal day 14. In this region, the density of microglia reached a maximum at P10, whereas the percentage of proliferative microglia peaked at P4 and declined significantly from P10 onwards (**Figure 4A-C**), later than in the hippocampus and cerebellum. In contrast, apoptosis peaked at P4 and the Ph index gradually increased with age, reaching maximal efficiency by P10 (**Figure 4D, E**), suggesting the progressive acquisition of phagocytosis efficiency as in the hippocampus and cerebellum. While the cortex exhibited a greater temporal overlap between proliferation and phagocytosis than other regions, here the percentage of microglia simultaneously displaying proliferative and phagocytic phenotypes remained extremely low throughout development, reaching the maximum at P2 (0.9±0.4% of total microglia) (**Figure 4F, G**). Correlation analyses across all measured parameters further confirmed a significant inverse relationship between proliferation and phagocytosis (R²L=L0.37, *p*L=L0.005; **Figure 4H**), reinforcing the notion that these functional processes are temporally segregated during early postnatal maturation. Altogether, these results showed that the developmental milestones of microglia are maintained across regions but with region-specific temporal milestones, suggesting that specific regional cues might independently regulate the timing of the P/Q switch and functional maturation.

**Figure 4.**
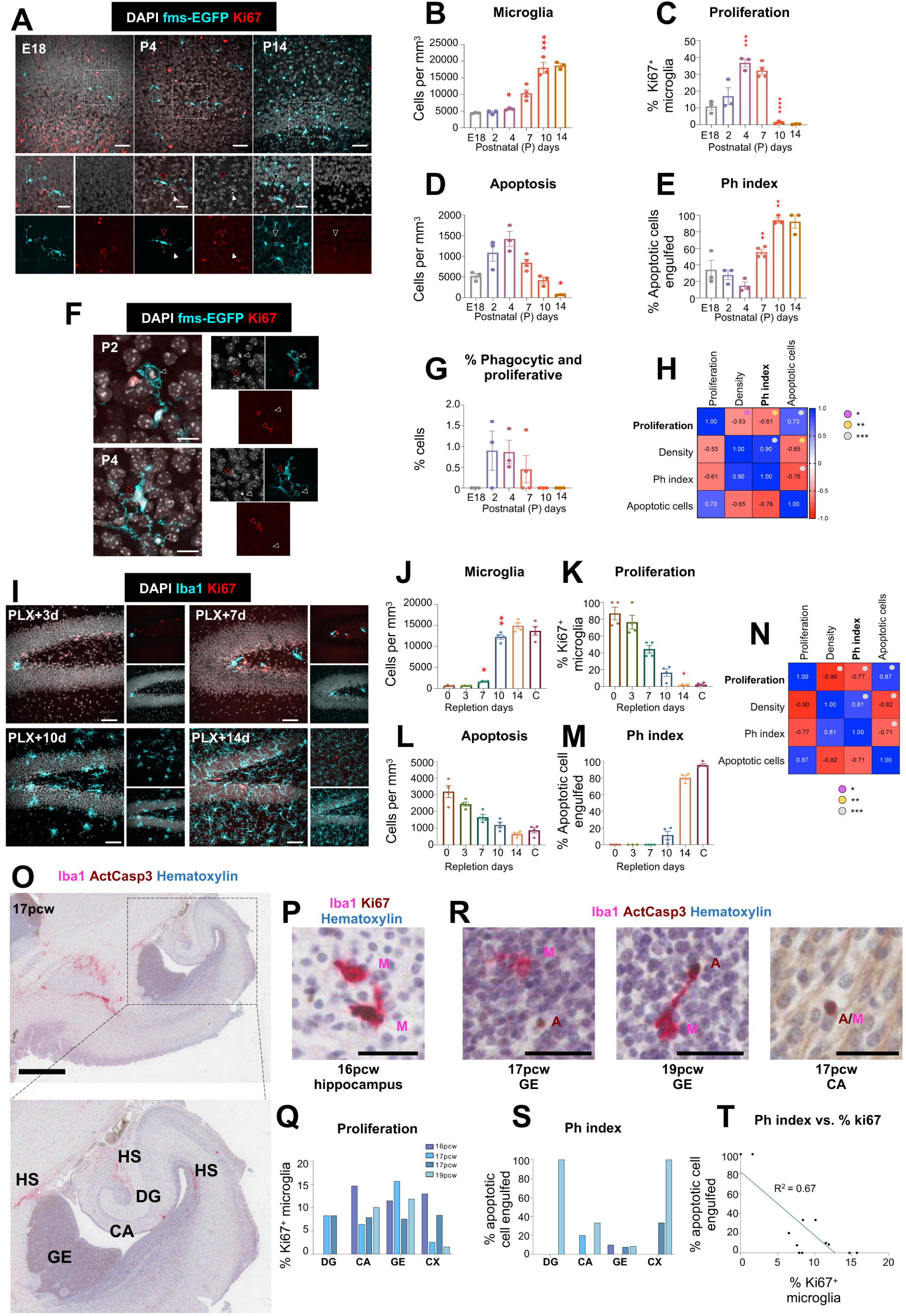
Microglial proliferation and phagocytosis correlate negatively across regions, conditions, and species. (**A**) Representative confocal images showing the somatosensory cortex at E18, P4, and P14 in fms-EGFP mice. Proliferation was identified by Ki67 expression (red), microglia as GFP positive cells (cyan), and nuclei with DAPI (white). White arrows indicate an apoptotic cell engulfed and red arrows indicate proliferative microglia. (**B**) Microglia density. (**C**) Percentage of microglia proliferation. (**D**) Density of apoptotic cells. (**E**) Phagocytic index (% of apoptotic cells phagocytosed by microglia). (**F**) High magnification representative confocal images of proliferative and phagocytic microglia at P2 and P4 in fms-EGFP mice. (**G**) Percentage of proliferative and phagocytic microglia over total microglia. (**H**) Multiple correlations in the somatosensory cortex showing the correlation coefficient R between proliferation, density, phagocytic index and density of apoptotic cells. **(I)** Representative confocal images showing the hippocampus 3, 7, 10, and 14 days after PLX3397 depletion. (**J**) Microglia density. (**K**) Percentage of microglia proliferation. (**L**) Density of apoptotic cells. (**M**) Phagocytic index (% of apoptotic cells phagocytosed by microglia). Bars show mean±SEM of n=4 mice. (**N**) Multiple correlations showing the correlation coefficient R between proliferation, density, phagocytic index and density of apoptotic cells. (**O**) Representative brightfield image of a section of a 17 pcw (post-conceptional week) human brain stained with Iba1 (magenta), activated caspase 3 (brown) and hematoxylin (blue), showing the hippocampal dentate gyrus (DG) and Cornu Ammonis (CA), the ganglionic eminence (GE), as well as two hot spots (HS) where microglia accumulate. Whole sections from the 4 specimens analyzed are shown in **Supp. Fig. 11A**. (**P**) Representative brightfield image of two proliferative microglia (M) in a 16pcw hippocampus, co-labeled with iba1 (magenta) and Ki67 (brown). (**Q**) Representative brightfield images of a non-phagocytosed apoptotic cell (A) with a nearby microglia (M) from a 17pcw GE (left panel), an apoptotic cell from a 19pcw GE (middle panel), and apoptotic microglia (A/M) from a 17pcw CA (right panel). Microglia are labeled with Iba1 (magenta), apoptotic cells with activated caspase 3 (brown), and nuclei with hematoxylin (blue). Bars show mean ± SEM of n=3 (B-H), n=4 (J-N) mice. Data were analyzed by one-way ANOVA followed by Tukey post hoc test, *p<0.05, **p<0.01, ***p<0.001. Significant correlations are shown by violet dots (p<0.05), tangerine dots (p<0.01), and gray dots (p<0.001). Only significant differences between consecutive ages are shown. Scale bars (**A**): upper row 50µm; bottom row 20µm. Thickness, left to right z=15.4µm, 16.8µm, 15.4µm; z=3.5µm, 6.3µm, 6.3µm. Scale bars (**F**): 50µm; Thickness z=2.8µm, 2.8µm. Scale bars (**I**): 50µm. Thickness, left to right A1 z= 22.4µm, 20.3µmm, 17.5µm, 20.3µm. Scale bar (**P**) 35μm. Scale bars (**R**): 30μm.

To determine whether this inverse relationship extended beyond developmental contexts, we investigated microglial dynamics in adult mice using a well-characterized pharmacological depletion-repopulation model with the colony-stimulating factor 1 receptor (CSF1R) inhibitor PLX3397 (PLX), fed in the diet for 14 days. After PLX removal, the density of microglia started to rise by 7 days, and the density recovered by 14 days (**Figure 4I, J**), while proliferative microglia dropped from 76.5%+8.2 at day 3 after PLX removal to 44.4%+4.6 at day 7 (**Figure 4K**). The phagocytosis efficiency did not significantly increase until day 14, as the number of apoptotic cells decreased over time (**Figure 4L, M**). Similar to the developmental pattern, microglial proliferation and phagocytosis remained inversely correlated during repopulation (R² = 0.59, *p* < 0.0001) (**Figure 4N**). Notably, no microglia were found to simultaneously engage in both proliferation and phagocytosis at any time point examined, reinforcing the idea that these functions represent temporally segregated cellular programs.

Finally, we tested the temporal relationship between proliferation and phagocytosis in the human developing brain. Microglia start to colonize the human brain from the 4^th^ postconceptional week (pcw), undergoing an initial embryonic wave of proliferation (5-8 pcw) followed by two fetal waves (9–12 pcw and 13–16 pcw) ^23^. The first recognizable rudiment of the human hippocampus with its subfields, neuroepithelium and infolding DG is around 14-16 pcw^43^. Therefore, we analyzed sections from 4 samples spanning 16-19 pcw (**Supp. Figure 11A, Supp. Table 1**). Due to the limited amount of microglia and apoptotic cells at these stages, we analyzed the hippocampus (DG and CA), the lateral telencephalic wall (immature temporal and parahippocampal cortices) and the lateral ganglionic eminence, where interneurons are produced (**Figure 4O**). We also observed hotspots of microglial accumulation close to the hippocampus with a high density of microglia (**Supp. Figure 11B, C**). Using color deconvolution in peroxidase/phosphatase-labeled sections to assess co-localization, we observed proliferative microglia (labeled with Iba1 and Ki67) across all regions and stages (**Figure 4P, Q**). To assess phagocytosis, we opted for a stringent definition of apoptotic cells (labeled with cleaved activated caspase 3) surrounded by a cytoplasmic pouch of microglia (labeled with Iba1; **Figure 4R**)^44^. We also observed few cases of apoptotic microglia (complete caspase3/Iba1 colocalization) but they were scarce. Overall, we determined that phagocytosis was more prevalent in later fetal stages (19pcw; **Figure 4S; Supp.** Figure 11D**, E**), leading to a significant negative correlation between microglial proliferation and phagocytosis (R^2^=0.672, *p*=0.001) (**Figure 4T**). Together, these findings demonstrate that the transition from a proliferative progenitor to a mature phagocyte constitutes a conserved and fundamental principle of microglial biology, operating across both development and repopulation contexts in mice and humans.

### Microglial functional maturation depends on intact proliferative dynamics

To directly assess the relationship between microglial proliferation and subsequent phagocytosis maturation during development, we examined whether disrupting proliferative signaling impaired microglial functional maturation. We pharmacologically inhibited CSF1R using the selective inhibitor GW2580, which has been described to prevent microglial proliferation without killing the cells^45^, administered once daily (50 mg/kg, intraperitoneally) over two early postnatal windows: P2–P5 and P2–P7. To assess recovery, a separate cohort was treated from P2–P5 and analyzed at P7. GW2580 treatment from P2–P5 significantly reduced hippocampal microglial proliferation and density (**Figure 5A-C**), accompanied by impaired maturation reflected in a decreased Ph index and reduced morphological complexity (**Figure 5D, E**; **Supp. Figure 12A, B**). Extending the treatment window to P2–P7 resulted in a similar, though non-significant, trend in density and proliferation reduction and impairments in phagocytosis efficiency and morphological complexity (**Figure 5B-E; Supp.** Figure 12A**, B**). Importantly, this effect of blocking CSF1R was transient, and microglial proliferation, phagocytosis efficiency and morphological complexity recovered control levels when assessed after cessation of treatment (P2–P5, analyzed at P7), (**Figure 5C-E Supp.** Figure 12A**, B).** However, a significant deficit in overall microglial density persisted (**Figure 5B**), indicating incomplete repopulation. The results in the cerebellum followed a similar pattern (**Supp. Figure 12C-I**). To assess whether the impairment of phagocytosis during development was primarily driven by reduced proliferation, we confirmed that CSF1R inhibition does not directly affect phagocytosis efficiency in adults (**Supp. Figure 12J-N**).

**Figure 5.**
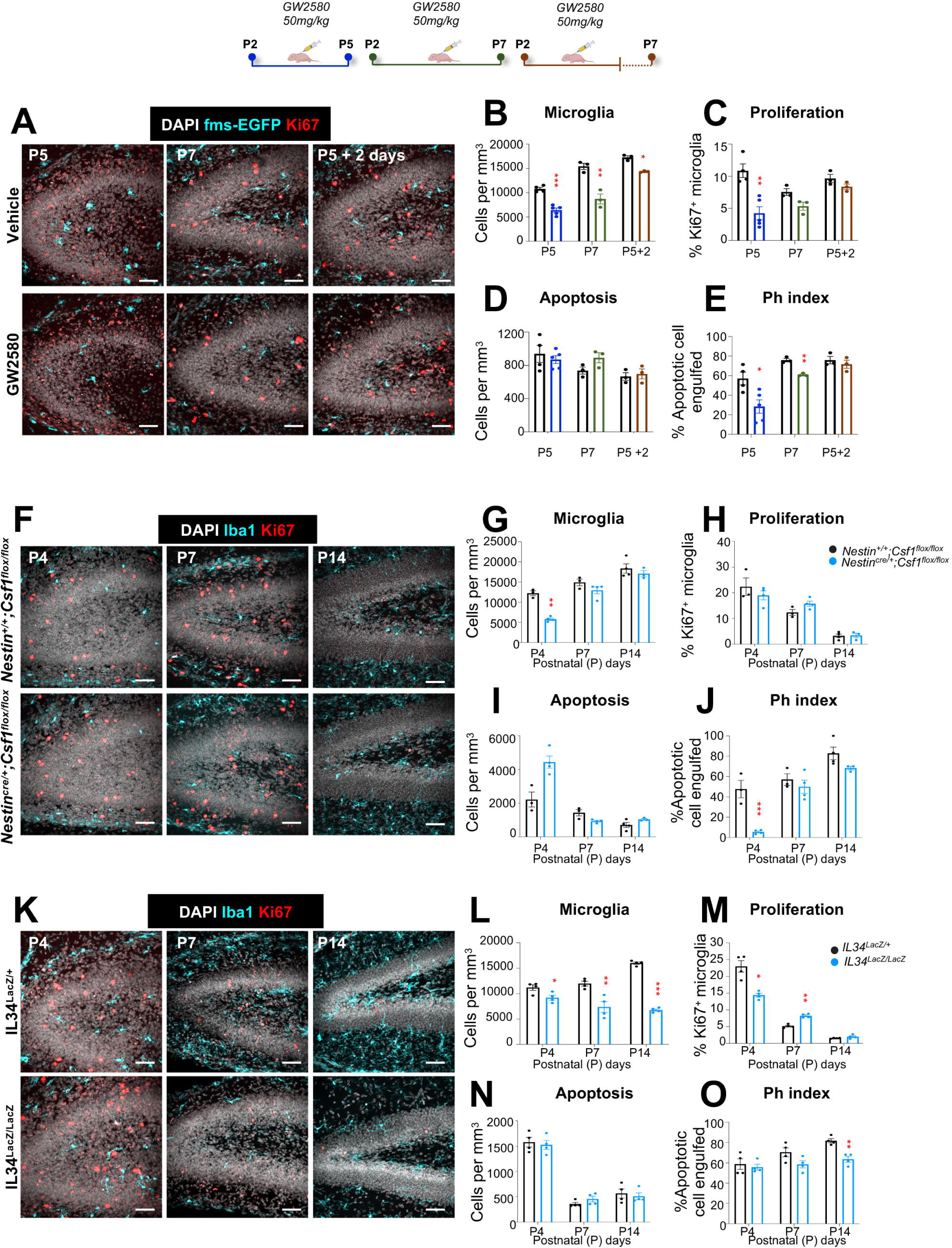
Pharmacological and transgenic inhibition of CSF1R signaling impairs proliferation and phagocytosis maturation. **(A)** Representative confocal images showing the hippocampus after intraperitoneal injection of GW2580 (50mg/kg) at P5, P7 and P7 after two days of recovery without injection in fms-EGFP mice. Proliferation was identified by Ki67 expression (red), microglia as Iba1 positive cells (cyan) and nuclei with DAPI (white). (**B**) Microglia density. (**C**) Percentage of microglia expressing Ki67. (**D**) Density of apoptotic cells. (**E**) Phagocytic index (% of apoptotic cells phagocytosed by microglia). Bars show mean±SEM of n=3 mice. (**F**) Representative confocal images showing the hippocampus at P4, P7 and P14 in control and mice with depletion of Csf1. (**G**) Microglia density. (**H**) Percentage of microglia proliferation. (**I**) Density of apoptotic cells. (**J**) Phagocytic index (% of apoptotic cells phagocytosed by microglia). (**K**) Representative confocal images showing the hippocampus at P4, P7 and P14 in control (Il34^LacZ/+^) and mice with ubiquitous depletion of Il34 (Il34^LacZ/LacZ^). (**L**) Microglia density. (**M**) Percentage of microglia proliferation. (**N**) Density of apoptotic cells. (**O**) Phagocytic index (% of apoptotic cells phagocytosed by microglia). (**R**) Multiple correlations showing the correlation coefficient R between proliferation, density, phagocytic index and density of apoptotic cells. Bars show mean±SEM of n=3 (B-F), n=3 control and n=4 Csf1 KO (H-L), n=4 (N-R). Data were analyzed via Two-way ANOVA followed by Tukey post hoc test, *p<0.05, **p<0.01, ***p<0.001. Significant correlations are shown by violet dots (p<0.05), tangerine dots (p<0.019, and gray dots p<0.001). Scale bars (A): 50µm. Thickness, left to right (upper panel) z=17.5µm, 18.2µm (middle panel) z=12.25µm, 11.2µm, (bottom panel), z=22.4µm, 21.7µm, 18.2µm. Experimental design drawing created with BioRender.com. Scale bars (G): 50µm. Thickness, left to right (upper panel) z=22.4µm, 18.2µm,17.5µm, (bottom panel) 18.2µm, 20.3 µm, 22.4µm. Scale bars (M): 50µm. Thickness, left to right (upper panel) z=21.7µm, 20.3µm, 17.5µm; (bottom panel) z=12.25µm, 11.2µm, 10.15µm.

We then confirmed these results in mice deficient for the CSF1R ligands *Csf1* and *Il34*. Given that hippocampal microglia primarily depend on *Il34*^46,47^*, Csf1* deficiency only resulted in a transient reduction in microglial density and phagocytosis at P4, with no major defects at later stages (P14), apart from minor alterations in morphological complexity (**Figure 5F-J; Supp.** Figure 13A**, B**). Conversely, in the cerebellum, where microglia are *Csf1*-dependent^47^, *Csf1* deficiency led to sustained reduction in microglial density, phagocytosis, and morphological complexity from P4 to P14 (**Supp. Figure 13C-I**). In contrast, *Il34* deficiency selectively impacted the hippocampus, where reduced proliferation at P4 resulted in persistent impairments in microglial density, phagocytosis, and morphology by P14 (**Figure 5K–O; Supp.** Figure 14A**, I**). Overall, these results reinforce that incomplete colonization due to proliferation deficits during this critical period compromises the ability of microglia to reach a fully mature, functional state.

### H3K4me3 dynamics coordinate the proliferation to maturation transition in developing microglia

We next explored the molecular mechanisms underlying the P/Q switch and the developmental maturation of microglia. First, we investigated whether the P/Q switch could be mediated by CSF1R and its ligands, IL-34 and CSF1, and analyzed whether changes in their expression patterns could underlie this switch. By RTqPCR, we found that *Il34* expression increased significantly over time in the hippocampus, and *Csf1* in the cerebellum, while *Csf1r* was upregulated in both regions (**Supp. Figure 15A**), in agreement with previous reports^47–49^. Thus, while our previous results (**Figure 5**) confirm the role of the CSF1R pathway in controlling microglial proliferation and/or survival, the increased expression of the pathway receptor and ligands over postnatal development contrasts with the synchronized drop of microglial proliferation at the P/Q switch at P3-P4.

To investigate alternative regulatory mechanisms underlying microglial development, we focused on epigenetic changes, which have been previously reported to follow sequential changes from embryonic to adult stages^15^, and performed chromatin immunoprecipitation followed by sequencing (ChIP-seq) targeting the active promoter mark H3K4me3^50^. Microglia were isolated from the hippocampus of fms-EGFP mice at P2 (proliferative stage), P7 (immediately post-switch stage), and P14 (mitotically quiescent stage) by FACS-sorting Pu.1+ nuclei following meningeal removal to minimize contamination by border-associated macrophages (**Supp. Figure 15B-D**). Flow cytometry confirmed that over 80% of Pu.1+ cells were microglia (GFP+, P2Y12+), validating sample purity (**Supp. Figure 15E, F**). Chromatin fragmentation (∼300 bp) and peak distribution (<1 kb from transcription start site) confirmed the technical quality of the ChIP-seq data (**Supp. Figure 15G-I**). Principal component analysis (PCA) revealed age-dependent shifts in the epigenetic landscape, with PC1 accounting for 29% of the variance and clearly separating samples by developmental stage (**Supp. Figure 15J, K**). Gene enrichment analysis of the top fifty genes contributing to each component indicated that PC1 was associated with developmental and maturation processes, while PC2 lacked significant biological enrichment, likely reflecting technical variability (**Supp. Figure 15L**).

Differential analysis identified 15 significantly regulated DNA regions (adjusted p < 0.1), with 13 regions showing decreased methylation and two exhibiting increased methylation over time (**Figure 6A, B**). Among the demethylated genes, we identified a cluster involved in cell proliferation, including *Ybx3*, *Mydgf*, *Iqgap3*, *Tmcc3*, and *Kif2c*, all of which are associated with cell survival, cytoskeletal regulation, or mitotic machinery^51–55^. These genes showed reduced promoter activity with age, consistent with a developmental decline in microglial proliferative capacity. A second set of demethylated genes, *Irf5*, *Ctsb*, and *Apoe*, are canonical markers of disease-associated microglia (DAM) and are typically upregulated in neurodegenerative contexts^56^. Their early postnatal demethylation suggests they may play roles in both development and neurodegenerative disease responses, pointing to a potential inverse development-aging regulation axis. A third category included genes involved in environmental sensing and migration: *P2ry1*, an ATP/ADP receptor potentially involved in chemotaxis, and *Camk2b*, a calcium signaling effector implicated in motility and differentiation^57,58^. The methylated genes *Lncppara* and *Fscn1* are associated with regulation of lipid metabolism and cytoskeletal remodeling during microglial maturation^59,60^. To capture finer and broader biological processes, we performed gene set enrichment analysis (GSEA), which revealed a predominant enrichment in pathways related to morphogenesis and differentiation (**Figure 6C**). These findings underscore a coordinated epigenetic program that silences proliferation-related genes while activating those involved in structural and functional maturation.

**Figure 6.**
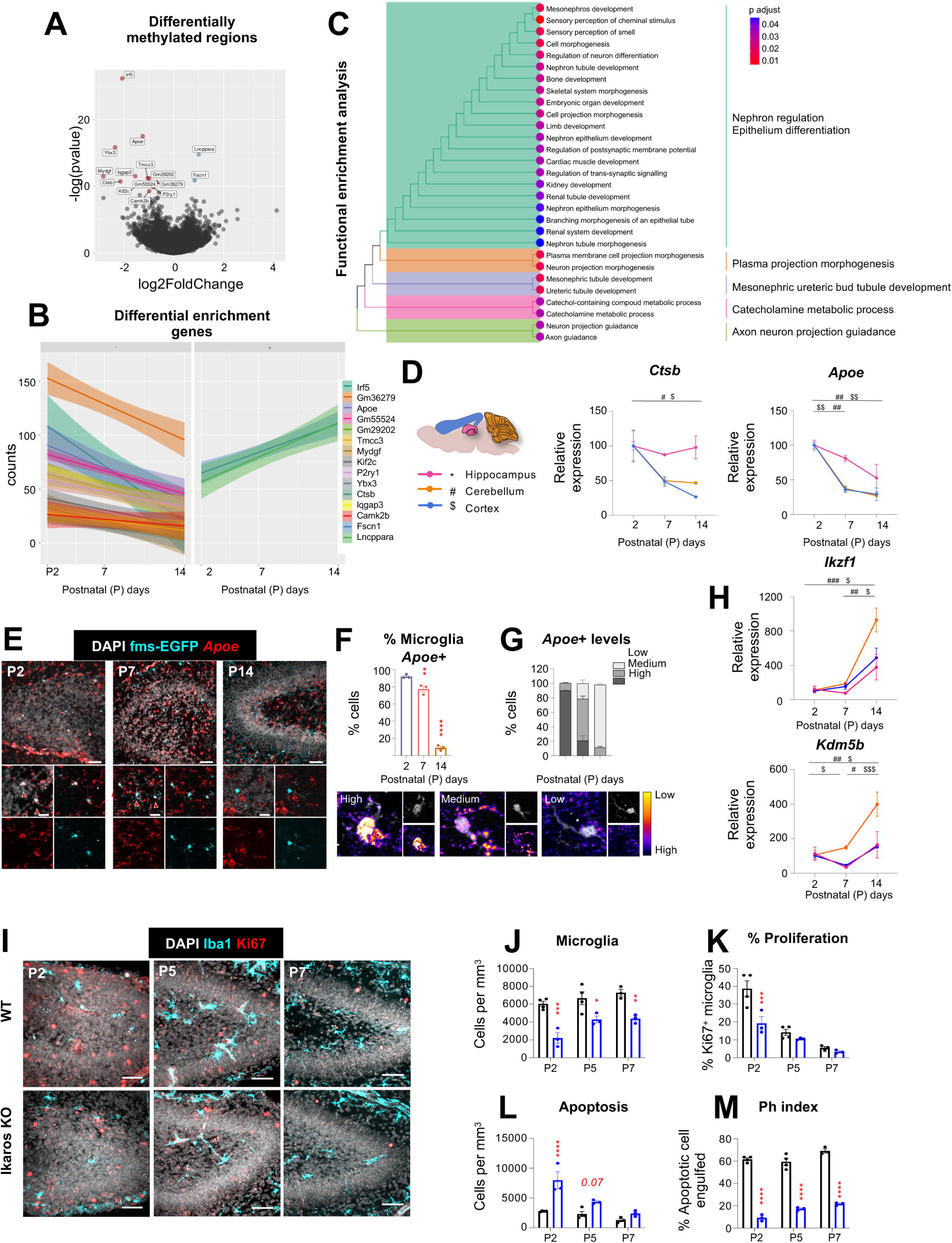
The demethylase inhibitor Ikaros controls the transition from proliferative progenitors to efficient phagocytes. (**A**) Volcano plot showing fifty differentially methylated regions. (**B**) Graph count of the fifteen differentially methylated regions. (**C**) Gene set enrichment analysis of genes with bottom fold changes. (**D**) Relative expression levels of *Ctsb* and *Apoe* mRNA at P2, P7, and P14 in the hippocampus, cerebellum and cortex of fms-EGFP mice (**E**) Representative confocal images showing the hippocampus, at P2, P7 and P14 in fms-EGFP mice. Microglia identified as GFP positive cells (cyan) and *Apoe* mRNA by red. High magnifications images show representative microglia from each age with *Apoe* expression (red arrow).(**F**) Percentage of microglia *Apoe*+ in the hippocampus. (**G**) *Apoe*+ microglia levels in the hippocampus. (**H**) Relative expression levels of *Ikzf1* and *KDM5B* mRNA at P2, P7 and P14 in the hippocampus, cerebellum and cortex of fms-EGFP mice. (**I**) Representative confocal images showing the hippocampus, at P2, P7 and P14 in control mice and mice with depletion of *Ikzf1*. (**J**) Microglia density. (**K**) Percentage of microglia proliferation. (**L**) Density of apoptotic cells. (K) Phagocytic index (% of apoptotic cells phagocytosed by microglia). Bars show mean ± SEM of n=3 (D, F-H), n=4 control and n=3 Ikaros KO (J-K). Data were analyzed by one-way ANOVA followed by Tukey post hoc test (D-H) or two-way ANOVA followed by Šídák’s multiple comparisons test (J-M). *p<0.05, **p<0.01, ***p<0.001. Only significant differences between consecutive ages are shown. Scale bars (I): 50µm. Thickness, left to right (upper panel) z=16.1µm, 16.1µm (middle panel) z=14.7µm, 14.0µm, (bottom panel), z=18.9µm, 17.5µm.

To determine whether changes in H3K4me3 promoter methylation correlated with gene transcription, we quantified the expression levels of key target genes at postnatal days P2, P7, and P14 in the hippocampus, cerebellum, and cortex. This analysis revealed consistent temporal and regional expression dynamics for genes such as *Ctsb* and *Apoe (***Figure 6D; Supp.** Figure 16A), which was further validated by RNAscope, showing a significant decreased in percentage of microglia positive for *Apoe* mRNA from P2 to P14 not only in the hippocampus, but also in the cerebellum and cortex (**Figure 6E, F; Supp.** Figure 16B-F). Additionally, microglia at P2 presented high levels of *Apoe* mRNA compared with positive microglia at P14, when expression levels were predominantly low (**Figure 6G; Supp.** Figure 16B-F). To extend these findings, we assessed gene expression patterns in five publicly available RNA-sequencing databases^15,16,61^. Three mouse datasets covering embryonic to postnatal stages consistently supported the age-dependent regulation of eight genes identified in our ChIP-seq analysis (*Apoe*, *Ybx3*, *Iqgap3*, *Ctsb*, *Kif2c*, *Camk2b*, *Gm29202*, and *Fscn1*) (**Supp. Figure 16G**). An exception was *Tmcc3*, which showed an increase in expression with age in two datasets, contrary to our findings (**Supp. Figure 16H**). Two additional human datasets covering prenatal and postnatal development ^62,63^ further corroborated the expression trends of several genes, including *Apoe*, *Mydgf*, and *Kif2c* **(Supp.** Figure 16H). Collectively, these findings highlight a largely conserved transcriptional trajectory across mice and humans for key epigenetically regulated genes during the microglial P/Q switch and functional maturation.

### The demethylase regulator Ikaros controls microglial proliferation and functional maturation

To directly test the role of epigenetic regulation in microglial development, we next focused on the modulation of H3K4 methylation. This process is primarily governed by two major enzyme families: the KMT2 methyltransferases, which catalyze the addition of methyl groups to H3K4, and the KDM5 demethylases, which remove these marks^64^. Importantly, both enzyme families are subject to regulatory control themselves. One of the most prominent regulators of demethylase activity is the Ikaros family of transcription factors. Among them, Ikzf1/Ikaros is a negative regulator of the demethylase KDM5B and controls ApoE methylation in B lymphocytes^65^. In the adult brain, Ikaros is exclusively expressed in microglia and its deletion results in abnormal morphology and inflammatory responses^66^. We observed that the expression of both *Ikaros* and *Kdm5b* mRNA increased from P2 to P14 in the hippocampus, cerebellum, and cortex (**Figure 6H**), consistent with a putative role of these factors in controlling microglia developmental maturation. In agreement, mice deficient in Ikaros showed reduced microglial density and proliferation (**Figure 6I-K**), and reduced phagocytosis accompanied by increased apoptosis from P2 to P7 (**Figure 6L, M**). Although this constitutive mouse model did not allow us to discriminate between direct effects of Ikzf1 on proliferation and/or phagocytosis, these results indicate that Ikaros is a key regulator of microglial functional maturation and identify a novel epigenetic mechanism orchestrating microglial development.

## DISCUSSION

In this paper, we have developed a novel data-driven mathematical model of brain colonization by microglia that allowed us to identify a critical milestone of their development: the synchronized switch between proliferating precursors to mitotically quiescent cells (P/Q switch; **Figure 7**). After the proliferation program is turned off, microglia mature morphologically and functionally to become efficient phagocytes. This sequence is conserved across brain regions, in depletion/repopulation paradigms, and occurs in the fetal human brain, pointing towards a conserved developmental trajectory. We then used experimental manipulation of the CSF1R pathway to demonstrate that proliferation-driven colonization is necessary for the microglial population to reach efficiency. However, the pattern of expression of CSF1R and its ligands CSF and IL34 across development did not match the timing of the P/Q switch, suggesting additional mediators. We focused on epigenetic mechanisms and found that the P/Q switch paralleled changes in histone methylation in genes involved in proliferation and disease-associated signatures, such as *Apoe*. Finally, we determined that the epigenetic modulator Ikaros controlled microglial proliferation and phagocytosis efficiency in the developing brain, illuminating a previously unidentified pathway of microglial maturation relevant to understanding the role of microglia in neurodevelopmental and neurodegenerative disorders.

**Figure 7.**
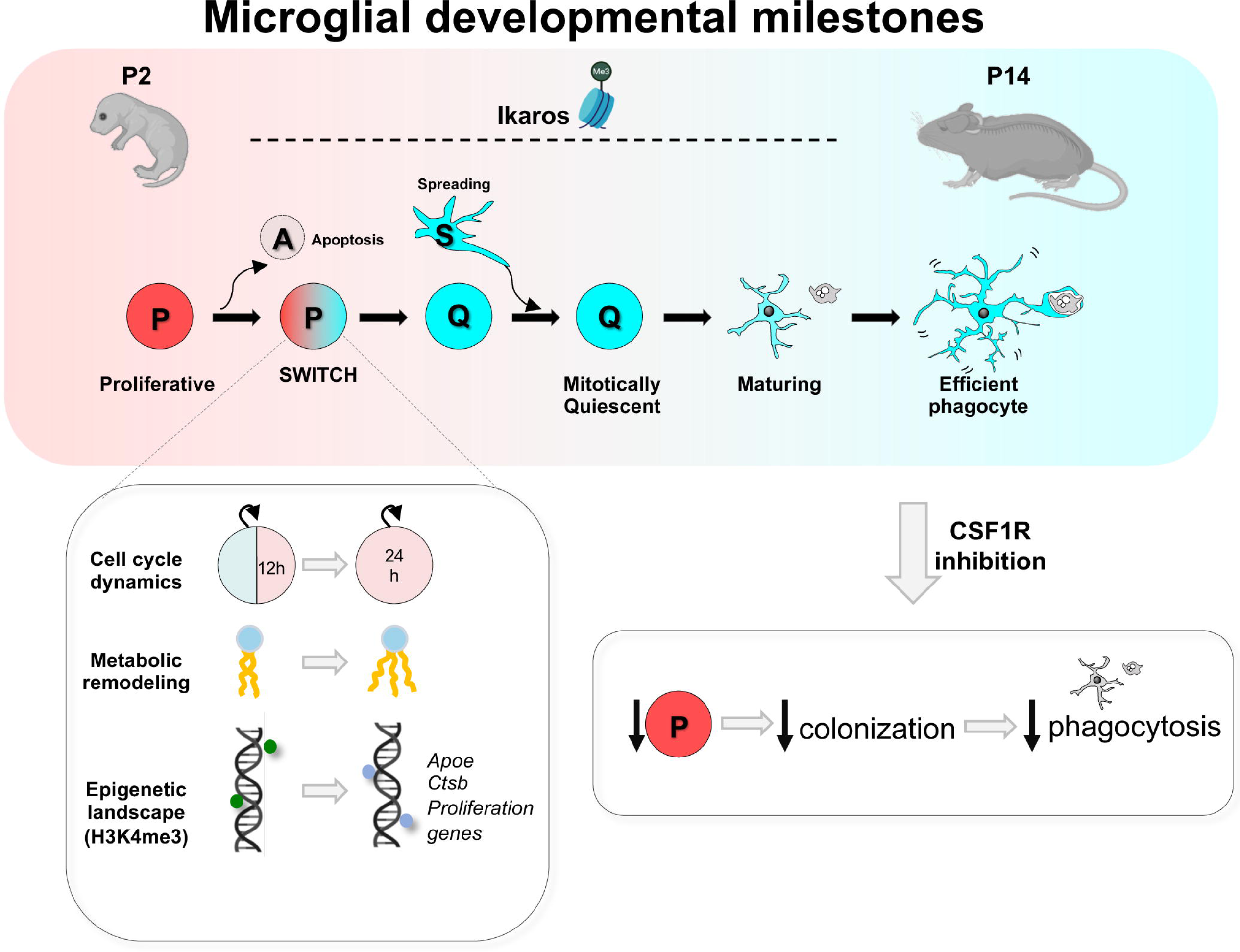
Microglia developmental milestones. During postnatal development, microglia undergo a conserved proliferative-to-quiescent (P/Q) transition around P3/P4, which precedes the acquisition of morphological complexity and efficient phagocytosis, and is accompanied by coordinated changes in cell-cycle dynamics, metabolic state, and the epigenetic landscape. Pharmacological or genetic disruption of CSF1R signaling during the proliferative phase impairs subsequent microglial complexity and phagocytic capacity, revealing an unexpected dependence of phagocytosis on proliferation-driven colonization. Stepwise developmental maturation of microglia is further associated with chromatin remodeling orchestrated by the epigenetic regulator Ikaros.

### Microglia synchronously switch off their proliferation program in early postnatal development

Our detailed longitudinal analysis of microglial colonization, powered by mathematical modeling, led us to identify the P/Q switch as a novel developmental milestone of microglia, as they synchronously turned off proliferation around postnatal day P3-P4. Local proliferation has long been recognized as a key event in microglial colonization in mice and humans^14,23^, and our model confirms that is indeed the major driving force of colonization in structures with postnatal development such as the hippocampus and cerebellum, with apoptosis and spreading between regions playing minor roles. While the two-population model fit microglial colonization data in both the hippocampus and cerebellum, it predicted opposite directions of spreading between sub-regions of the two structures (inwards and outwards, respectively), suggesting site-specific cues may govern this behavior. Overall, the synchronized phenotypic switch underscores the precision of microglial cell cycle control, consistent with observations that adult microglia in physiological conditions are predominantly quiescent^32^ and that microglial tumors are exceedingly rare^67^.

How this synchronization is orchestrated is a key question. The mathematical model predicted that proliferation was driven largely by endogenous factors, i.e., related to microglia themselves. In agreement, we demonstrated that the P/Q switch was reflected in a lengthening of the cell cycle and changes in the metabolic and lipidomic profile that demonstrated a transition from a metabolically diverse, proliferative state to a mitotically quiescent phenotype enriched in lipid metabolism. Furthermore, we identified novel epigenetic mechanisms operating on microglial development by showing that dynamic remodeling of H3K4me3 coordinates a developmental program that represses proliferation-associated genes while simultaneously activating transcriptional networks linked to structural and functional maturation. These results expand previous reports demonstrating that chromatin changes in enhancers are related to transitions in microglia development^15^. A similar epigenetic switch in maturation has also been described in other cell types like neurons^68^, suggesting a possible general developmental principle. We identified Ikaros as a critical regulator of microglial maturation, revealing a previously unrecognized role for this demethylase inhibitor in governing the transition from proliferative to functionally competent microglia. Of particular interest is the relationship between microglial proliferation and phagocytosis, Ikaros, and ApoE, a well-known Alzheimer risk gene^69^. Both Ikaros^66^ and ApoE^70^ expression are increased in brain tissue from Alzheimeŕs disease patients, leading to the intriguing possibility that this developmental pathway is reactivated in the diseased brain.

Our mathematical model also predicted that the transition from P to Q cells, i.e., switching off proliferation, was driven by factors related to the brain environment, specifically to the change in volume that occurs during postnatal development. It has been suggested that contact-inhibition mechanisms, which would be expected to be activated more as density increases, could be responsible for stopping microglial proliferation^71^. Our data agree with this hypothesis to some extent, as we found a negative correlation between proliferation and cell density. However, we also found that the P/Q switch occurred at P4, but the density peaked much later, at P14, and decreased up to P60, as the brain kept growing. Thus, it is possible that other mechanisms related to the brain environment operate to control the number of microglia. For instance, the P/Q switch is coetaneous with major developmental milestones such as the emergence of early sharp waves in the hippocampus^72^ and the loss of the proliferative potential of Purkinje neurons in the cerebellum^73^. Similarly, the drop in microglial proliferation at P7 coincides with thalamocortical innervation and intracortical circuit formation^74^, suggesting an uncharted role of neurons and/or neuronal activity in microglial developmental maturation. Conversely, these results also suggest that alterations in microglial development could affect the establishment of brain connectivity, representing a potential window of vulnerability in neurodevelopmental disorders.

### Phagocytosis efficiency emerges as a progressive feature of microglial maturation

In contrast to the prevailing view that microglia are inherently efficient phagocytes, we found that phagocytosis efficiency increased progressively during development, reaching maturity by P14. Early inefficiency was not due to saturation, as apoptotic load remained well below their clearance capacity in adults ^37^, but was related to individual cell factors, such as reduced surveillance and sensome gene expression in early time points; and to population factors, such as reduced microglial cell density. Thus, early microglia responded insufficiently to ethanol-induced apoptotic challenges, and only at P14 were they able to compensate the increase in apoptosis with an equivalent increase in phagocytosis. Reduced microglial phagocytosis efficiency in the developing brain resulted in longer apoptotic cell clearance time, compared to adult microglia^75^. Hence, the faster phagocytosis dynamics in adults highlights the progressive functional maturation of microglia, which emerges as a key functional hallmark of microglial development.

Whether immature microglia are also insufficiently responsive to other types of cargo remains to be determined. For example, thalamic microglia engulf synaptic inputs as early as P5^76^, and in the hippocampus synaptic engulfment has been shown to peak at P15^77^. Similarly, single cell transcriptomic studies identified a microglial signature called ATM (axonal tract microglia) that peaked around P4 and was engaged in myelin phagocytosis^16^. However, the different methods used in those studies prevent the assessment of phagocytosis efficiency, i.e., whether all the debris is actually removed, and this gap should be addressed in future studies.

### The sequence “proliferation to phagocytosis efficiency” is a conserved milestone of microglia maturation

A striking finding of our study is the conserved negative correlation between proliferation and phagocytosis in the hippocampus, cerebellum, and somatosensory cortex. Importantly, this principle was also observed in a depletion/repopulation paradigm in mice, and extended to human fetal brain tissue, where later developmental stages showed increased phagocytosis activity and reduced proliferation. Furthermore, at any given time point, region, model, and species, we found that few microglia were engaged in proliferation and phagocytosis simultaneously. This restricted overlap likely reflects a fundamental functional compromise, whereby active cell cycle engagement interferes with the execution of specialized functions^78^. Our pharmacological and genetic perturbations further demonstrated that intact proliferative dynamics and colonization are essential for acquiring efficient phagocytosis. This inverse relationship between proliferation and phagocytosis is consistent with findings from pathological contexts: in Alzheimer’s disease models, proliferating microglia near amyloid-β plaques fail to phagocytose deposits^79^, whereas CSF1R inhibition can drive microglia towards a state responsible for enhanced amyloid-β clearance^80^, reinforcing the notion that the coordination between these two processes is critical to brain damage responses.

We also speculate that the proliferation *vs* phagocytosis dilemma may be critical for generating immunocompetent human cerebral organoids (HCOs), a promising cellular model to generate accurate models that closely reflect human neurodevelopment. HCOs are generated from induced pluripotent stem cells (iPSCs) that are reprogrammed to generate neuroectodermal cells, such as neurons and astrocytes. Hence, conventional HCOs lack microglia, which are added using a variety of recently developed protocols that mimic the invasion and colonization that occur during brain development^81^. Thus, it is likely that human microglia added to the HCOs should follow the developmental milestones shown here, from proliferative progenitors to efficient phagocytes, in order to generate immunocompetent HCOs.

In summary, in this study we identified the milestones of microglial development during the postnatal period by integrating mathematical modeling with experimental validation. We discovered a conserved developmental trajectory from proliferation to efficient phagocytosis, driven by coordinated changes in cell cycle dynamics, metabolism, and epigenetic landscape, with critical implications for our understanding of the role of microglia in the diseased brain and for generating immunocompetent HCOs as avatars of the human brain.

## METHODS

### Mice

All experiments were performed in *fms*-EGFP (MacGreen) mice from postnatal day 2 (P2) to 2-month old, except where indicated, in which microglia constitutively express the green fluorescent reporter under the expression of the *fms* promoter ^27,82^. Analysis of microglia maturation was also performed in Ubi^+/+^;Reelin^flox/flox^ / Ubi^cre/+^; Reelin^flox/flox^ with C57BL/6 background mice^83^; IL34^LacZ/+^ / IL34^LacZ/LacZ^ with C57BL/6 background mice ^49^ (Greter et al. 2012);*Nestin^+/+^;Csf1^flox/flox^/ Nestin^cre/+^;Csf1^flox/flox^* with C57BL/6 background mice ^84,85^; Ikzf1 deficient mice ^86^and in C57BL/6 mice fed with PLX3397 diet for 15 days. In all experiments female and male mice were pooled. Mice were housed in 12:12 h light cycle with ad libitum access to food and water. Both males and females were used and pooled together. Some fms-EGFP mice were treated with an intraperitoneal injection of ethanol (2.5g/kg or 6mg/kg in saline solution). Another group of fms-EGFP mice was sequentially injected with EdU (Sigma-Aldrich) and BrdU (Sigma-Aldrich) (50mg/kg, each) at different time intervals. Another group of fms-EGFP mice was injected with GW2580 (50mg/kg, ip). Conditional Reelin deletion at early postnatal stages was achieved by administering tamoxifen (TMX; 1 mg/ml in 2.5% ethanol and 97.5% peanut oil) via daily intragastric injection (0.05 mg/pup) from P1 to P3. Mice were then sacrificed at P4, P7 and P14. All procedures followed the European Directive 2010/63/EU and were approved by the Ethics Committees of the University of the Basque Country EHU/UPV (Leioa, Spain; M20/2020/265 y M20/2022/020).

### Immunofluorescence

Immunostaining was performed following standard procedures ^87,88^. 50μm-thick sagittal free-floating brain vibratome sections were incubated in permeabilization solution (0.3% Triton-X100, 0.5% BSA in PBS; Sigma) for 2h at room temperature. Then, they were incubated overnight with the primary antibodies diluted in the permeabilization solution at 4 °C. After washing with PBS, sections were incubated with fluorochrome-conjugated secondary antibodies and DAPI (5Lmg/mL; Sigma) diluted in the permeabilization solution for 2 h at room temperature. After washing with PBS, tissue sections were mounted on glass slides with DakoCytomation Fluorescent Mounting Medium (Agilent).

For BrdU labeling an antigen retrieval procedure was performed prior to the fluorescent immunostaining by incubating in 2M HCl for 30min at 37°C, followed by washing with 0.1M sodium tetraborate for 10min at RT prior to the blockade of the sections. For reelin labeling an antigen retrieval procedure was performed prior to the fluorescent immunostaining by incubating in sodium citrate 10mM for 30 min at 90°C, followed by washing with PBS three times prior to the blockade of the sections.

For EdU detection, sections were incubated 30 minutes at RT with a label mix with 1% copper sulfate (CuSO_4_), 10% ascorbic acid and sulfo-Cy5-azide in PBS (all from Sigma), and then washed three times with PBS 0.3% Triton-X100.

| ANTIBODIES |  |  |  |  |
| --- | --- | --- | --- | --- |
| PRIMARY | HOST | DILUTION | SOURCE | CATALOGUE |
| GFP | Chicken | 1:1000 | Aves Lab | GFP-1020 |
| Iba1 | Guinea pig | 1:1000 | Synaptic systems | 234 308 |
| Ki67 | Rabbit | 1:750 | Abcam | ab16667 |
| P2Y12 | Rabbit | 1:1000 | AnaSpec | AS-55043A |
| Pu.1 | Rabbit | 1:400 | Cell Signaling | #2258 |
| BrdU | Mouse | 1:500 | Invitrogen | B35128 |
| Reelin | Mouse | 1:1000 | Abcam | Ab78540 |
| SECONDARY | HOST | DILUTION | SOURCE | CATALOGUE |
| AlexaFluor 488 anti-chicken | Donkey | 1:500 | Fisher | 17777517 |
| AlexaFluor RRX anti-rabbit | Donkey | 1:500 | Jackson Immunology Research | 711-295-152 |
| AlexaFluor 647 anti-rabbit | Donkey | 1:500 | Jackson Immunology Research | 711-605-152 |
| AlexaFluor RRX anti-guinea pig | Donkey | 1:500 | Jackson Immunology Research | 712-005-153 |
| DAPI |  | 1:1000 | Sigma-Aldrich | D9542 |

### RNAscope in situ hybridization

RNAscope was performed on 4% PFA treated fms-EGFP mice brains by conventional procedures. After 24h post-fixation in 4% PFA, brains were embedded in a sucrose gradient from 10, 20 to 30% until they sank, then frozen at −80°C in O.C.T. Compound (Cell Path). 15μm sections were cut onto Superfrost Plus slides using a LEICA CM1950 Cryostat (Leica) and kept at −80°C. RNAscope was carried out using the RNAscope Fluorescent Multiplex Kit (ACDBio) according to the manufacturer’s instructions with the catalog probe APOE (ACDBio, 313271) along with the provided positive control probe (UBC) and a negative probe targeting a bacterial gene. For the development of the fluorescent signal the fluorophore OPAL 570 dye (Akoya) was used. RNAscope was followed by immunofluorescence with anti-Iba1.

### Image analysis and quantification

All fluorescence immunostaining images were collected using a SP8 laser scanning microscope (Leica) with a 40x oil-immersion objective and a z-step of 0.7 μm. A 63X objective and a z-step of 0.3 μm was used for detailed images. At least 3 to 5 15-20 μm-thick z-stacks located at random positions containing the DG and CA were collected per hippocampal section, and a minimum of 6 sections per series were analyzed.

Quantitative analysis of microglia cell numbers and proliferation was performed using unbiased stereology methods as previously described ^88^. In brief, microglia cells were identified by GFP expression (in fms-EGFP mice) or Iba1, and proliferation by Ki67 nuclear staining. The number of microglia and proliferative microglia cells was quantified using ImageJ analysis software. The quantification per volume was estimated using the area of the region and thickness of the stack, followed by and estimation per hippocampus by measuring the volume of the hippocampus in Panoramic MIDI II Slide Scanner (3DHistech) pictures at 20x.

Cell death and phagocytosis analyses were carried out using unbiased stereology methods described in ^89^. Apoptotic cells were determined by their nuclear morphology visualized with DAPI. Phagocytosis was defined as the formation of a 3D pouch completely surrounding an apoptotic cell ^24^. In brain tissue sections, the number of apoptotic and microglia cells was estimated in the volume of the DG contained in the *z*-stack (determined by multiplying the thickness of the stack by the area of the DG at the center of the *z*-stack). To obtain the absolute numbers, that is, cells per septal hippocampus, the density values were multiplied by the volume of the septal hippocampus, which was calculated using ImageJ from images obtained with a 20× objective in Panoramic MIDI II Slide Scanner (3DHistech).

Microglia morphology analysis was semi-automatically performed using a built macro package for ImageJ (Cell_Shaper v5.1 https://www.achucarro.org/es/downloads/) based on the Sholl technique. The plugin calculates the number of intersections in each sphere shell for each cell, as well as the radius length of the last sphere that intersects a microglia ramification (termed here maximum length of microglia processes).

### Chromatin immunoprecipitation followed by sequencing

Microglia were isolated from mouse hippocampi. Dissections were mechanically disaggregated in cold conditions by a Dounce homogenizer followed by an automatic mixer at low potency. Nuclei extraction was performed in cold low-sucrose buffer (0.32M sucrose, 10mM HEPES pH 8, 5mM CaCl2, 3mM Mg(CH3COO)2, 0.1mM EDTA, 0.1% Triton-X-100, 1mM DTT), and tissue clogs were removed by filtering the cell suspension through a 40μm nylon strainer and myelin removed by using Percoll gradients. Following myelin removal, the cell pellet was resuspended in PBS and after three rounds of 5-minute centrifugation at 1000xg and resuspension in PBS, cells were incubated in GFP (1:1000) and Pu.1 (1:1000) for 30 minutes at RT, followed by incubation in goat anti-chicken 488 (1:1000) and goat anti-rabbit RXX (1:1000) for another 30 minutes at RT. After a 5-minute centrifugation at 1000xg, the pellet was resuspended in 500μl of sorting buffer (25 mM HEPES, 5 mM EDTA, 1% BSA, in HBSS). Microglia nuclei sorting was performed using a FACS Jazz instrument (BD Biosciences), in which the population of red-fluorescent cells was selected and collected in RIPA-SDS buffer (Diagenode) and stored at −80°C until processing.

Sorted Pu.1+ nuclei were lysed in RIPA-SDS buffer for 10 minutes at 4°C, was further sonicated using COVARIS M220 focused-ultrasonicator until reaching a mean size distribution of 300bp. Chromatin size was analyzed in a 1% agarose gel load. Chromatin was immunoprecipitated using True MicroChIP Kit (Diagenode) with H3K4me3 antibody (C15410003, Diagenode), and purified using MicroChIP Diapure Columns (Diagenode). Specificity of immunoprecipitation was assessed by real-time quantitative PCR prior to sequencing by NZYSpeedy qPCR Green Master Mix (NZYtech) with GAPDH and Myoglobin exon 2 primer pair (included in Diagenode True MicroChIP kit) and chromatin concentration assessed by Qubit Fluorometric quantification (ThermoFisher).

Chromatin immunoprecipitated (ChIP) DNA Libraries and sequencing were performed at the Genome Analysis Platform at CIC bioGUNE. DNA quality was assessed using Agilent 2100 Bioanalyzer (Agilent Technologies). Sequencing libraries were prepared using kit “*Microplex-library-prep-v3*” (Diagenode, Cat. # **C05010001**) and indexes “*Diagenode 24 Dual indexes for MicroPlex Kit v3*” (Diagenode, Cat. # **C05010003**), following Microplex-library-prep-v3 User Guide (**version 2 15_06_2021**). Library preparation protocol was started with 0.03-1.5 ng of ChIP DNA. Paired-end reading was performed with a read length of 100 bp and a sequencing depth of 40M reads using HiSeq200 sequencer following Illumina protocols. For sequencing data analysis, the nf-core/chipseq pipeline was followed^90^ specifying STAR^91^ as aligner and providing the Ensembl genome of *Mus musculus* (dna.primary_assembly ang gtf v109) as reference.

This pipeline carries out the initial filtering, alignment, normalization, peak calling, consensus peakset identification and count matrix creation. We imported the count matrix into R, following the analysis with DESeq2^92^, which provides methods for differential expression tests based on negative binomial generalized linear models. After initial filtering and exploration analysis (PCA), we applied the “Likelihood ratio test” LTR for hypothesis testing of timeline data (as suggested by DESeq2 vignettes). The differential expressed genes were selected based on the p-adjusted (< 0.1 Benjamini-Hochberg) value. Gene set enrichment analysis (GSEA) was carried out following clusterProfiler^93^ vignettes and ggplot2 was used for data visualization. The information for data download and processing is detailed in our GitHub page: https://github.com/rodrisenovilla/Pereira-Iglesias2025/tree/main

#### Metabolomics

##### Extraction

Metabolites form microglial cell pellets were extracted with 100 µL of cold methanol:acetonitrile:water (5:3:2). Samples were vortexed for 30 min at 4 °C and centrifuged (12,000 rcf, 10 min, 4 °C). Supernatants were collected and divided for metabolomics (40 µL) and lipidomics (60 µL) analysis, then dried using a SpeedVac. Metabolomics samples were reconstituted in 22 µL of 0.1% formic acid, and lipidomics samples were reconstituted in 22 µL of methanol prior to UHPLC–MS analysis.

##### Metabolomics

Samples were analyzed on a Thermo Vanquish UHPLC coupled to an Orbitrap Exploris 120 mass spectrometer. Separation was achieved using an Acquity UHPLC BEH C18 column (2.1 × 100 mm, 1.7 µm) with a 5-minute gradient in negative ion mode at 45 °C. The mobile phases consisted of 10 mM ammonium acetate in water (A) and 10 mM ammonium acetate in 50:50 acetonitrile:methanol (B). Data were converted to mzXML using RawConverter and processed with El-Maven (Elucidata) for peak integration and metabolite assignment. Pathway enrichment and statistical analyses were performed in MetaboAnalyst. Samples were injected in randomized order with pooled technical mixtures interspersed for quality control.

##### Lipidomics

Lipid extracts were analyzed using a Thermo Vanquish UHPLC coupled to a Q Exactive mass spectrometer. Separation was performed on a Kinetex C18 column (2.1 × 30 mm, 1.7 µm) using 10 µL injections and 5-minute gradients in both positive and negative ion modes at 50 °C. Mobile phase A consisted of 25:75 acetonitrile:water with 5 mM ammonium acetate, and mobile phase B consisted of 90:10 isopropanol:acetonitrile with 5 mM ammonium acetate. Data were processed using LipidSearch v5.0 (Thermo Fisher Scientific). Sample order was randomized, and pooled technical mixtures were injected regularly to monitor instrument performance.

#### qPCR

RNA from FACS-sorted microglia was isolated using a Total RNA isolation kit (NZYtech) and retrotranscribed using the Reverse Transcriptase kit (NZYtech) according to the manufacturer’s instructions. Primers were purchased from Thermo-Fisher (Table M3). Their specificity was assessed using melting curves. BiomarkTM HD technology was used in collaboration with the Genomic Service of the University of the Basque Country (UPV/EHU). Normalization was performed according to the NormFinder algorithm that selects the most stable reference genes ^94^.

RNA from whole brains was isolated by Total RNA isolation kit (NZYtech) and retrotranscribed using the Reverse Transcriptase kit (NZYtech) according to the manufacturer’s instructions. Real-time qPCR was performed following MIQE guidelines (Minimal Information for Publication of Quantitative Real Time Experiments) ^95^. Three biological replicas were run using the qPCR green master mix kit (NZYtech) according to the manufacturer’s instructions.

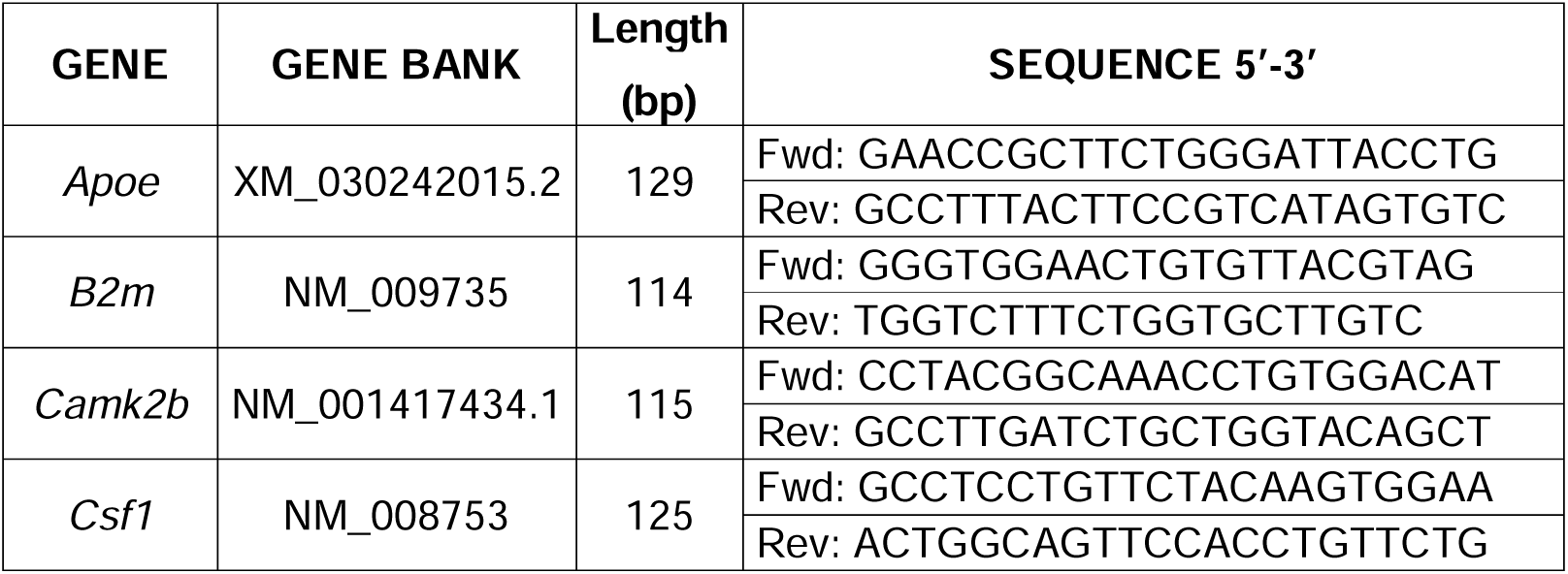
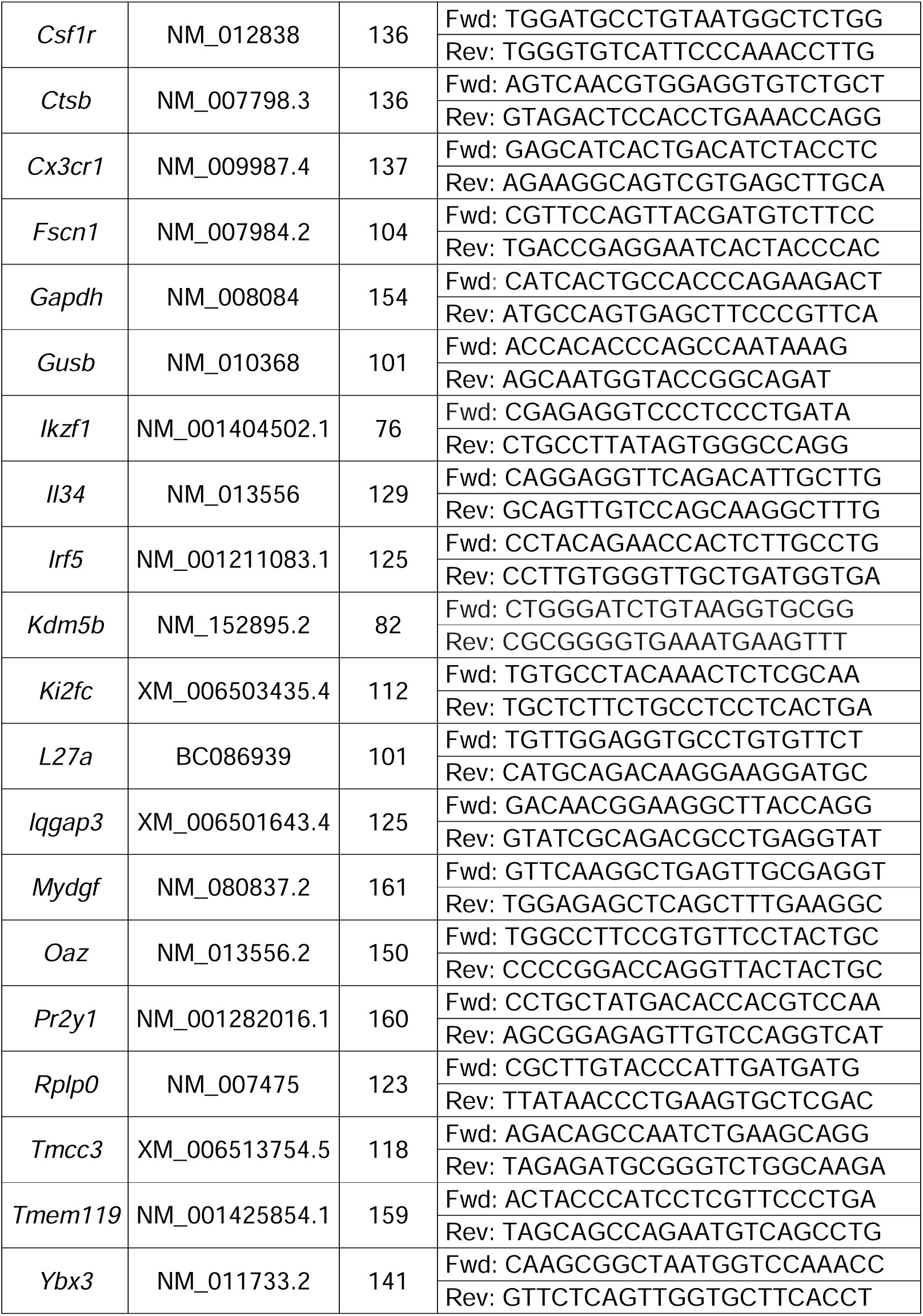

#### 2-photon imaging

P2 and P14 (±1 day) fms-eGFP mice were sacrificed by decapitation followed by brain dissection to obtain fresh 350 μm acute slices immersed in modified artificial cerebrospinal fluid (maCSF) (195 mM sucrose, 2.5 mM KCl, 1.25 mM NaH_2_PO_4_, 28 mM NaHCO_3_, 0.5 mM CaCl_2_, 1 mM L-ascorbic acid, 3 mM pyruvate, 7 mM glucose, 7 mM MgCl_2_ and 7 mM HEPES) at 4°C during slicing with a vibratome (Leica). Acute slices were then kept in artificial cerebrospinal fluid (aCSF) (118.93 mM NaCl, 2.5 mM KCl, 1 mM NaH_2_PO_4_, 25.95 mM NaHCO_3_, 1.58 mM MgCl_2_, 5 mM HEPES, 10 mM glucose, 2.5 mM CaCl_2_) with constant carbogen flux (95% O2/ 5% CO2). To image microglia, acute slices were placed in a superfusion system that perfused oxygenated aCSF. Recordings were obtained using a Femtomics two-photon microscope equipped with a 20X objective. A 40 μm stack with a 2 μm z-step was obtained every 2 minutes for a 10 minutes timecourse. Microglial process velocity analysis was performed using a built macro package for ImageJ (ProMolJ_Pack_v1.5 https://www.achucarro.org/es/downloads/) and the area surveilled was analyzed as the mean area per time frame.

### Human tissue analysis

#### Ethics statement and demographics of cases

The study was conducted according to the guidelines of the Declaration of Helsinki and approved by the ethics committees and the Oxford Brain Bank (Rec approval: 23/sc/0241, South Central Oxford C). We identified cases from human development from the 15^th^ −20^th^ pcw for this study. Immunohistochemistry steps were consistent with methods outlined below for tissue-processing for a total of 4 cases (3M:1F). All demographics are summarized in **Supp. Table 1**. All fetal scans and images are available upon request. Written informed consent was obtained from all mothers for use of these samples. Exclusion criteria were congenital abnormalities, genetic disorders, brain trauma, periventricular leukomalacia, and hypoxic ischaemic encephalopathy, infection, and brain trauma.

#### Anatomical areas

We focused on the forebrain along the temporal axis which includes the hippocampal formation, its neuroepithelium, the future temporal cortex, parahippocampal gyrus, and medial and lateral ganglionic eminences. The hippocampal formation is visible from the 14-16^th^ pcw^43^. Nissl and PAS-AB labelings were used for the delineation of the anatomical boundaries of these structures.

#### Immunohistochemistry and brightfield slide imaging

Paraffin-embedded blocks were cut into thin sections of 12 um on a microtome for immunohistochemistry. Brightfield immunohistochemistry experiments were performed using antibodies against microglia with the following dilutions: rabbit (019-19741, Wako, Cambridge Biosciences, UK) or mouse (ab283319, Abcam, UK), IBA1 at 1:1000; mouse anti-Ki67 (GA62661, Dako Omnis, Agilent, UK) at 1:400, and rabbit active cleaved caspase 3 (ab3623, Merck Millipore, UK) at 1:500. The first step was deparaffinization of formalin-fixed paraffin embedded sections in 100% xylene solution and rehydration in descending concentrations of diluted ethanol (100%, 96%, 90%, 70%). Antigen retrieval was done by heat induced epitope opening using citric acid buffer (pH = 6.2) for 30 min in a microwave. Thereafter, sections were pre-treated with dual enzyme block to block endogenous peroxidase and phosphatase activity. Sections were blocked with a solution of 5% Bovine serum albumin + Tween20 (0.1%) + normal horse serum (5%) in 1X PBS and then incubated with primary antibodies overnight. The next day, secondary antibodies were applied using either the Immunopress duet kit (MP7724 or MP7714, Vector labs, UK) with anti-mouse epitopes visualized with DAB chromogen in brown or magenta and anti-rabbit epitopes visualized with alkaline phosphatase in magenta or brown depending on the target. Sections were counterstained with haematoxylin or methyl green and coverslipped with permanent mounting medium before imaging. Imaging was done using high-resolution histological slide scanners: Aperio Imagescope (Oxford, UK) for analysis at 0.45 x 0.45 um pixel resolution and Olympus VS200 0.09 x 0.09 um for subcellular pixel resolution.

#### Statistical analysis

GraphPad Prism (San Diego, CA, USA) was used to perform the statistical analysis. Data were tested for normality and homoscedasticity. When the data did not comply with these assumptions, a logarithmic transformation or a square root was performed, and the data were analyzed using parametric tests. Two-sample experiments were analyzed by Students’ t-test and more than two sample experiments with one-way or two-way ANOVA. In case that homoscedasticity or normality was not achieved with a logarithmic transformation, data were analyzed using a Kruskal-Wallis ranks test, followed Dunn method as a posthoc test. Two sample non-parametric data were analyzed using Mann Whittney U test. Only p < 0.05 is reported to be significant. Multiple correlations were analyzed by Pearson correlation coefficients and represented with a heatmap of the R^2^ values.

#### Mathematical model description

We investigated three different types of mathematical models based on ordinary differential equations (ODEs) to account for the non-monotonic (increasing-then-decreasing) behavior of microglia densities over time within the cerebellum and hippocampus. Each mathematical model is given by a system of ODEs, where each differential equation tracks the change in microglia density within a specific layer of the cerebellum/hippocampus.

The first mathematical framework is a modified version of a classical logistic model for population growth in which cells increase exponentially to attain an equilibrium density (**Supp. Fig. 3AB**). In our case, we take the equilibrium density to be inversely proportional to the volume of the sub-region, hence it decreases over time for each sub-region of the cerebellum or hippocampus. We account for spreading between different sub-regions by assuming linear rates of transition between neighboring compartments. Apoptosis is similarly incorporated as a linear decay term in each ODE. Here, “linear” denotes that the mechanism occurs with a constant doubling- or halving-time.

The second mathematical model assumes linear rates for proliferation, apoptosis, and spreading (**Supp. Fig. 3CD**). To account for the non-monotonic behavior of experimental data, we further assume that each ODE has an additional positive time-dependent source, which represents the migration of microglia cells from outside the cerebellum/hippocampus into each respective sub-region. Each of these sources is assumed to decrease exponentially in time.

The final mathematical model assumes that microglia in each layer can be decomposed into two sub-populations (**Fig. 1H-M; Supp.** Fig. 3E-I). Both subtypes can proliferate, undergo apoptosis, transform to the other type, and spread to neighboring layers of the cerebellum or hippocampus. All mechanisms occur with linear rates that represent “natural” or “intrinsic” processes since they do not depend on other cell types. However, to produce non-monotonic solutions we assume that all subtype populations can also undergo these processes at “extrinsic” rates that are mediated by a time-dependent “environmental factor” that is assumed to be proportional to the volume of each sub-region. The ODEs for both sub-populations are identical, and only after parameter estimation do we assess their distinguishing features.

The systems of ODEs were solved in Python (version 3.10.12), using an implicit first-order backward differentiation formula from the SciPy library^96^. More details on the mathematical models and simulations can be found in the Supplementary Information.

#### Parameter estimation

We apply a Bayesian approach to parameter estimation of the mathematical models. As a first approximation, we take experimental observations to be independent and normally distributed with mean values equal to the mathematical model output. This assumption, which is common in mathematical biology, enables us to determine the likelihood in closed form. The prior parameter distribution for all parameters, including the initial cell densities and standard deviation of the normal distributions for the likelihood, is assumed to be uniformly distributed with a lower bound of zero. Posterior distributions are inferred using a Metropolis-Hastings MCMC (Markov chain Monte Carlo) sampler with adaptive proposal covariance. This is implemented in the parameter estimation toolbox PINTS^97^.

We applied the Akaike information criterion (AIC), Bayesian information criterion (BIC), and the Watanabe-Akaike information criterion (WAIC)^98,99^ to compare how well the three types of models fit the data (**Fig. 1J**). All three metrics are based on logarithmic scales and quantify the log-normalized likelihood of the model, penalizing by either the number of parameters (AIC), number of data points (BIC), or how badly the model is predicted to match new observations (WAIC). Lower scores indicate more accurate and parsimonious model fits to the data.

## Acknowledgements

This work was supported by grants from the Spanish Ministry of Science and Innovation Competitiveness MICIU/AEI/10.13039/501100011033 and by “ERDF A way of making Europe” (RTI2018-099267-B-I00, PID2022-136698OB-I00 and RYC-2013-12817 to AS), a Basque Government Department of Education project (PIBA 2020_1_0030; http://www.euskadi.eus/basque-government/department-education/), a Basque Government Consolidated Research Group (IT1473-22), and an Alzheimer Association award (AARG-NTF-24-1304352) to AS; an EPSRC grant no EP/R014604/1 to JAC; Spanish Ministry of Science and Innovation Competitiveness projects to ES/YM (PID2022-138105OB-C21) and AG (PID2024-157024OB-I00); grants from CNRS, CIRB and ANR (ANR-23-CE16-0001 and ANR-23-TERC-0017) to MST; grants from ISCIII (FORT23/00008), AEI (PID2022-136526OB-I00 & PID2024-157045OB-I00) and by the regional Government of Andalucia (Biotechnology Plan applied to Health PRTR--C17.I1) to AP, and grants a Springboard grant funded by the British Council (No 1170803491) and an NIH grant (4R01NS124848-02) to DAM.

MPI, MGD and SB are recipients of predoctoral fellowships from the Spanish Ministry of Science and Innovation. JMT is a recipient of a predoctoral fellowship from the Basque Government. RSG is recipient of Fundacion Tatiana predoctoral fellowship. BMR is the recipient of a Junta de Andalucia predoctoral fellowship. SV was supported by a French ministerial PhD fellowship. WDM and JAC were supported by the Advanced Grant Nonlocal-CPD (Nonlocal PDEs for Complex Particle Dynamics: Phase Transitions, Patterns and Synchronization) of the European Research Council Executive Agency (ERC) to JAC under the European Union’s Horizon 2020 research and innovation program (grant agreement No. 883363). WDM and JAC would also like to thank the Isaac Newton Institute for Mathematical Sciences, Cambridge, for support and hospitality during the program Mathematics of Movement, where work on this paper was undertaken. Achucarro and UPV/EHU SGIker technical and human support is gratefully acknowledged. We acknowledge the Oxford Brain Bank, supported by the Brains for Dementia Research (BDR) (Alzheimer Society and Alzheimer Research UK), the NIHR Oxford Biomedical Research Centre. Finally, WDM, CF and JAC would also like to thank the Basque Center of Applied Mathematics for hosting them during several stages of this research.

## REFERENCES

1. Sierra, A., Paolicelli, R. C. & Kettenmann, H. Cien Años de Microglía: Milestones in a Century of Microglial Research. Trends in Neurosciences vol. 42 Preprint at 10.1016/j.tins.2019.09.004 (2019).

2. Ginhoux, F. et al. Fate mapping analysis reveals that adult microglia derive from primitive macrophages. Science (1979) 330, (2010).

3. Kierdorf, K. et al. Microglia emerge from erythromyeloid precursors via Pu.1-and Irf8-dependent pathways. Nat Neurosci 16, (2013).

4. Hickman, S., Izzy, S., Sen, P., Morsett, L. & El Khoury, J. Microglia in neurodegeneration. Nature Neuroscience vol. 21 Preprint at 10.1038/s41593-018-0242-x (2018).

5. Cowan, M. & Petri, W. A. Microglia: Immune regulators of neurodevelopment. Frontiers in Immunology vol. 9 Preprint at 10.3389/fimmu.2018.02576 (2018).

6. Wu, S. et al. Il34-Csf1r Pathway Regulates the Migration and Colonization of Microglial Precursors. Dev Cell 46, (2018).

7. Renee, M. et al. CSF1 receptor signaling is necessary for microglia viability, which unmasks a cell that rapidly repopulates the microglia-depleted adult brain. Neuron 82, (2015).

8. Zusso, M. et al. Regulation of postnatal forebrain amoeboid microglial cell proliferation and development by the transcription factor runx1. Journal of Neuroscience 32, (2012).

9. Butovsky, O. et al. Identification of a unique TGF-β-dependent molecular and functional signature in microglia. Nat Neurosci 17, (2014).

10. Zöller, T. et al. Silencing of TGFβ signalling in microglia results in impaired homeostasis. Nat Commun 9, (2018).

11. Olson, M. C. et al. PU.1 is not essential for early myeloid gene expression but is required for terminal myeloid differentiation. Immunity 3, (1995).

12. Gosselin, D. et al. Environment drives selection and function of enhancers controlling tissue-specific macrophage identities. Cell 159, (2014).

13. Schulz, C. et al. A lineage of myeloid cells independent of myb and hematopoietic stem cells. Science (1979) 335, (2012).

14. Barry-Carroll, L. et al. Microglia colonize the developing brain by clonal expansion of highly proliferative progenitors, following allometric scaling. Cell Rep 42, (2023).

15. Matcovitch-Natan, O. et al. Microglia development follows a stepwise program to regulate brain homeostasis. Science (1979) 353, (2016).

16. Hammond, T. R. et al. Single-Cell RNA Sequencing of Microglia throughout the Mouse Lifespan and in the Injured Brain Reveals Complex Cell-State Changes. Immunity 50, (2019).

17. Li, Q. et al. Developmental Heterogeneity of Microglia and Brain Myeloid Cells Revealed by Deep Single-Cell RNA Sequencing. Neuron 101, (2019).

18. Hickman, S. E. et al. The microglial sensome revealed by direct RNA sequencing. Nat Neurosci 16, (2013).

19. Márquez-Ropero, M., Benito, E., Plaza-Zabala, A. & Sierra, A. Microglial Corpse Clearance: Lessons From Macrophages. Frontiers in Immunology vol. 11 Preprint at 10.3389/fimmu.2020.00506 (2020).

20. Utz, S. G. et al. Early Fate Defines Microglia and Non-parenchymal Brain Macrophage Development. Cell 181, (2020).

21. Morsch, M. et al. In vivo characterization of microglial engulfment of dying neurons in the zebrafish spinal cord. Front Cell Neurosci 9, (2015).

22. Cunningham, C. L., Martínez-Cerdeño, V. & Noctor, S. C. Microglia regulate the number of neural precursor cells in the developing cerebral cortex. Journal of Neuroscience 33, (2013).

23. Menassa, D. A. et al. The spatiotemporal dynamics of microglia across the human lifespan. Dev Cell 57, (2022).

24. Sierra, A. et al. Microglia shape adult hippocampal neurogenesis through apoptosis-coupled phagocytosis. Cell Stem Cell 7, (2010).

25. Diaz-Aparicio, I. et al. Microglia Actively Remodel Adult Hippocampal Neurogenesis through the Phagocytosis Secretome. J Neurosci 40, (2020).

26. Sierra, A., Abiega, O., Shahraz, A. & Neumann, H. Janus-faced microglia: Beneficial and detrimental consequences of microglial phagocytosis. Frontiers in Cellular Neuroscience Preprint at 10.3389/fncel.2013.00006 (2013).

27. Sasmono, R. T. et al. A macrophage colony-stimulating factor receptor-green fluorescent protein transgene is expressed throughout the mononuclear phagocyte system of the mouse. Blood 101, (2003).

28. Leto, K. et al. Consensus Paper: Cerebellar Development. Cerebellum vol. 15 Preprint at 10.1007/s12311-015-0724-2 (2016).

29. Schultz, C. & Engelhardt, M. Anatomy of the hippocampal formation. in The Hippocampus in Clinical Neuroscience vol. 34 (2014).

30. Tan, Y. L., Yuan, Y. & Tian, L. Microglial regional heterogeneity and its role in the brain. Molecular Psychiatry vol. 25 Preprint at 10.1038/s41380-019-0609-8 (2020).

31. Kee, N., Sivalingam, S., Boonstra, R. & Wojtowicz, J. M. The utility of Ki-67 and BrdU as proliferative markers of adult neurogenesis. J Neurosci Methods 115, (2002).

32. Askew, K. et al. Coupled Proliferation and Apoptosis Maintain the Rapid Turnover of Microglia in the Adult Brain. Cell Rep 18, (2017).

33. Frotscher, M., Haas, C. A. & Förster, E. Reelin controls granule cell migration in the dentate gyrus by acting on the radial glial scaffold. Cerebral Cortex 13, (2003).

34. Jossin, Y. Reelin functions, mechanisms of action and signaling pathways during brain development and maturation. Biomolecules vol. 10 Preprint at 10.3390/BIOM10060964 (2020).

35. Harris, L., Zalucki, O. & Piper, M. BrdU/EdU dual labeling to determine the cell-cycle dynamics of defined cellular subpopulations. J Mol Histol 49, (2018).

36. Calegari, F., Haubensak, W., Haffher, C. & Huttner, W. B. Selective lengthening of the cell cycle in the neurogenic subpopulation of neural progenitor cells during mouse brain development. Journal of Neuroscience 25, (2005).

37. Marquez-Ropero, M. et al. Metabolic and transcriptional adaptations to phagocytosis sustain microglia functionality and regenerative properties. Preprint at 10.1101/2025.09.18.676800 (2025).

38. Konishi, H. et al. Astrocytic phagocytosis is a compensatory mechanism for microglial dysfunction. EMBO J 39, (2020).

39. Abiega, O. et al. Neuronal Hyperactivity Disturbs ATP Microgradients, Impairs Microglial Motility, and Reduces Phagocytic Receptor Expression Triggering Apoptosis/Microglial Phagocytosis Uncoupling. PLoS Biol 14, (2016).

40. Kamei, R. & Okabe, S. In vivo imaging of the phagocytic dynamics underlying efficient clearance of adult-born hippocampal granule cells by ramified microglia. Glia 71, (2023).

41. Cho, S. H. et al. CX3CR1 protein signaling modulates microglial activation and protects against plaque-independent cognitive deficits in a mouse model of Alzheimer disease. Journal of Biological Chemistry 286, (2011).

42. Haynes, S. E. et al. The P2Y12 receptor regulates microglial activation by extracellular nucleotides. Nat Neurosci 9, (2006).

43. Kier, E. L., Kim, J. H., Fulbright, R. K. & Bronen, R. A. Embryology of the human fetal hippocampus: Mr imaging, anatomy, and histology. American Journal of Neuroradiology 18, (1997).

44. Perochon, T. et al. Unraveling microglial spatial organization in the developing human brain with DeepCellMap, a deep learning approach coupled with spatial statistics. Nature Communications 16, (2025).

45. Neal, M. L. et al. Pharmacological inhibition of CSF1R by GW2580 reduces microglial proliferation and is protective against neuroinflammation and dopaminergic neurodegeneration. FASEB Journal 34, (2020).

46. Wang, Y. et al. IL-34 is a tissue-restricted ligand of CSF1R required for the development of Langerhans cells and microglia. Nat Immunol 13, (2012).

47. Easley-Neal, C., Foreman, O., Sharma, N., Zarrin, A. A. & Weimer, R. M. CSF1R Ligands IL-34 and CSF1 Are Differentially Required for Microglia Development and Maintenance in White and Gray Matter Brain Regions. Front Immunol 10, (2019).

48. Wei, S. et al. Functional overlap but differential expression of CSF-1 and IL-34 in their CSF-1 receptor-mediated regulation of myeloid cells. J Leukoc Biol 88, (2010).

49. Greter, M. et al. Stroma-Derived Interleukin-34 Controls the Development and Maintenance of Langerhans Cells and the Maintenance of Microglia. Immunity 37, (2012).

50. Beacon, T. H. et al. The dynamic broad epigenetic (H3K4me3, H3K27ac) domain as a mark of essential genes. Clinical Epigenetics vol. 13 Preprint at 10.1186/s13148-021-01126-1 (2021).

51. Coles, L. S., Diamond, P., Occhiodoro, F., Vadas, M. A. & Shannon, M. F. Cold shock domain proteins repress transcription from the GM-CSF promoter. Nucleic Acids Res 24, (1996).

52. Große-Segerath, L. et al. Identification of myeloid-derived growth factor as a mechanically-induced, growth-promoting angiocrine signal for human hepatocytes. Nat Commun 15, (2024).

53. Leone, M. et al. IQGAP3, a YAP target, is required for proper cell-cycle progression and genome stability. Molecular Cancer Research 19, (2021).

54. Wang, Y. H. et al. Transmembrane and coiled-coil domain family 3 (TMCC3) regulates breast cancer stem cell and AKT activation. Oncogene 40, (2021).

55. Bakhoum, S. F., Thompson, S. L., Manning, A. L. & Compton, D. A. Genome stability is ensured by temporal control of kinetochore-microtubule dynamics. Nat Cell Biol 11, (2009).

56. Deczkowska, A. et al. Disease-Associated Microglia: A Universal Immune Sensor of Neurodegeneration. Cell vol. 173 Preprint at 10.1016/j.cell.2018.05.003 (2018).

57. Léon, C. et al. The P2Y1 receptor is an ADP receptor antagonized by ATP and expressed in platelets and megakaryoblastic cells. FEBS Lett 403, (1997).

58. Proietti Onori, M. & van Woerden, G. M. Role of calcium/calmodulin-dependent kinase 2 in neurodevelopmental disorders. Brain Res Bull 171, (2021).

59. Montaigne, D., Butruille, L. & Staels, B. PPAR control of metabolism and cardiovascular functions. Nature Reviews Cardiology vol. 18 Preprint at 10.1038/s41569-021-00569-6 (2021).

60. Jansen, S. et al. Mechanism of actin filament bundling by fascin. Journal of Biological Chemistry 286, (2011).

61. Hochgerner, H., Zeisel, A., Lönnerberg, P. & Linnarsson, S. Conserved properties of dentate gyrus neurogenesis across postnatal development revealed by single-cell RNA sequencing. Nat Neurosci 21, (2018).

62. Han, C. Z. et al. Human microglia maturation is underpinned by specific gene regulatory networks. Immunity 56, (2023).

63. Yaqubi, M. et al. Analysis of the microglia transcriptome across the human lifespan using single cell RNA sequencing. J Neuroinflammation 20, (2023).

64. Soares, L. M. et al. Determinants of Histone H3K4 Methylation Patterns. Mol Cell 68, (2017).

65. Schwickert, T. A. et al. Ikaros prevents autoimmunity by controlling anergy and Toll-like receptor signaling in B cells. Nat Immunol 20, (2019).

66. Ballasch, I. et al. Ikzf1 as a novel regulator of microglial homeostasis in inflammation and neurodegeneration. Brain Behav Immun 109, (2023).

67. Mathews, A. et al. Microglioma in a child - A further case in support of the microglioma entity and distinction from histiocytic sarcoma. Clin Neuropathol 35, (2016).

68. Ciceri, G. et al. An epigenetic barrier sets the timing of human neuronal maturation. Nature 626, (2024).

69. Raulin, A. C. et al. ApoE in Alzheimer’s disease: pathophysiology and therapeutic strategies. Molecular Neurodegeneration vol. 17 Preprint at 10.1186/s13024-022-00574-4 (2022).

70. Yamada, T., Kondo, A., Takamatsu, J. ich, Tateishi, J. & Goto, I. Apolipoprotein E mRNA in the brains of patients with Alzheimer’s disease. J Neurol Sci 129, (1995).

71. Barry-Carroll, L. & Gomez-Nicola, D. The molecular determinants of microglial developmental dynamics. Nat Rev Neurosci 25, 414–427 (2024).

72. Cossart, R. & Khazipov, R. HOW DEVELOPMENT SCULPTS HIPPOCAMPAL CIRCUITS AND FUNCTION. Physiol Rev 102, (2022).

73. Joyner, A. L. & Bayin, N. S. Cerebellum lineage allocation, morphogenesis and repair: impact of interplay amongst cells. Development (Cambridge) vol. 149 Preprint at 10.1242/dev.185587 (2022).

74. Nwabudike, I. & Che, A. Early-life maturation of the somatosensory cortex: sensory experience and beyond. Front Neural Circuits 18, 1–8 (2024).

75. Kamei, R. & Okabe, S. In vivo imaging of the phagocytic dynamics underlying efficient clearance of adult-born hippocampal granule cells by ramified microglia. Glia 71, (2023).

76. Schafer, D. P. et al. Microglia Sculpt Postnatal Neural Circuits in an Activity and Complement-Dependent Manner. Neuron 74, (2012).

77. Paolicelli, R. C. et al. Synaptic pruning by microglia is necessary for normal brain development. Science (1979) 333, (2011).

78. Perea-Resa, C., Bury, L., Cheeseman, I. M. & Blower, M. D. Cohesin Removal Reprograms Gene Expression upon Mitotic Entry. Mol Cell 78, (2020).

79. Villacampa, N. et al. Proliferating microglia exhibit unique transcriptomal and functional alterations in Alzheimer’s disease. Alzheimers Dement 17, (2021).

80. Piccioni, G. et al. Switch to phagocytic microglia by CSFR1 inhibition drives amyloid-beta clearance from glutamatergic terminals rescuing LTP in acute hippocampal slices. Transl Psychiatry 14, (2024).

81. Cuesta-Puente, X., et al. Building Immunocompetent Cerebral Organoids From a Developmental Perspective. GLIA Preprint at 10.1002/glia.70062 (2025).

82. Sierra, A., Gottfried-Blackmore, A. C., Mcewen, B. S. & Bulloch, K. Microglia derived from aging mice exhibit an altered inflammatory profile. Glia 55, (2007).

83. Pardo, M. et al. Adult-specific Reelin expression alters striatal neuronal organization: implications for neuropsychiatric disorders. Front Cell Neurosci 17, (2023).

84. Tronche, F. et al. Disruption of the glucocorticoid receptor gene in the nervous system results in reduced anxiety. Nat Genet 23, (1999).

85. Harris, S. E. et al. Meox2Cre-mediated disruption of CSF-1 leads to osteopetrosis and osteocyte defects. Bone 50, (2012).

86. Wang, J. H. et al. Selective defects in the development of the fetal and adult lymphoid system in mice with an Ikaros null mutation. Immunity 5, (1996).

87. Abiega, O. et al. Neuronal Hyperactivity Disturbs ATP Microgradients, Impairs Microglial Motility, and Reduces Phagocytic Receptor Expression Triggering Apoptosis/Microglial Phagocytosis Uncoupling. PLoS Biol 14, (2016).

88. Beccari, S., Diaz-Aparicio, I. & Sierra, A. Quantifying Microglial Phagocytosis of Apoptotic Cells in the Brain in Health and Disease. Curr Protoc Immunol 122, (2018).

89. Beccari, S., Diaz-Aparicio, I. & Sierra, A. Quantifying Microglial Phagocytosis of Apoptotic Cells in the Brain in Health and Disease. Curr Protoc Immunol 122, (2018).

90. Ewels, P. A. et al. The nf-core framework for community-curated bioinformatics pipelines. Nature Biotechnology vol. 38 Preprint at 10.1038/s41587-020-0439-x (2020).

91. Dobin, A. et al. STAR: Ultrafast universal RNA-seq aligner. Bioinformatics 29, (2013).

92. Love, M. I., Huber, W. & Anders, S. Differential gene expression analysis based on the negative binomial distribution. Genome Biol 15, (2014).

93. Wu, T. et al. clusterProfiler 4.0: A universal enrichment tool for interpreting omics data. Innovation 2, (2021).

94. Andersen, C. L., Jensen, J. L. & Ørntoft, T. F. Normalization of real-time quantitative reverse transcription-PCR data: A model-based variance estimation approach to identify genes suited for normalization, applied to bladder and colon cancer data sets. Cancer Res 64, (2004).

95. Bustin, S. A. et al. The MIQE guidelines: Minimum information for publication of quantitative real-time PCR experiments. Clin Chem 55, (2009).

96. Virtanen, P. et al. SciPy 1.0: fundamental algorithms for scientific computing in Python. Nat Methods 17, (2020).

97. Clerx, M. et al. Probabilistic Inference on Noisy Time Series (PINTS). J Open Res Softw 7, (2019).

98. Cavanaugh, J. E. & Neath, A. A. The Akaike information criterion: Background, derivation, properties, application, interpretation, and refinements. Wiley Interdisciplinary Reviews: Computational Statistics vol. 11 Preprint at 10.1002/wics.1460 (2019).

99. Gelman, A., Hwang, J. & Vehtari, A. Understanding predictive information criteria for Bayesian models. Stat Comput 24, (2014).

