## Supplementary text for "Epigenetic control of microglial developmental milestones from proliferative progenitors to efficient phagocytes"

Includes supplementary figures and legends, supplementary Table 1, and supplementary mathematical model.

### SUPPLEMENTARY FIGURE LEGENDS

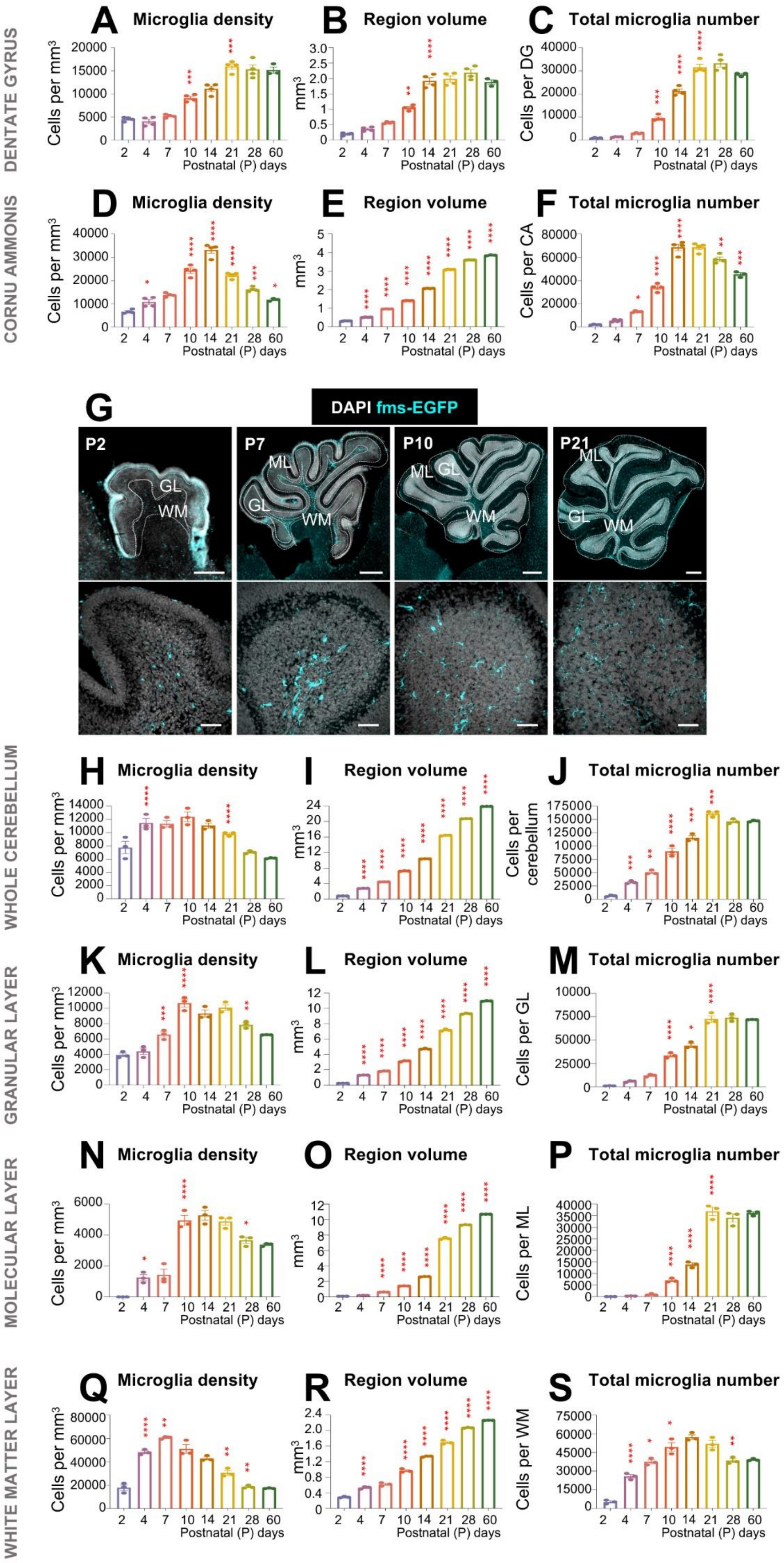
DENTATE GYRUS
A Microglia density
B Region volume
C Total microglia number
CORNU AMMONIS
D Microglia density
E Region volume
F Total microglia number
G
DAPI fms-EGFP
P2
P7
P10
P21
ML
GL
WM
WHOLE CEREBELLUM
H Microglia density
I Region volume
J Total microglia number
GRANULAR LAYER
K Microglia density
L Region volume
M Total microglia number
MOLECULAR LAYER
N Microglia density
O Region volume
P Total microglia number
WHITE MATTER LAYER
Q Microglia density
R Region volume
S Total microglia number
Cells per mm³
mm³
Cells per DG
Cells per CA
Cells per cerebellum
Cells per GL
Cells per ML
Cells per WM
Postnatal (P) days

**Supplementary Figure 1. Microglia density across hippocampal and cerebellar subregions during postnatal development.** (**A**) Density of microglia in DG. (**B**) DG region volume. (**C**) Total microglia number per DG volume. (**D**) Density of microglia in CA. (**E**) CA region volume. (**F**) Total microglia number per CA volume. Bars show mean±SEM of n=4 mice. (**G**) Representative slide scanner and confocal images showing the cerebellum at P2, P7, P10, and P21 in fms-EGFP mice. DAPI (white) staining shows nuclei and GFP (cyan) microglia. (**H**) Density of microglia in the cerebellum. (**I**) Cerebellum region volume. (**J**) Total microglia numbers per cerebellar volume. (**K**) Density of microglia in GL. (**L**) GL region volume. (**M**) Total microglia numbers per GL volume. (**N**) Density of microglia in ML. (**O**) ML region volume. (**P**) Total microglia number per ML volume. (**Q**) Density of microglia in WM. (**R**) WM region volume. (**S)** Total microglia number per WM volume.

Bars show mean ± SEM of n=3 mice (A-F; H-S). Data were analyzed by one-way ANOVA followed by Tukey post hoc test, *p<0.05, **p<0.01, ***p<0.001. Only significant differences between consecutive ages are shown. Scale bars (**G**): upper panel 500µm; bottom panel 50µm. Thickness, left to right bottom row z=13.3µm, 17.5µm 18.9µm, 16.8µm.

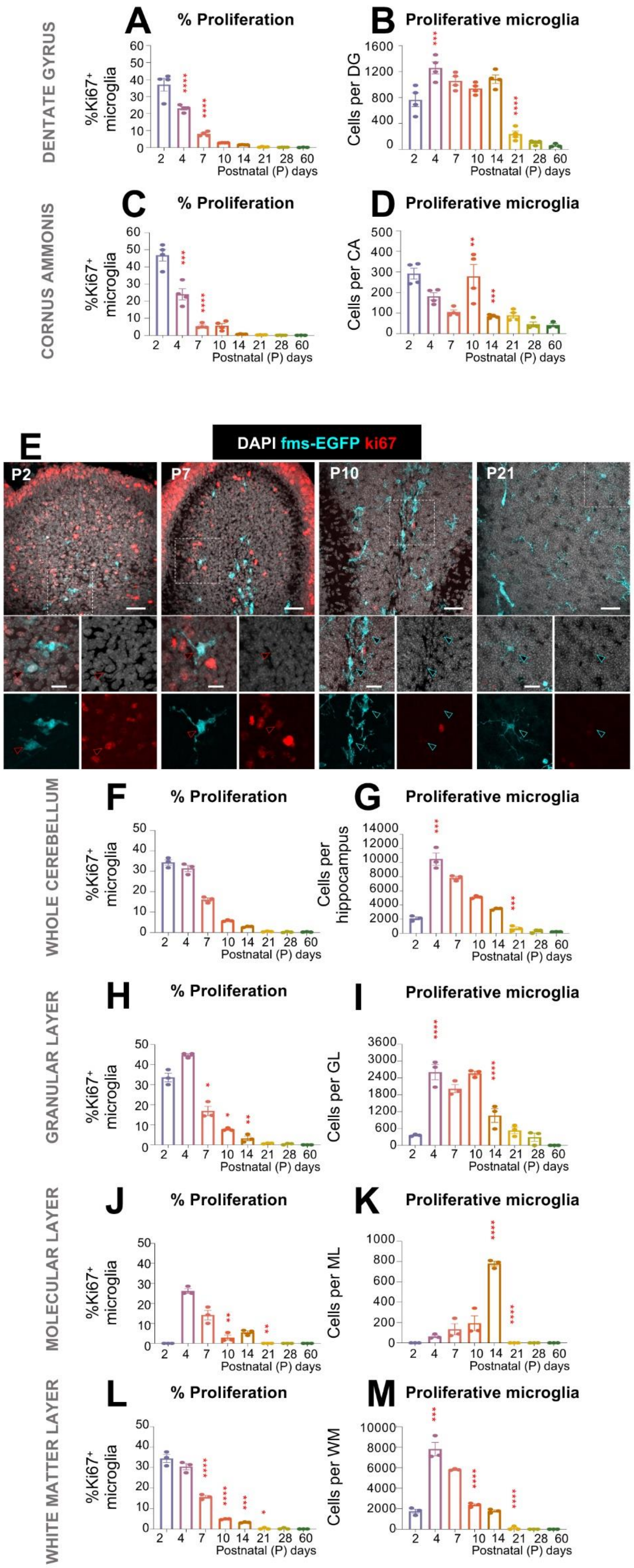
DENTATE GYRUS
A % Proliferation
%Ki67+ microglia
Postnatal (P) days
B Proliferative microglia
Cells per DG
CORNUS AMMONIS
C % Proliferation
D Proliferative microglia
Cells per CA
E
DAPI fms-EGFP ki67
P2
P7
P10
P21
WHOLE CEREBELLUM
F % Proliferation
G Proliferative microglia
Cells per hippocampus
GRANULAR LAYER
H % Proliferation
I Proliferative microglia
Cells per GL
MOLECULAR LAYER
J % Proliferation
K Proliferative microglia
Cells per ML
WHITE MATTER LAYER
L % Proliferation
M Proliferative microglia
Cells per WM

**Supplementary Figure 2. Microglia proliferation across hippocampal and cerebellar subregions during postnatal development**. (**A**) Percentage of proliferative microglia in DG. (**B**) Total number of proliferative microglia in DG. (**C**) Percentage of proliferative microglia in CA. (**D**) Total number of proliferative microglia in CA. (**E**) Representative confocal images showing the cerebellum at P2, P7, P10, and P21 in fms-EGFP mice. Proliferation was identified by Ki67 expression (red), microglia as GFP positive cells (cyan) and nuclei with DAPI (white). (**F**) Percentage of proliferative microglia in the cerebellum. (**G**) Total number of proliferative microglia in the cerebellum. (**H**) Percentage of proliferative microglia in GL. (**I**) Total number of proliferative microglia in GL. (**J**) Percentage of proliferative microglia in ML. (**K**) Total number of proliferative microglia in ML. (**L**) Percentage of proliferative microglia in WM. (**M**) Total number of proliferative microglia in WM. Bars show mean ± SEM of n=4 (A-D) mice. Bars show mean ± SEM of n=3 (F-M) mice. Data was analyzed by one-way ANOVA followed by Tukey post hoc test, *$p<0.05$, **$p<0.01$, ***$p<0.001$. Only significant differences between consecutive ages are shown.

Scale bars: upper panel 50µm; bottom panel 20µm. Thickness, left to right upper panel z=6.3µm, 7.7µm, 7.7µm, 10.5µm; bottom panel z=2.1µm, 2.1µm, 2.1µm, 2.1µm.

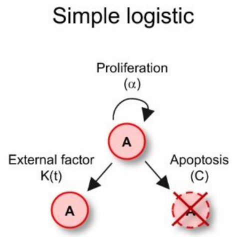


**A** $$\frac{dW}{dt} = \underbrace{\alpha W}_{\text{proliferation}} \cdot \underbrace{\left(1-\frac{W}{K_0 e^{-\gamma t}}\right)}_{\text{fraction of cells surviving competition}} - \underbrace{\beta W,}_{\text{migration/apoptosis}}$$

**B**

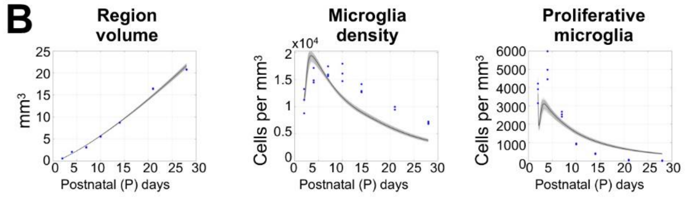


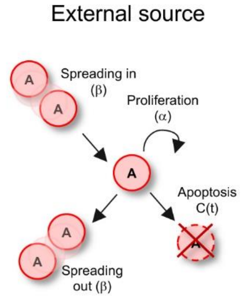


**C** $$\frac{\mathrm{d}W}{\mathrm{d}t} = \underbrace{\alpha W}_{\text{proliferation}} - \underbrace{\beta W}_{\text{apoptosis/migration out}} + \underbrace{S(t),}_{\text{influx from outside the white matter}}$$

**D**

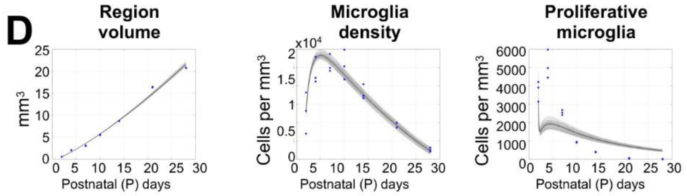


**E**

$$\frac{\mathrm{d}P_w}{\mathrm{d}t} = (\underbrace{\alpha_{w,1}}_{\text{intrinsic prol.}} - \underbrace{\alpha_{w,2}}_{\text{intrinsic diff.}} - \underbrace{\beta_{w,1}}_{\text{migration}} - \underbrace{c_{w,1}}_{\text{apoptosis}})P_w + (\underbrace{k_{w,1}}_{\text{extrinsic prol.}} - \underbrace{k_{w,2}}_{\text{extrinsic diff.}})E_w P_w,$$

$$\frac{\mathrm{d}Q_w}{\mathrm{d}t} = \alpha_{w,2}P_w + (\underbrace{\alpha_{w,3}}_{\text{intrinsic prol.}} - \underbrace{\beta_{w,2}}_{\text{migration}} - \underbrace{c_{w,2}}_{\text{apoptosis}})Q_w + (k_{w,2})P_w E_w,$$

$$\frac{\mathrm{d}K_w}{\mathrm{d}t} = \underbrace{\alpha_{w,1}P_w + \alpha_{w,3}Q_w + k_{w,1}E_w P_w}_{\text{increase due to cell proliferation}} - \underbrace{\delta_w K_w,}_{\text{decrease in Ki67}^+\text{ expression}}$$

**F**

Cerebellum

| Model | AIC | BIC | DIC | WAIC |
|---|---|---|---|---|
| Logistic | -125.93 | -31.91 | -161.73 | -162.92 |
| Linear | -142.84 | -39.11 | -183.64 | -181.60 |
| **Two population** | **-271.70** | **-109.61** | **-358.43** | **-357.46** |

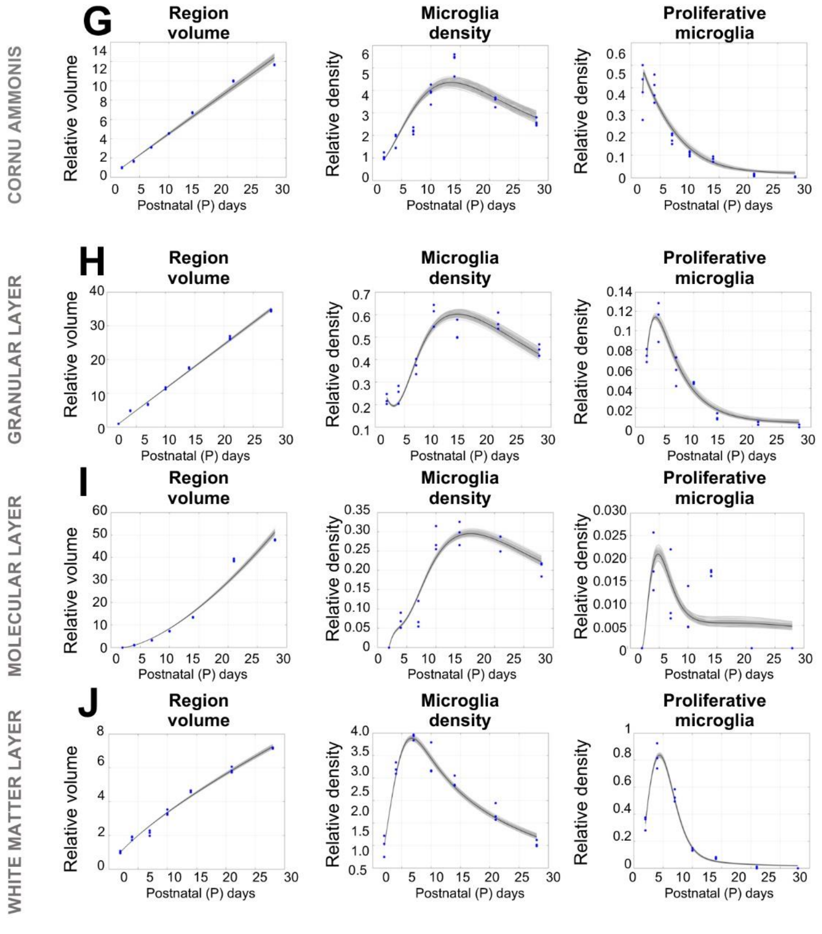

**Supplementary Figure 3. Mathematical model development in the hippocampus and cerebellum by subregions.** To develop the mathematical model, we used the data of the volume, microglial density, and proliferation from the cerebellar WM. We built a series of models using partial differential equations with increased complexity. (**A**) First, we developed a logistic growth model that assumed that cells could proliferate ($\alpha$) but also underwent competition with each other (K(t)) and apoptosis ($\beta$). The model provided a good estimation of the volume and microglial density, but not their proliferation (**B**). We developed a more complex linear model (**C**), that besides proliferation ($\alpha$) and apoptosis (C), accounted for an incoming source of cells (S(t)) external to the cerebellum. The model provided a good estimation of the volume and microglial density, but not the proliferation (**D).** (**E**) The two-population model allowed microglia to distribute into two different mathematical populations, A and B, and took into account four possible interactions: proliferation ($\alpha1$), differentiation ($\alpha2$), spreading between regions ($\beta$), and apoptosis ($C$). The proliferation and differentiation could occur by intrinsic ($K_1$) or extrinsic rates ($K_2$), the latter one involving an environmental factor related to the volume growth (E). (**F**) Structure volume, microglia density and proliferation in CA. (**G**) Structure volume, microglia density and proliferation in GL. (**H**) Structure volume, microglia density and proliferation in ML. (**I**) Structure volume, microglia density and proliferation in WM. The volume, density of microglia and proliferative cells are normalized by their respective average initial value at P2 or, for the molecular layer, at P4. Dots represented the raw data and the blue line and grey shaded area the model fit.

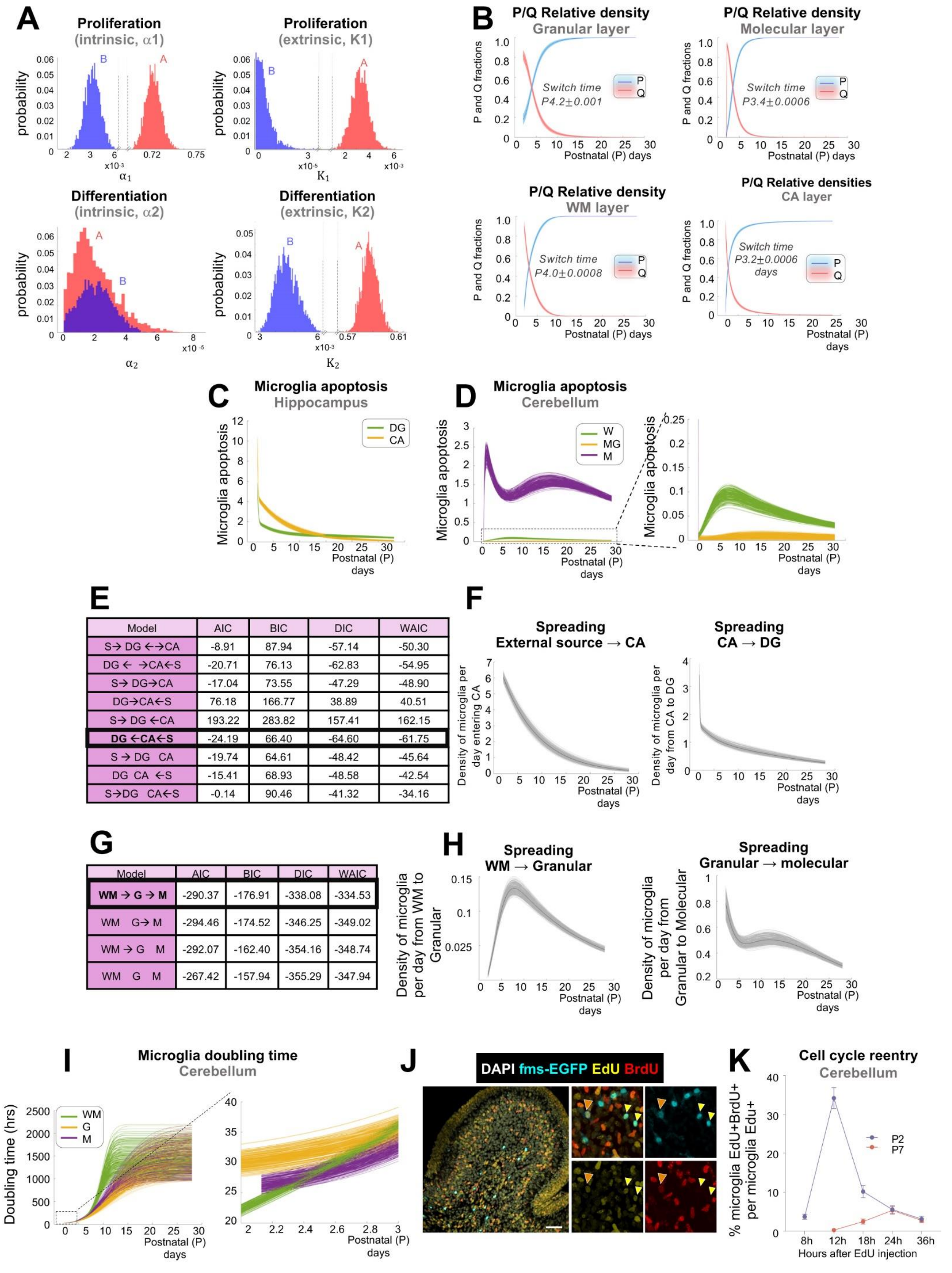


| Model | AIC | BIC | DIC | WAIC |
|---|---|---|---|---|
| S→ DG ←→CA | -8.91 | 87.94 | -57.14 | -50.30 |
| DG ← →CA←S | -20.71 | 76.13 | -62.83 | -54.95 |
| S→ DG→CA | -17.04 | 73.55 | -47.29 | -48.90 |
| DG→CA←S | 76.18 | 166.77 | 38.89 | 40.51 |
| S→ DG ←CA | 193.22 | 283.82 | 157.41 | 162.15 |
| **DG ←CA←S** | -24.19 | 66.40 | -64.60 | -61.75 |
| S → DG CA | -19.74 | 64.61 | -48.42 | -45.64 |
| DG CA ←S | -15.41 | 68.93 | -48.58 | -42.54 |
| S→DG CA←S | -0.14 | 90.46 | -41.32 | -34.16 |

| Model | AIC | BIC | DIC | WAIC |
|---|---|---|---|---|
| **WM → G → M** | -290.37 | -176.91 | -338.08 | -334.53 |
| WM G→ M | -294.46 | -174.52 | -346.25 | -349.02 |
| WM → G M | -292.07 | -162.40 | -354.16 | -348.74 |
| WM G M | -267.42 | -157.94 | -355.29 | -347.94 |

**Supplementary Figure 4. Switch from a proliferative to quiescent population identified through parameter analysis. (A)** Estimations of the intrinsic and environment-dependent proliferation and differentiation parameter distributions in the cerebellum by Markov Chain Monte Carlo (MCMC) sampling. (**B**) Relative density of the proliferative and quiescent populations with time in the granular, molecular, WM, and CA layers. Predicted microglia apoptosis in the hippocampus (**C**) and cerebellum (**D**). (**E**) Predicted directions of spreading in the hippocampus between DG, CA and an external source (S). Optimal fitness of the model was obtained when microglia spreading had an inward direction: from an external source to CA, and from CA to the DG (**F**). (**G**) Predicted directions of spreading in the cerebellum between the granular, molecular and WM layer. Optimal fitness of the model was obtained when microglia spreading had an outward direction: from the WM to the granular and to the molecular layer (**H**). **(I)** Two-population model prediction of lengthening of microglia population doubling time with age in the cerebellum. (**J**) Representative confocal images of the hippocampus from P2 fms-EGFP mouse injected with EdU and BrdU (8h apart) DAPI (white) staining shows nuclei, EdU (yellow) cells at t=0 and BrdU (red) cells at t=8h. High magnification show EdU cells (yellow arrow) and $EdU^+BrdU^+$ cells (orange arrow). (**K**) Percentage of $EdU^+BrdU^+$ microglia at P2 and P7 in the cerebellum. Bars show mean ± SEM of n=3 mice. Scale bars: 50µm. Thickness=10.5µm.

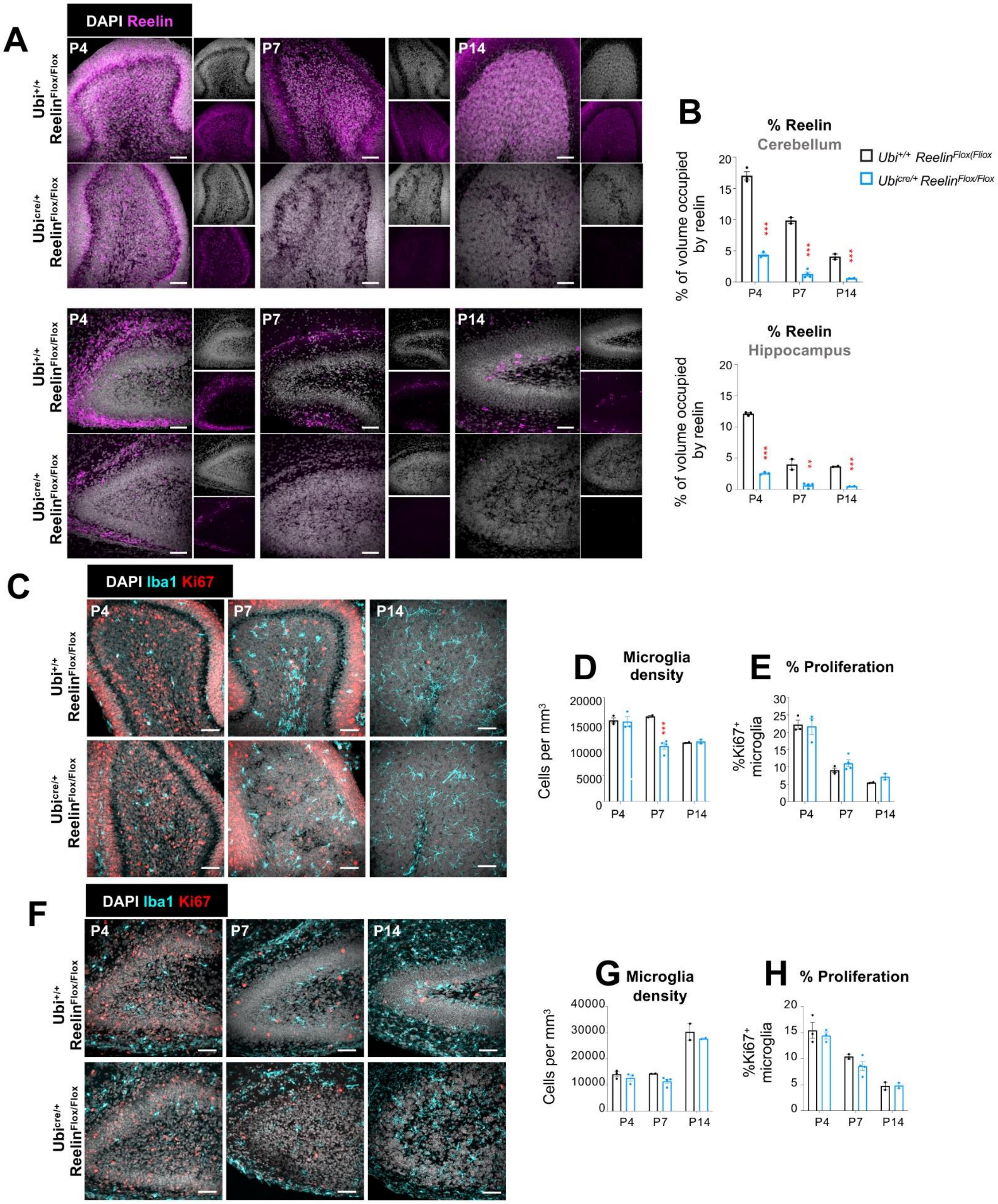


**Supplementary Figure 5. Reelin depletion delays microglia colonization in the cerebellum without major impact on microglia developmental milestones. (A)** Representative confocal images showing the cerebellum and hippocampus at P4, P7, and P14 in control and Reelin KO mice. Nuclei identified with DAPI (white) and reelin expression in magenta. (**B**) Reelin expression in the cerebellum and in the hippocampus. Bars show mean±SEM of n=3 mice (P4), n=2 control and n=4 reelin KO (P7) and n=2 (P14). (**C**) Representative confocal images of the cerebellum at P4, P7 and P14 in control and reelin KO mice. Proliferation was identified by Ki67 expression (red), microglia as Iba1 positive cells (cyan) and nuclei with DAPI (white). (**D**) Microglia density in the cerebellum. (**E**) Percentage of microglia proliferation in the cerebellum. Bars show mean±SEM of n=3 mice (P4), n=2 control and n=4 reelin KO (P7) and n=2 (P14). (**F**) Representative confocal images showing the hippocampus at P4, P7 and P14 in control and reelin KO mice. Proliferation was identified by Ki67 expression (red), microglia as Iba1 positive cells (cyan) and nuclei with DAPI (white). (**O**) Microglia density in the hippocampus. (**P**) Percentage of microglia proliferation in the hippocampus.

Bars show mean±SEM of n=3 mice (P4), n=2 control and n=4 reelin KO (P7) and n=2 (P14). Data were analyzed by two-way ANOVA followed by Bonferroni post hoc test, *p<0.05, **p<0.01, ***p<0.001 comparing control (black) vs. reelin KO (blue) mice. Scale bars (**A**): 50µm. Thickness, left to right upper panel cerebellum z=17.5µm, 18.2µm, 15.4µm; bottom panel cerebellum z= 18.2µm, 20.3µm, 17.5µm; upper panel hippocampus z=21.7µm, 20.3µm, 17.5µm; bottom panel hippocampus z=22.4µm, 21.7µm, 18.2µm. Scale bars (**C**): 50µm. Thickness, left to right upper panel z=21.7µm, 20.3µm, 17.5µm; bottom panel z=12.25µm, 11.2µm, 10.15µm. Scale bars (**F**): 50µm. Thickness, left to right upper panel z=15.4µm, 11.9µm, 10.15µm; bottom panel z=10.15µm, 14.7µm, 15.4µm bottom panel.

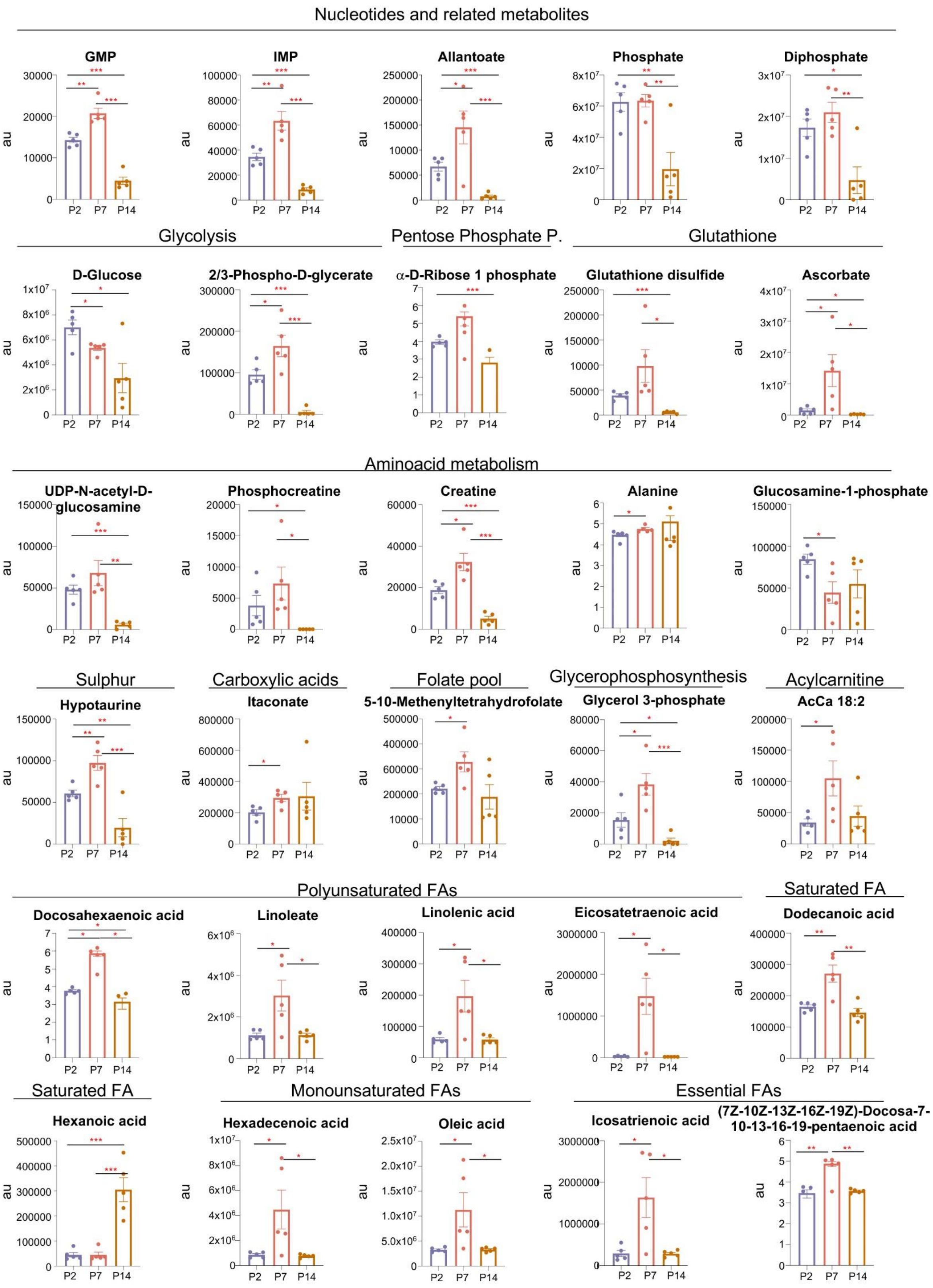
Nucleotides and related metabolites
GMP
IMP
Allantoate
Phosphate
Diphosphate
Glycolysis
Pentose Phosphate P.
Glutathione
D-Glucose
2/3-Phospho-D-glycerate
α-D-Ribose 1 phosphate
Glutathione disulfide
Ascorbate
Aminoacid metabolism
UDP-N-acetyl-D-glucosamine
Phosphocreatine
Creatine
Alanine
Glucosamine-1-phosphate
Sulphur
Carboxylic acids
Folate pool
Glycerophosphosynthesis
Acylcarnitine
Hypotaurine
Itaconate
5-10-Methenyltetrahydrofolate
Glycerol 3-phosphate
AcCa 18:2
Polyunsaturated FAs
Saturated FA
Docosahexaenoic acid
Linoleate
Linolenic acid
Eicosatetraenoic acid
Dodecanoic acid
Saturated FA
Monounsaturated FAs
Essential FAs
Hexanoic acid
Hexadecenoic acid
Oleic acid
Icosatrienoic acid
(7Z-10Z-13Z-16Z-19Z)-Docosa-7-10-13-16-19-pentaenoic acid
au
P2 P7 P14

**Supplementary Figure 6. Microglia metabolic composition during postnatal development.** Microglia metabolic composition in nucleotides and related metabolites, metabolites related to glycolysis, the pentose phosphate pathway, gluthation homeostasis amino acid metabolism and aminosugars, sulfur metabolism, carboxylic acids, metabolites of the folate fool, metabolites related to glycerophosphosynthesis, and polyunsaturated, saturated, monounsaturated, and essential fatty acids (FAs) at P2, P7 and P14 in fms-EGFP mice.

Bars show mean±SEM of n=5 mice. Data were analyzed by one-way ANOVA followed by Tukey post hoc test, *$p<0.05$, **$p<0.01$, ***$p<0.001$. Only significant differences between consecutive ages are shown.

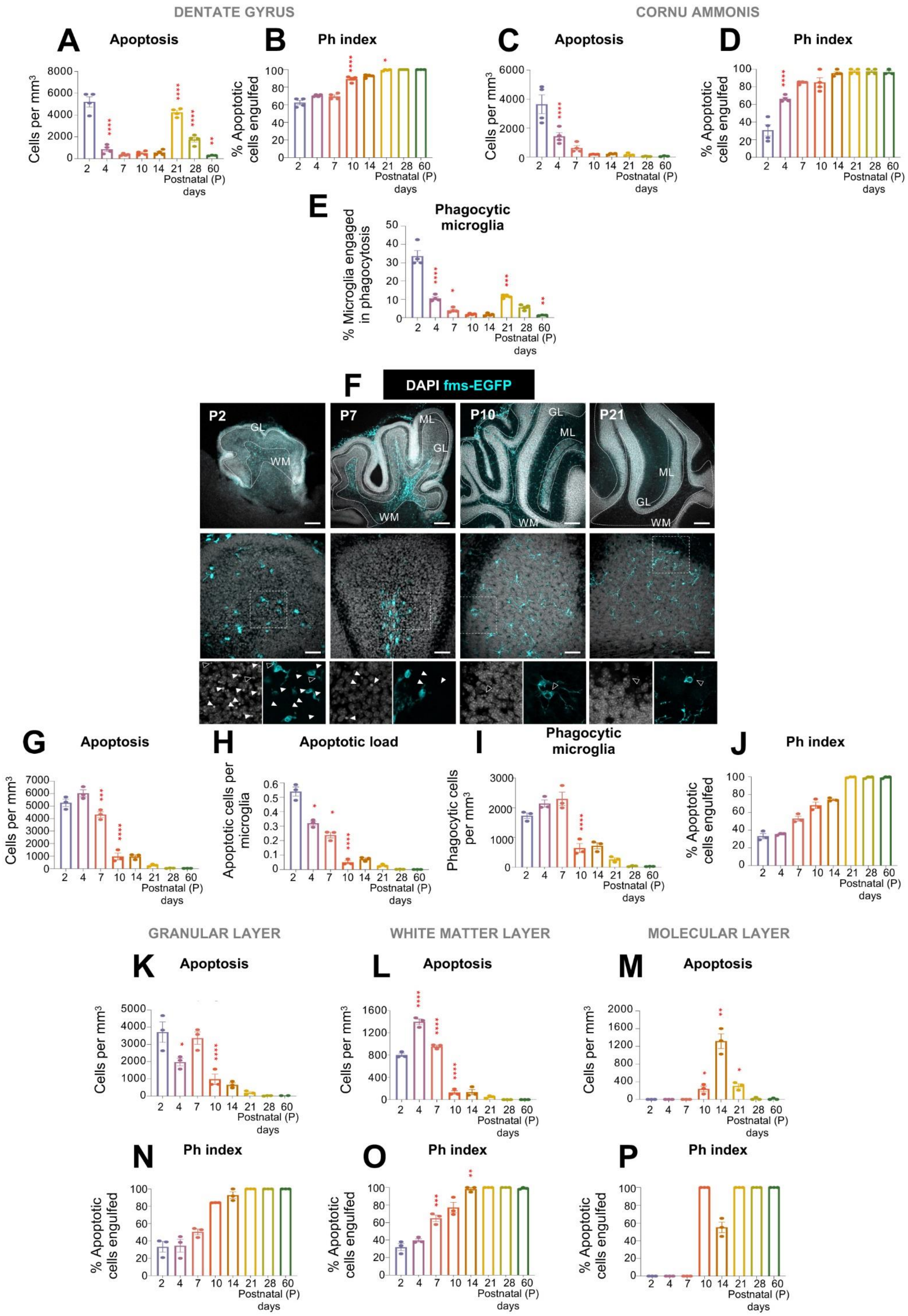
DENTATE GYRUS
CORNU AMMONIS
A Apoptosis
B Ph index
C Apoptosis
D Ph index
E Phagocytic microglia
Cells per mm³
% Apoptotic cells engulfed
% Microglia engaged in phagocytosis
Postnatal (P) days
F DAPI fms-EGFP
P2
P7
P10
P21
GL
WM
ML
G Apoptosis
H Apoptotic load
I Phagocytic microglia
J Ph index
Apoptotic cells per microglia
Phagocytic cells per mm³
GRANULAR LAYER
WHITE MATTER LAYER
MOLECULAR LAYER
K Apoptosis
L Apoptosis
M Apoptosis
N Ph index
O Ph index
P Ph index

**Supplementary Figure 7. Microglia phagocytosis efficiency across hippocampal and cerebellar subregions during postnatal development.** (**A**) Density of apoptotic cells in the DG. (**B**) Phagocytic index (% of apoptotic cells phagocytosed by microglia) in the DG. (**C**) Density of apoptotic cells in CA. (**D**) Phagocytic index (% of apoptotic cells phagocytosed by microglia) in CA. Bars show mean±SEM of n=4 mice (**E**) Representative confocal images showing the cerebellum at P2, P7, P10, and P21 in fms-EGFP mice. Apoptotic cells were identified by their condensed nuclear morphology detected by DAPI (white) and microglia as GFP positive cells (cyan). High magnification images in the white dotted square show non-engulfed (P2 and P7, white arrow) and engulfed (P10 and P21, black arrows). (**F**) Density of apoptotic cells in the cerebellum. (**G**) Clearance ratio (ratio of apoptotic cells per microglia) in the cerebellum. (**H**) Microglia phagocytic cells per cerebellum. (**I**) Phagocytic index (% of apoptotic cells phagocytosed by microglia) in the cerebellum. Density of apoptotic cells in the GL (**J**), WM (**K**) and ML (**L**). (**I**) Phagocytic index (% of apoptotic cells phagocytosed by microglia) in the GL (**M**), WM (**N**) and ML (**O**).

Bars show mean±SEM of n=4 mice (A-D) and n=3 (F-O). Data were analyzed by one-way ANOVA followed by Tukey post hoc test, *p<0.05, **p<0.01, ***p<0.001. Only significant differences between consecutive ages are shown. Scale bars: upper panel 200µm; middle panel 50µm; bottom panel 20 µm. Thickness, left to right: upper panel z=9.6µm, 11.2µm, 13.3µm, 9.6µm; middle panel z=12.6µm, 14.7µm 13.3µm 12.5µm; bottom panel z=2.1µm, 2.1µm, 2.1µm, 2.1µm.

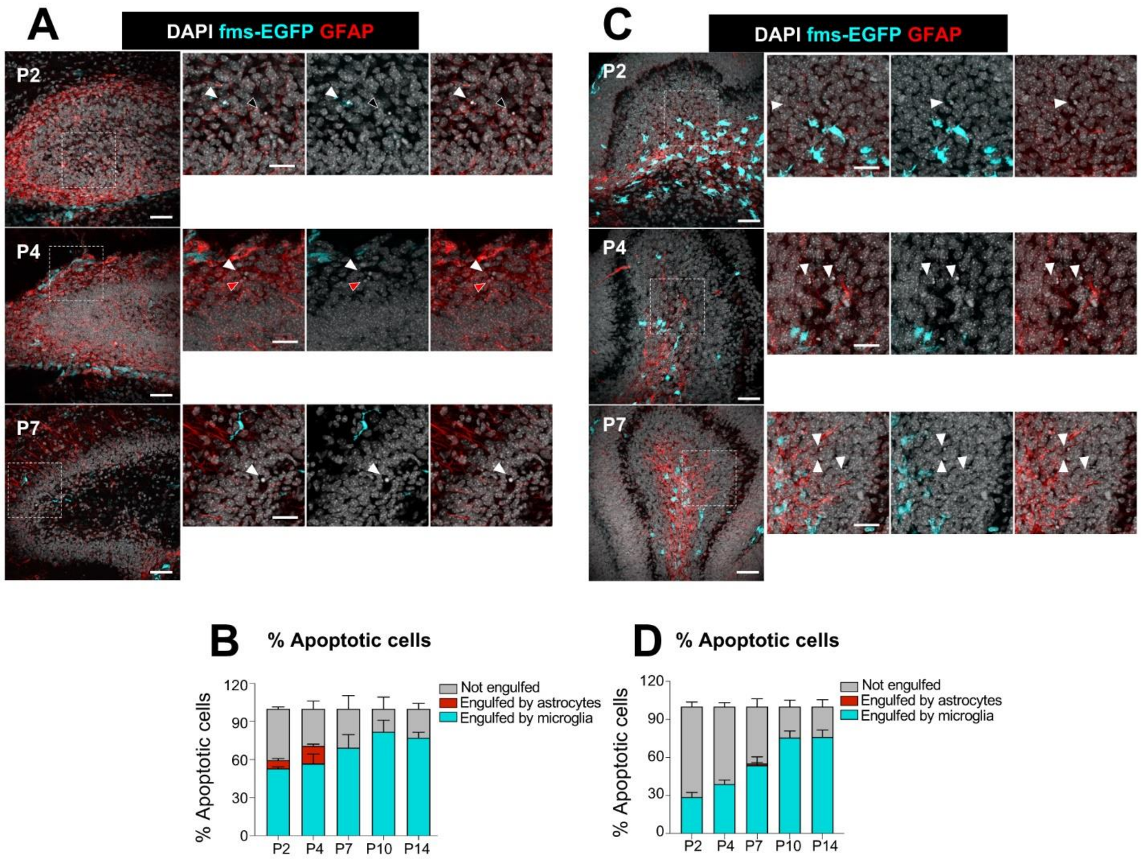


**Supplementary Figure 8. Astrocytes do not compensate for inefficient microglia phagocytosis during early postnatal development in the hippocampus and cerebellum.** (**A**) Representative confocal images showing the hippocampus at P2, P4, and P7 in fms-EGFP mice. Apoptotic cells were identified by their condensed nuclear morphology detected by DAPI (white), microglia as GFP positive cells (cyan) and astrocytes as GFAP positive cells (red). High magnification images in the white dotted square show not engulfed (white arrow), engulfed by microglia (black arrows), and engulfed by astrocytes (red arrows) apoptotic cells. (**B**) Percentage of apoptotic cells engulfed by astrocytes (red), microglia (cyan) or not engulfed (grey) in the hippocampus. (**C**) Representative confocal images showing the cerebellum at P2, P4 and P7 in fms-EGFP mice. (**D**) Percentage of apoptotic cells engulfed by astrocytes (red), microglia (cyan) or not engulfed (grey) in the cerebellum. Bars show mean±SEM of n=3 mice. Scale bars: 50µm and 20 µm square inserts. Thickness, left to right A upper panel, z=6.3µm, 6.3 µm; middle panel z=4.9µm, 4.9µm; bottom panel z=6.3µm, 6.3µm; C upper panel z=7µm, 7µm; middle panel z= 6.3µm, 6.3 µm; bottom panel 6.3µm, 6.3 µm.

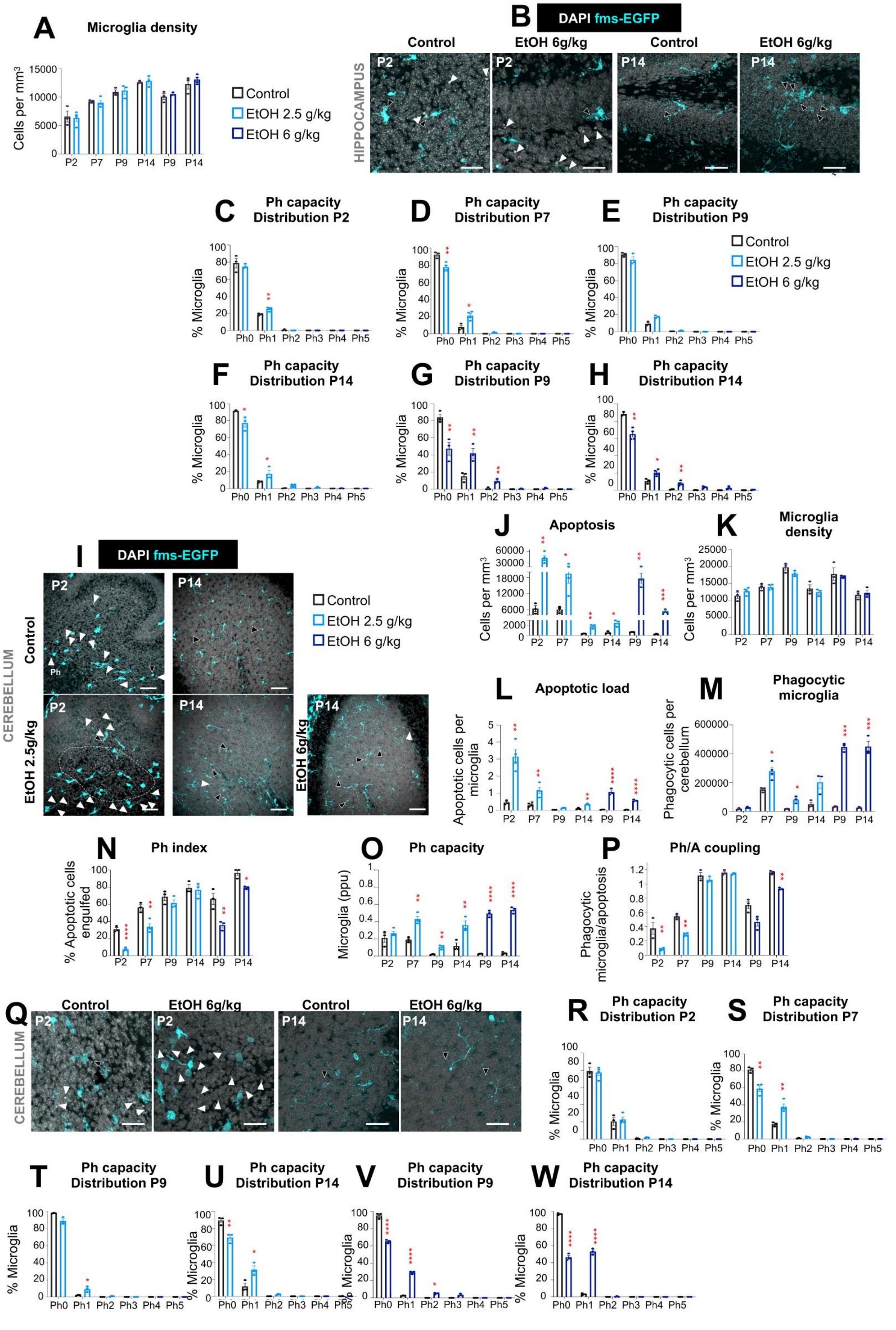
A Microglia density
Cells per mm3
Control
EtOH 2.5 g/kg
EtOH 6 g/kg
B DAPI fms-EGFP
Control
EtOH 6g/kg
HIPPOCAMPUS
C Ph capacity Distribution P2
D Ph capacity Distribution P7
E Ph capacity Distribution P9
F Ph capacity Distribution P14
G Ph capacity Distribution P9
H Ph capacity Distribution P14
% Microglia
Ph0 Ph1 Ph2 Ph3 Ph4 Ph5
I DAPI fms-EGFP
CEREBELLUM
Control
EtOH 2.5g/kg
EtOH 6g/kg
J Apoptosis
K Microglia density
Cells per mm3
L Apoptotic load
Apoptotic cells per microglia
M Phagocytic microglia
Phagocytic cells per cerebellum
N Ph index
% Apoptotic cells engulfed
O Ph capacity
Microglia (ppu)
P Ph/A coupling
Phagocytic microglia/apoptosis
Q Control EtOH 6g/kg Control EtOH 6g/kg
CEREBELLUM
R Ph capacity Distribution P2
S Ph capacity Distribution P7
T Ph capacity Distribution P9
U Ph capacity Distribution P14
V Ph capacity Distribution P9
W Ph capacity Distribution P14
P2 P7 P9 P14 P9 P14

**Supplementary Figure 9**. **Late postnatal microglia respond to ethanol-induced apoptosis by increasing pouch formation in the hippocampus and cerebellum.** (**A**) Microglia density in the hippocampus. (**B**) Representative confocal images showing the cerebellum at P2 and P14 in fms-EGFP mice in control conditions and after IP ethanol injection of 6g/kg. Apoptotic cells were identified by their condensed nuclear morphology detected by DAPI (white) and microglia as GFP positive cells (cyan). Black arrows indicate microglia phagocytosis and white arrows indicate apoptotic cells not engulfed. (**C-H**) Ph capacity distribution (proportion of microglia with one or more phagocytic pouches) in mice treated with 2.5mg/kg ethanol at P2 (C), P7 (D), P9 (E) and P14 (F) or 6mg/kg ethanol at P9 (G) and P14 (H). (**I**) Representative confocal images showing the cerebellum at P2, P7, P9, and P14 in fms-EGFP mice in control conditions and after IP ethanol injection of 2.5g/kg or 6g/kg. White dotted square indicates accumulation of apoptotic cells. (**J**) Density of apoptotic cells in the cerebellum. (**K**) Microglia density in the cerebellum. (**L**) Clearance ratio (ratio of apoptotic cells per microglia in the cerebellum). (**M**) Phagocytic microglia in the cerebellum. (**N**) Phagocytic index in the cerebellum (% of apoptotic cells phagocytosed by microglia). (**O**) Phagocytic capacity in the cerebellum (proportion of microglia with one or more phagocytic pouches). Bars show mean±SEM of n=3 mice. (**P**) Coupling between phagocytosis and apoptosis in the hippocampus. (**Q**) Representative confocal images showing the cerebellum at P2 and P14 in fms-EGFP mice in control conditions and after IP ethanol injection of 6g/kg. (**R-W**) Ph capacity distribution (proportion of microglia with one or more phagocytic pouches) in mice treated with 2.5mg/kg ethanol at P2 (R), P7 (S), P9 (T) and P14 (U) or 6mg/kg ethanol at P9 (V) and P14 (W).

Bars show mean±SEM of n=3 mice (A. C-H; J-P; R-W). Data were analyzed by multiple t-test, *p<0.05, **p<0.01, ***p<0.001 comparing vehicle (black) vs. ethanol injected (blue) mice. Scale bars (B): 50µm. Thickness, left to right z=6.3µm, 9.1µm, 7.7µm, 8.4µm. Scale bars (I): 50µm. Thickness, left to right upper panel, z=10.5µm, 11.2µm; bottom panel z=13.3µm, 10.5µm, 17.5µm. Scale bars (Q): 50µm. Thickness, left to right z=6.3µm, 9.1µm, 7.7µm, 8.4µm.

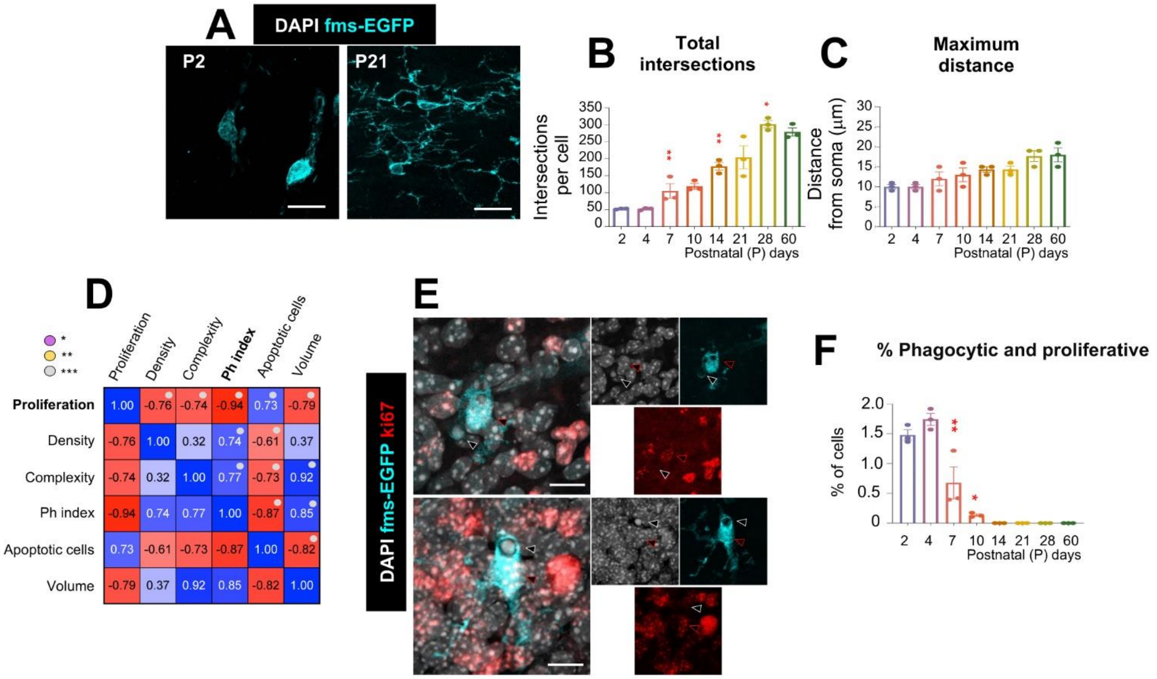


**Supplementary Figure 10. Microglia transition from an ameboid to a ramified morphology in the postnatal cerebellum, as they switch from proliferation to functional maturation.** (**A**) Representative confocal images of microglia (cyan) in the cerebellum at P2 and P21 in fms-EGFP mice. (**B, C**) Total number of intersections (reflecting the number of processes) (B) and maximum distance of the processes (C) in cerebellar microglia. Graphs show mean±SEM of n=3 in the cerebellum. (**D**) Multiple correlations in cerebellum showing the correlation coefficient R between proliferation, density, complexity (number of processes x distance), phagocytic index, density of apoptotic cells, and volume. (**E**) Representative confocal images of proliferative and phagocytic microglia in the cerebellum and hippocampus at P2 and P4 in fms-EGFP mice. Proliferative microglia were identified by Ki67 expression (red) and microglia as GFP positive cells (cyan). DAPI (white) staining shows nuclei. White arrows indicate an apoptotic cell engulfed and red arrows indicate proliferative microglia. Percentage of proliferative and phagocytic microglia over total microglia in the cerebellum (**F**).

Bars show mean ± SEM of n=3 mice (B, C, F). Data were analyzed by one-way ANOVA followed by Tukey post hoc test, *p<0.05, **p<0.01, ***p<0.001. Only significant differences between consecutive ages are shown. Scale bars (**A**): 20µm. Thickness, left to right z=5.6, 4.2µm. Scale bars (**E**): 20µm. Thickness, left to right z=2.8µm, 2.8µm.

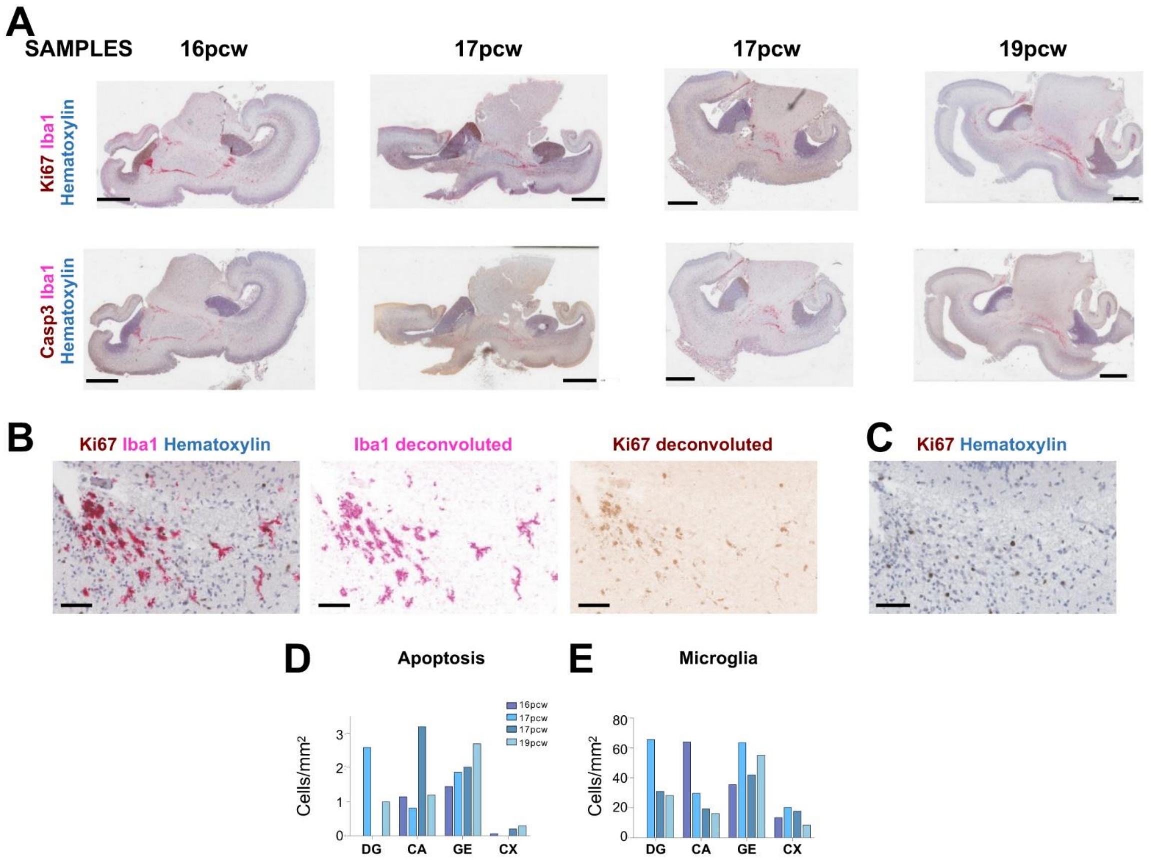


**Supplementary Figure 11. Analysis of microglial development in the human brain.** (**A**) Low magnification images of the human brain sections analyzed showing staining for Ki67 (brown, upper row) or activated caspase 3 (brown, bottom row), together with Iba1 (magenta) and hematoxylin (blue). (**B**) Example of a hot spot in the hippocampal fissure, showing the triple Ki67, Iba1 and hematoxylin staining and the resulting deconvolution. (**C**) Ki67 single staining of a consecutive section, showing fewer Ki67 cells than in the deconvoluted image shown in B. (**D**) Density of apoptotic cells across time points and regions. (**E**) Density of microglia across time points and regions.

Scale bars (A): 4mm; (B,C): 60μm.

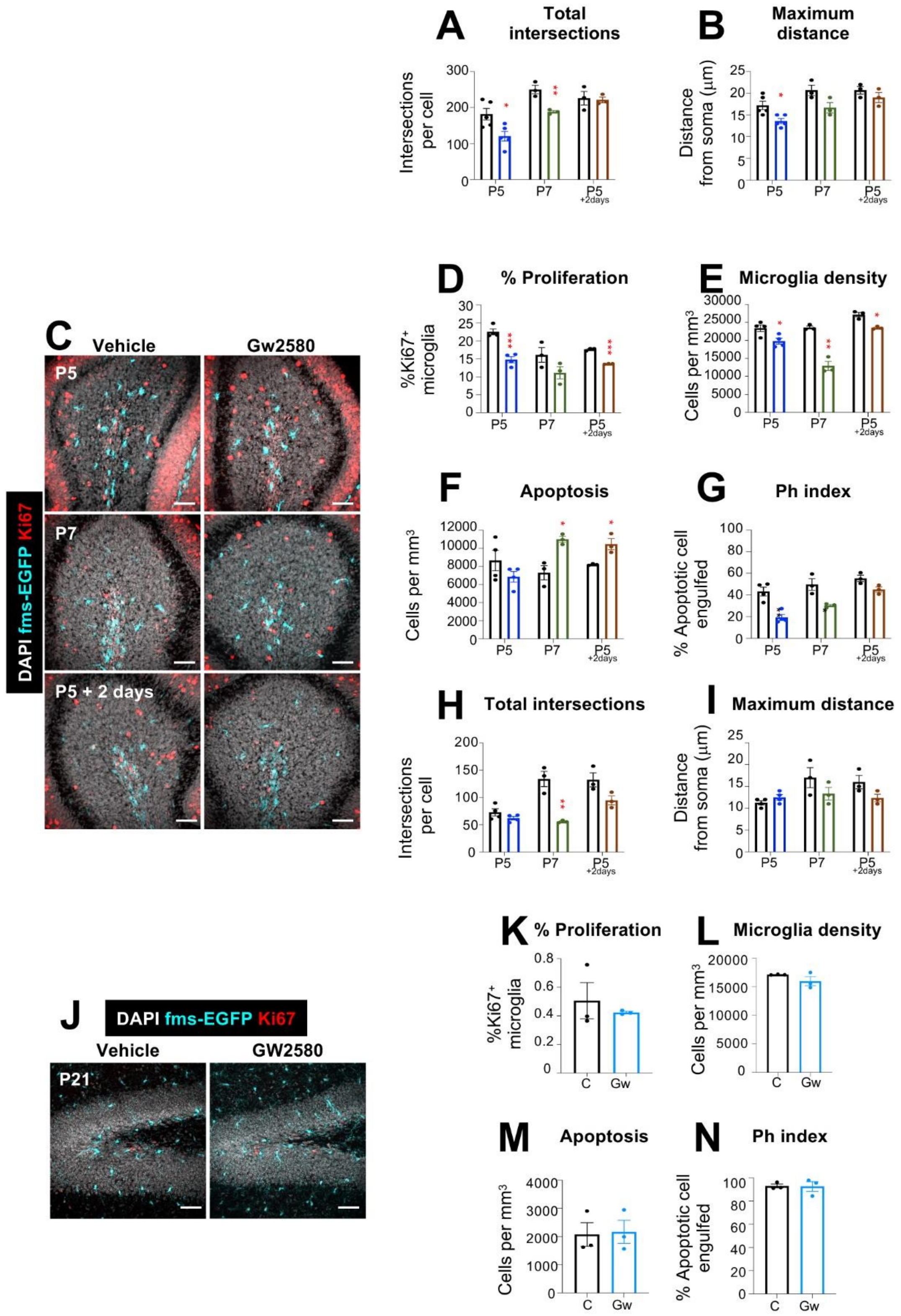


**Supplementary Figure 12. Microglia maturation in the cerebellum after CSF1R inhibition.** (**A, B**) Total number of intersections (reflecting the number of processes) (A) and maximum distance of the processes (B) in hippocampal microglia. Graphs show mean±SEM of n=5 in the hippocampus. **(C)** Representative confocal images showing the cerebellum after intraperitoneal injection of GW2580 (50mg/kg) at P5, P7 and P7 after two days of recovery without injection in fms-EGFP mice. Proliferation was identified by Ki67 expression (red), microglia as Iba1 positive cells (cyan) and nuclei with DAPI (white). (**D**) Percentage of microglia proliferation. (**E**) Microglia density. (**F**) Density of apoptotic cells. (**G**) Phagocytic index (% of apoptotic cells phagocytosed by microglia). (**H, I**) Total number of intersections (reflecting the number of processes) (H) and maximum distance of the processes (I) in cerebellar microglia. (**J**) Representative confocal images showing the hippocampus after intraperitoneal injection of GW2580 (50mg/kg) at P21 in fms-EGFP mice. (**K**) Percentage of microglia proliferation. (**L**) Microglia density. (**M**) Density of apoptotic cells. (**N**) Phagocytic index (% of apoptotic cells phagocytosed by microglia).

Bars show mean±SEM of n=5 (P5) and n=3 P7 and P5+2 days mice (A-B) and n=4 (P5) and n=3 P7 and P5+2 days mice (D-I) and n=3 (K-N). Multiple t-test, *p<0.05, **p<0.01, ***p<0.001 comparing control vs. GW2580-treated mice. Unpaired T test, p<0.05, **p<0.01, ***p<0.001 (L-O). Scale bar (**C**): 50µm. Thickness, left to right upper panel z=17.5µm, 25.2µm, 13.3µm; middle panel z=13.3µm, 17.5µm, 18.9µm; bottom panel z=16.8µm, 16.1µm, 17.5µm. Scale bar (**K**): 50µm. Thickness, left to right 25.2µm, 21.7µm.

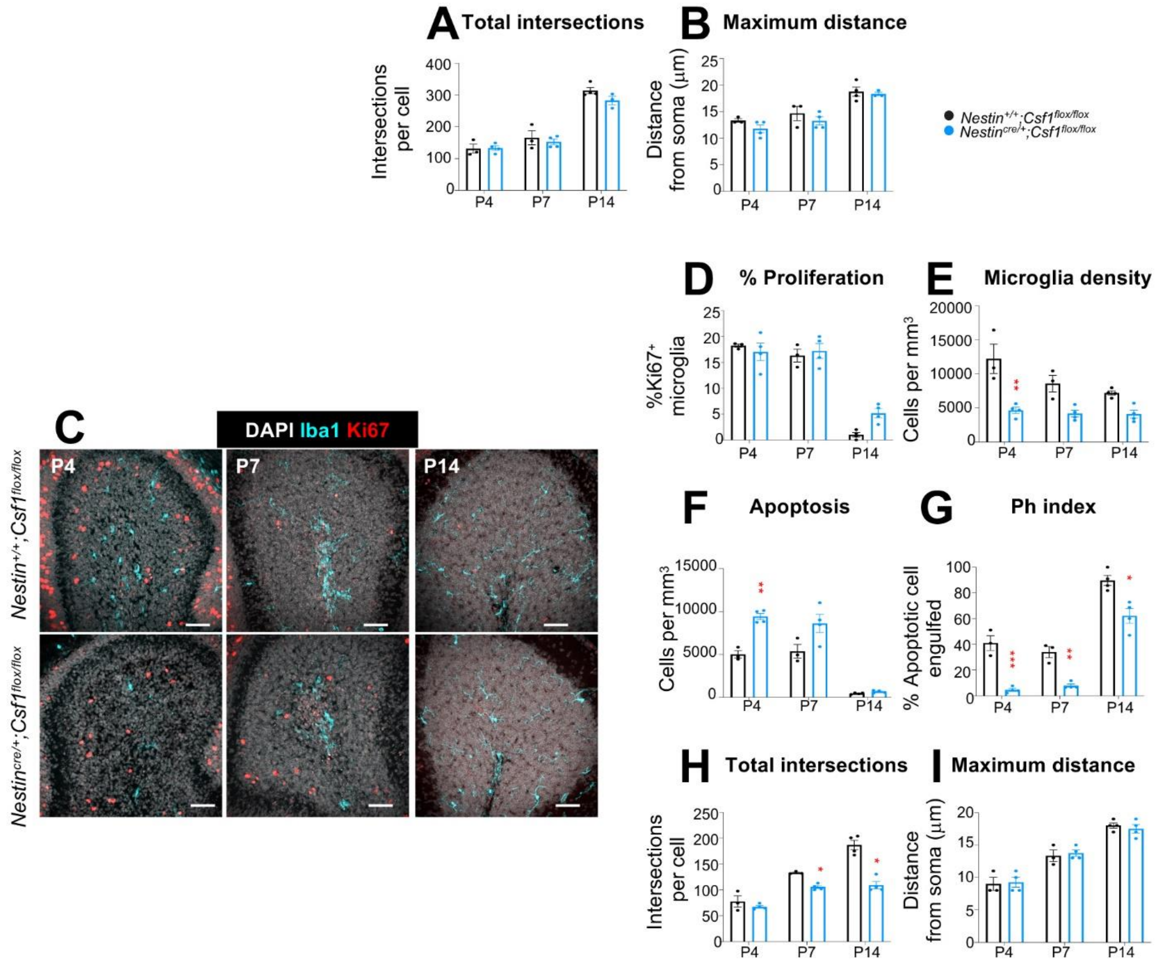


**Supplementary Figure 13. Microglia maturation in the cerebellum of a Csf1KO mouse model.** (**A, B**) Total number of intersections (reflecting the number of processes) (A) and maximum distance of the processes (B) in cerebellar microglia. Graphs show mean±SEM of n=4. **(C)** Representative confocal images showing the cerebellum in control and Csf1KO mouse model. Proliferation was identified by Ki67 expression (red), microglia as Iba1 positive cells (cyan) and nuclei with DAPI (white). (**D**) Percentage of microglia proliferation. (**E**) Microglia density. (**F**) Density of apoptotic cells. (**G**) Phagocytic index (% of apoptotic cells phagocytosed by microglia). (**H, I**) Total number of intersections (reflecting the number of processes) (H) and maximum distance of the processes (I) in cerebellar microglia.

Bars show mean±SEM of n=3 control and n=4 Csf1KO. Data were analyzed by two-way ANOVA followed by Bonferroni post hoc test *p<0.05, **p<0.01, ***p<0.001. Scale bar (**C**): 50µm. Thickness, left to right upper row z=22.4µm, 18.2µm,17.5µm, bottom row 18.2µm, 20.3µm, 22.4µm.

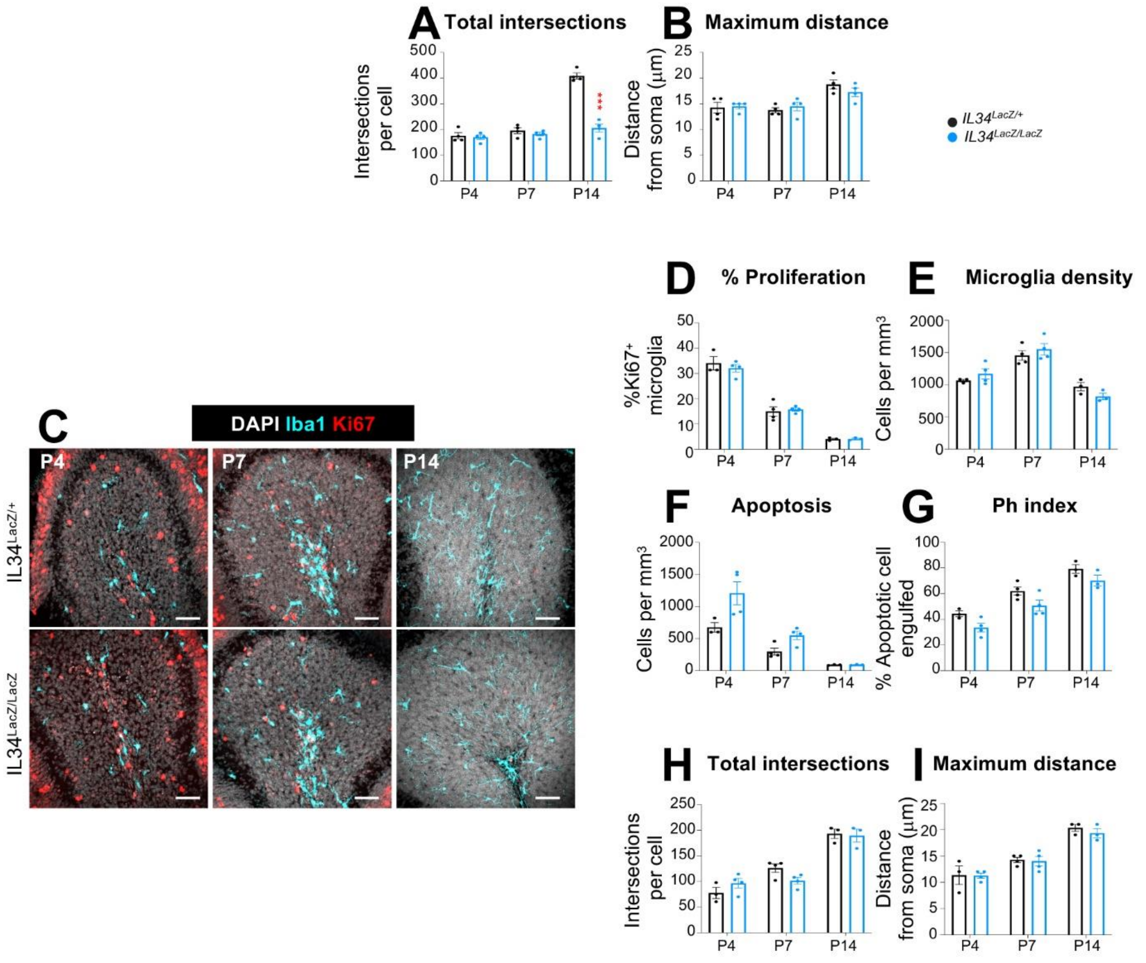


**Supplementary Figure 14. Microglia maturation in the cerebellum of a IL34KO mouse model.** (**A, B**) Total number of intersections (reflecting the number of processes) (A) and maximum distance of the processes (B) in cerebellar microglia. Graphs show mean±SEM of n=4. **(C)** Representative confocal images showing the cerebellum at P4, P7 and P14 in control (Il34$^{LacZ/+}$) and mice with ubiquitous depletion of Il34 (Il34$^{LacZ/LacZ}$). Proliferation was identified by Ki67 expression (red), microglia as Iba1 positive cells (cyan) and nuclei with DAPI (white). (**D**) Percentage of microglia proliferation. (**E**) Microglia density. (**F**) Density of apoptotic cells. (**G**) Phagocytic index (% of apoptotic cells phagocytosed by microglia). (**H, I**) Total number of intersections (reflecting the number of processes) (H) and maximum distance of the processes (I) in cerebellar microglia.

Bars show mean±SEM of n=4 (A,B), n=3 P4 control and P14, and n=4 P4 Il34 KO and P7 mice(D-I). Data were analyzed by two-way ANOVA followed by Bonferroni post hoc test *p<0.05, **p<0.01, ***p<0.001.. Scale bar (**C**): 50µm. Thickness, left to right upper panel A1 z=17.5µm, 18.2µm, 15.4µm, bottom panel z=22.4µm, 21.7µm, 18.2µm.

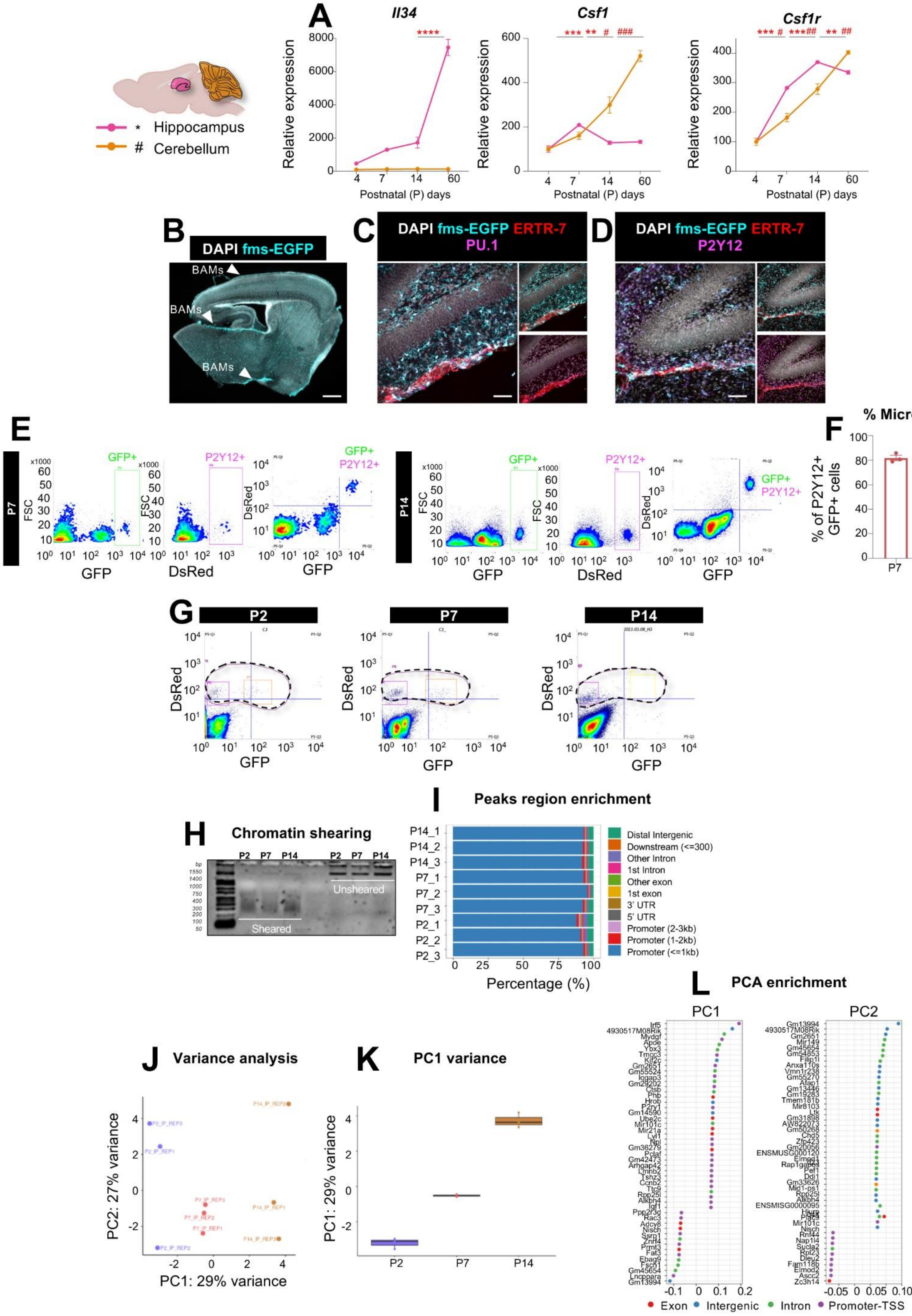
A
Il34
Csf1
Csf1r
Hippocampus
Cerebellum
Relative expression
Postnatal (P) days
B
DAPI fms-EGFP
BAMs
C
DAPI fms-EGFP ERTR-7 PU.1
D
DAPI fms-EGFP ERTR-7 P2Y12
E
GFP+
P2Y12+
FSC
DsRed
GFP
P7
P14
F
% Microglia
% of P2Y12+ GFP+ cells
G
P2
P7
P14
H
Chromatin shearing
Sheared
Unsheared
I
Peaks region enrichment
Distal Intergenic
Downstream (<=300)
Other Intron
1st Intron
Other exon
1st exon
3' UTR
5' UTR
Promoter (2-3kb)
Promoter (1-2kb)
Promoter (<=1kb)
Percentage (%)
J
Variance analysis
PC2: 27% variance
PC1: 29% variance
K
PC1 variance
PC1: 29% variance
L
PCA enrichment
PC1
PC2
Exon
Intergenic
Intron
Promoter-TSS

**Supplementary Figure 15. Isolation of a highly enriched hippocampal microglia population for ChIP-sequencing. (A)** Expression levels of Il34, Csf1 and Csf1r mRNA from hippocampi and cerebellum at P2, P7, P14 and P60 of fms-EGFP mice, normalized with Gapdh. Data was analyzed by one-way ANOVA followed by Tukey post hoc test, # *p<0.05, ## **p<0.01, ### ***p<0.001 (# for the cerebellum, * for the hippocampus). Only significant differences between consecutive ages are shown. Experimental design drawing created with BioRender.com. (**B**) Representative image of a sagittal brain slice at P7 in fms-EGFP mice. Microglia and macrophages from the meninges (BAMs) identified as GFP positive cells (cyan) and nuclei by DAPI (white). White arrows indicate macrophages from the meninges. (**C**) Representative confocal image of the hippocampus at P14 in fms-EGFP mice. Meninges identified by ERTR-7 (red) and microglia and macrophages nuclei by PU.1 (magenta). (**D**) Meninges identified by ERTR-7 (red) and microglia by P2Y12 (magenta). **(E**) Flow cytometry analysis of Pu.1 cells identified as microglia (fms-$GFP^+$, $P2Y12^+$) at P7 in fms-EGFP mice in the hippocampus. (**F**) Percentage of microglia in the hippocampus at P7 and P14. (G) $Pu.1^+$ microglia nuclei isolation by FACS-sorting at P2, P7 and P14. (**H**) Chromatin fragmentation with a mean size distribution of 300 bp. (**I**) Sequencing with H3K4me3 with peak region enrichment located in promoters at a distance of less than 1 kb. (**J**) PCA analysis between H3k4me3 immunoprecipitated samples (n=3 independent samples, from n=7 P2, n=4 P7 and n=2 P14 pooled mice) from $Pu.1^+$ sorted microglia from P2, P7 and P14 fms-EGFP hippocampi. (**K**) PC1 variance. (**L**) Fifteen genes with the highest weights for both PC1 and PC2.

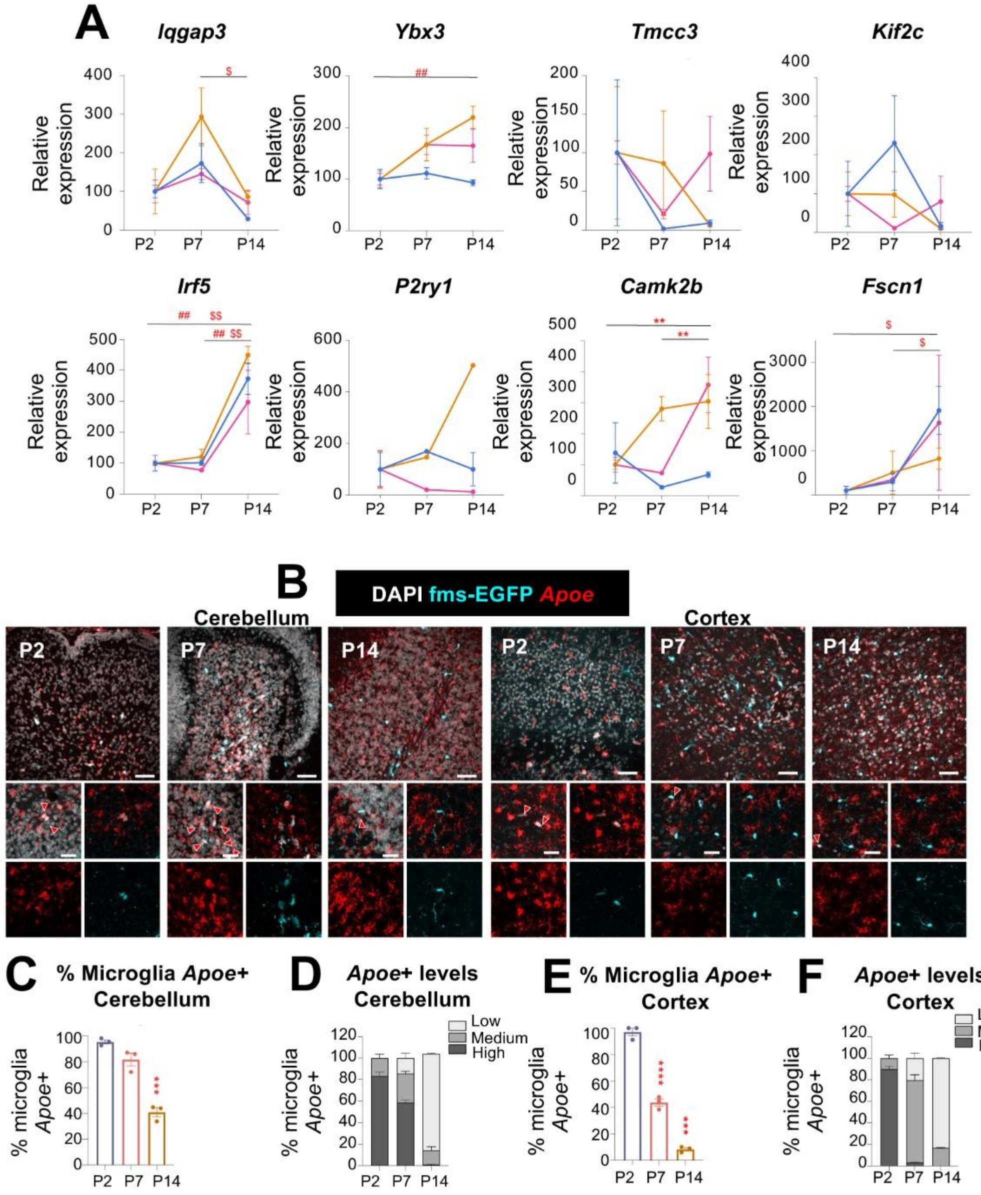


**G**

Increased expression — Decreased expression — Not changed — Not found

| | DATABASE | SPECIES | AGES | REGION | POPULATION | ANALYSIS METHOD |
|---|---|---|---|---|---|---|
| 1 | Matcovitch et al., 2016 | Mouse | Embryonic: E10.5 E 11.5 E12.5 E13.5 E16.5 / Postnatal: P3 P6 P9 / Adult | Whole brain, except Adult hippocampus | Microglia | Bulk RNAseq |
| 2 | Hammond et al., 2019 | Mouse | Embryonic: E14.5 / Postnatal: P4/P5, P30, P100, P50 | Whole brain | Microglia | scRNAseq |
| 3 | Hochgemer et al., 2018 | Mouse | Embryonic: E16.5 / Postnatal: P0, P5, P19, P23, P120, P132 | Hippocampus | Microglia cluster | scRNAseq |
| 4 | Han Claudia et al., 2023 | Human | Fetal: 10w-12w 13w-17w / Postnatal: 6w-21, 2y-9y 13y-17y | Fetal whole brain, postnatal cortical tissue derived from epileptic resections of patients | Microglia | Microglia bulk RNAseq |
| 5 | Yaqubi et al., 2023 | Human | prenatal: GW13, GW18, GW19 / pediatric: 1.5, 2, 2 years / adolescent: 10, 13, 14 years / adult: 41, 51, 62 years | Unknown: *"collected tissues were obtained from normal appearing, superficial tissue, removed to access underlying pathology"* | Microglia | scRNAseq |

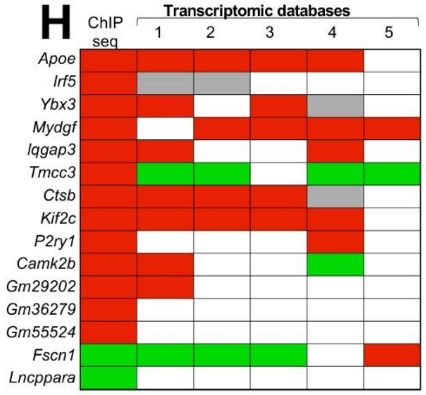

**Supplementary Figure 16. The expression pattern of the target genes aligns with their epigenetic landscape.** (**A**) Expression levels of the differentially methylated genes identified by ChIP-seq H3k4me3 mark from $Pu.1^{+}$ sorted microglia at P2, P7, and P14 in the hippocampus, cerebellum and cortex of fms-EGFP mice (n=3 per region). Relative expression *Iqgap3, Ybx3, Tmcc3, Kif2c, Irf5, P2ry1, Camk2b, Fscn1*. (**B**) Representative confocal images showing the cerebellum and the cortex at P2, P7 and P14 in fms-EGFP mice. Microglia identified as GFP positive cells (cyan) and Apoe mRNA by red. High magnifications images show representative microglia from each age with Apoe expression (red arrow). (**C**) Percentage of microglia Apoe+ in the cerebellum. (**D**) Apoe+ microglia levels in the cerebellum. **(E**) Percentage of microglia Apoe+ in the cortex. (**F**) Apoe+ microglia levels in the cortex. (**G**) Table with the tendency expression pattern (green: increased, red: decreased, grey: not changed, white: not found) of our target genes in the available databases from RNA sequencing studies: 1. (Matcovitch-Natan et al., 2016), 2. (Hammond et al., 2019), 3. (Hochgerner et al., 2018), 4. (Han et al., 2023), 5. (Yaqubi et al., 2023). (**H**) Summary heatmap of the tendency of expression pattern for the different target genes in the different databases.

Bars show mean±SEM of n=3 (A, C-F). One-way ANOVA, *p<0.05, **p<0.01, ***p<0.001. Scale bar (**B**): 50µm upper row, 20µm lower row. Thickness, left to right z= 6.3µm, 6.3µm, 7.7µm, 6.3µm, 9.1µm, 7.7µm.

**Supplementary Table 1. Case demographics.** The table summarizes the case identifier, the gestational age (GA), the postconceptional week (PCW), the sex (M, male; F, female), the source (OBB, Oxford Brain Bank), the cause of death and whether there were any histological abnormalities observed.

| **Case** | **GA** | **PCW** | **Sex** | **Source** | **Cause of Death** | **Histology** |
|---|---|---|---|---|---|---|
| Fetus 1 (NP41-2014) | 18 | 16 | M | OBB | Termination of pregnancy | Normal |
| Fetus 2 (B5349) | 19 | 17 | M | OBB | Termination of pregnancy | Normal |
| Fetus 3 (NP53-2012) | 19 | 17 | F | OBB | Termination of pregnancy | Normal |
| Fetus 4 (NP155-2010) | 21 | 19 | M | OBB | Termination of pregnancy | Normal |

**Supplementary mathematical model**

Volume of cerebellum and hippocampus

Functions to represent the volume of the cerebellar and hippocampal regions are fit to the experimental data. Based on qualitative observation, it was determined that the volume of each region grew at either a linear rate or according to a power law. The equations used to fit the volume of the cerebellum and hippocampus regions can be found below.

$$V_W(t) = V_W^0 \left(\frac{t}{t_0}\right)^{\gamma_w}, \qquad (1)$$

$$V_G(t) = V_G^0 \cdot \left(\gamma_g(t - t_0) + 1\right), \qquad (2)$$

$$V_M(t) = V_M^1 \cdot \left(\frac{t - t_0}{t_1 - t_0}\right)^{\gamma_m}, \qquad (3)$$

where $t_0$ denotes the initial time of the experiments (here, $t_0$is day P2) and $t_1$ the first day of the experiment in which the molecular layer has non-zero volume (here, $t_1$ is day P4). The constants $V_W^0$, $V_G^0$, and $V_M^1$ denote the volume of the white matter, granular, and molecular layers at day P2, P2, and P4, respectively. The parameters $\gamma_w$, $\gamma_g$, and $\gamma_m$ are fit to the experimental data (see paragraphs below for more details).

The volumes of the DG and CA regions of the hippocampus are similarly determined by

$$V_D(t) = G_D \left(\frac{t}{t_0}\right)^{\gamma_d} + (V_D^0 - G_D), \qquad (4)$$

$$V_C(t) = V_C^0 \cdot (\gamma_c(t - t_0) + 1), \qquad (5)$$

where $V_D^0$and $V_C^0$ refer to the volume of the DG and CA layers at day P2, respectively, and $G_D$, $\gamma_d$ and $\gamma_c$ are parameters that must be fit to data.

Logistic model

We adapt a common logistic model for population growth to account for the increasing-then-decreasing behavior of microglia cell densities during the experiments. The classical logistic model assumes that the cell density grows exponentially at low densities, but at high densities becomes constant due to increased competition between cells for resources and space. A key component of the classical logistic model is that cells undergo apoptosis upon contacting each other and is represented by a term commonly referred to as the "carrying capacity", which is equal to the long-time density that cells attain (i.e., the equilibrium density). We modify the logistic model to have the carrying capacity vary with time, because the framework otherwise cannot yield non-monotonic cell densities such as those observed in experiments. Since the units of the external factor are inversely proportional to volume, we make a simplifying assumption that the time-dependence of the carrying capacity varies proportionally to the volume of each cerebellar region. Furthermore, we represent cell apoptosis and spreading between different cerebellar regions by introducing terms that, for simplicity, are assumed to be linearly proportional to the cell density. The rates at which the density of microglia in the white matter, granular, and molecular layers at time $t$ (given by $W(t)$, $G(t)$, and $M(t)$, respectively) change are thus represented as the following system of ordinary differential equations (ODEs):

$$\frac{\mathrm{d}W}{\mathrm{d}t} = \alpha_W W - \alpha_W W^2 \cdot \frac{V_W(t)}{N_W} - \beta_W W + \beta_G G - c_W W, \qquad (6)$$

$$\frac{\mathrm{d}G}{\mathrm{d}t} = \alpha_G G - \alpha_G G^2 \cdot \frac{V_G(t)}{N_G} - \beta_G G - B_G G + \beta_W W + B_M M - c_G G, \qquad (7)$$

$$\frac{\mathrm{d}M}{\mathrm{d}t} = \alpha_M M - \alpha_M M^2 \cdot \frac{V_M(t)}{N_M} - B_M M + \beta_G G - c_M M, \qquad (8)$$

where $\alpha_W$, $\alpha_G$, $\alpha_M$, $N_W$, $N_G$, $N_M$, $\beta_W$, $\beta_G$, $B_G$, $B_M$, $c_W$, $c_G$, $c_M$ are parameters that must be fit to the datasets (see the paragraphs below for more details). The volumes of the white matter, granular, and molecular regions ($V_W(t)$, $V_G(t)$, $V_M(t)$, respectively) are the same as those in Eqn.s (1) – (3).

Linear model

The linear model offers an alternative framework to represent how microglia densities change over time. In the framework, cells can either proliferate, spread to different regions, or undergo apoptosis. It offers a simpler interpretation than the logistic model because all these processes are assumed to occur at linear rates, by which we mean that the time it takes for the population to double (or halve) due to one of these mechanisms is constant throughout the experiment. However, linear models cannot exhibit non-monotonic behavior that we see in the available data without introducing an additional layer of complexity. To achieve this, we assume that microglia can spread to the cerebellum regions from other areas of the brain at some (possibly time-dependent) rate. These assumptions can be combined to write a system of ODEs for how the density of microglia cells change within the different cerebellar layers. Using the same notations as before, these are given by

$$\frac{\mathrm{d}W}{\mathrm{d}t} = \alpha_W W - \beta_W W + \beta_G G - c_W W + S_W(t), \qquad (9)$$

$$\frac{\mathrm{d}G}{\mathrm{d}t} = \alpha_G G - \beta_G G - B_G G + \beta_W W + B_M M - c_G G + S_G(t), \qquad (10)$$

$$\frac{\mathrm{d}M}{\mathrm{d}t} = \alpha_M M - B_M M + \beta_G G - c_M M + S_M(t), \qquad (11)$$

where $S_W(t)$, $S_G(t)$, and $S_M(t)$are the functions denoting the rate at which cells enter the white matter, granular, and molecular layers from regions beyond the cerebellum. The values of $\alpha_W$, $\alpha_G$, $\alpha_M$, $\beta_W$, $\beta_G$, $B_G$, $B_M$, $c_W$, $c_G$, $c_M$ are constant parameters that must be fit to the datasets (see the paragraphs below for more details). To close this system of ODEs, we must assume the form that these source functions have. While there are many choices available, we decided for simplicity to use exponentially decaying functions for each layer, reasoning that cells would be less likely to spread to different regions as the mouse ages. In other words, we assume that

$$S_W(t) = S_W^0 e^{-t/\tau_W}, \qquad (12)$$

$$S_G(t) = S_G^0 e^{-t/\tau_G}, \qquad (13)$$

$$S_M(t) = S_M^0 e^{-t/\tau_M}, \qquad (14)$$

where $S_W^0$, $S_G^0$, $S_M^0$, $\tau_W$, $\tau_G$, and $\tau_M$ are all constant parameters that will be fit based on the data.

Two-population model

For the final model we consider that the microglia population can be decomposed into an "A" subtype and a "B" subtype. Both subtypes are described with identical mechanisms, leading to equivalent sets of ODEs for their dynamics, and thus are only distinguished when their parameters are fit to data. Both cell types can proliferate, undergo apoptosis, and spread to different compartments. Additionally, we account for the possibility that cells can switch between (or, in other words, "differentiate" into) the A and B types. We assume that all these mechanisms occur at linear rates, as this is the simplest assumption to make. We refer to these terms as "natural" or "intrinsic", since they do not depend on cell types beyond the microglia themselves. However, as noted above, linear models cannot explain the observed non-monotonic cell densities unless the latter oscillate in time. To resolve this problem, we added additional terms in the ODEs for proliferation and A-B switching that vary in time, and which assume that the A and B cell type interact with an “environmental factor”, which is distinct from microglia but is also present in the cerebellum (this other cell type is intentionally vague; it could include neurons or other glial cells such as astrocytes). This leads to the following system of ODEs, which calculate the rate at which the densities of A and B microglia in the white matter, granular, and molecular regions of the cerebellum (denoted by $A_W$, $B_W$, $A_G$, $B_G$, $A_M$, and $B_M$, respectively) change as a function of time, $t$:

$$\frac{\mathrm{d}A_W}{\mathrm{d}t} = (\alpha_1 - \alpha_2 - \beta_{w,1} - c_{w,1})A_W + B_{g,1}A_G + d_B B_W + (k_{w,1} - k_{w,2})A_W E_W(t) + k_{w,3}E_W(t)B_W, \quad (15)$$

$$\frac{\mathrm{d}B_W}{\mathrm{d}t} = (\alpha_3 - d_B - \beta_{w,2} - c_{w,2})B_W + B_{g,2}B_G + \alpha_2 A_W + (k_{w,4} - k_{w,3})B_W E_W(t) + k_{w,2}E_W(t)A_W, \quad (16)$$

$$\frac{\mathrm{d}A_G}{\mathrm{d}t} = (\alpha_1 - \alpha_2 - \beta_{g,1} - c_{g,1})A_G + B_{m,1}A_M + d_B B_G + (k_{g,1} - k_{g,2})A_G E_G(t) + k_{g,3}E_G(t)B_G + \beta_{w,1}A_W - B_{g,1}A_G, \quad (17)$$

$$\frac{\mathrm{d}B_G}{\mathrm{d}t} = (\alpha_3 - d_B - \beta_{g,2} - c_{g,2})B_G + B_{m,2}B_M + \alpha_2 A_G + (k_{g,4} - k_{g,3})B_G E_G(t) + k_{g,2}E_G(t)A_G + \beta_{w,2}B_W - B_{g,2}B_G, \quad (18)$$

$$\frac{\mathrm{d}A_M}{\mathrm{d}t} = (\alpha_1 - \alpha_2 - c_{m,1})A_M + \beta_{g,1}A_G + d_B B_M + (k_{m,1} - k_{m,2})A_M E_M(t) + k_{m,3}E_M(t)B_M - B_{m,1}A_M, \quad (19)$$

$$\frac{\mathrm{d}B_M}{\mathrm{d}t} = (\alpha_3 - d_B - c_{m,2})B_M + \beta_{g,2}B_G - B_{m,2}B_M + \alpha_2 A_W + (k_{m,4} - k_{m,3})B_M E_M(t) + k_{m,2}E_M(t)A_M, \quad (20)$$

where $E_W(t)$, $E_G(t)$, and $E_M(t)$ denote the density of environmental factors at time $t$ and the other terms correspond to constant parameters, which are fit based on the data (see the paragraphs below for more details on the fitting procedure). The total density of microglia within each layer can be determined by adding the densities of their respective A and B densities.

We reason that, since environmental factors exist alongside microglia within each respective layer of the cerebellum, the density of these environmental factors can be taken to be approximately proportional to the volume of the layer in which it resides. In other words, we assume that

$$E_W(t) = F_1 \cdot V_W(t),$$

$$E_G(t) = F_2 \cdot V_G(t),$$

$$E_M(t) = F_3 \cdot V_M(t),$$

for some constants $F_1$, $F_2$, and $F_3$. In principle, other functions could be used to describe how the environmental factor changes in time, although we have not considered them here.

Modeling proliferation

We construct a system of ODEs to estimate the density of Ki67+ cells within each layer of the cerebellum. For the logistic, linear, and two-population model, these systems of ODEs follow the same basic design. Specifically, terms in the models related to proliferation are also included in the system of ODEs describing the density of Ki67+ cells, since the marker begins to be expressed when cells begin to undergo division. We also assume that cells do not instantaneously stop expressing Ki67 after division is completed. For simplicity, we assume that the rate at which the population of cells stops expressing Ki67 is linear; that is, the time it takes for the density of Ki67+ cells to halve would be constant if no other cells were to divide.

For the logistic model, this leads us to the following ODEs for the Ki67+ cell density in the white matter, $K_W(t)$, granular, $K_G(t)$, and molecular, $K_M(t)$, regions:

$$\frac{\mathrm{d}K_W}{\mathrm{d}t} = \alpha_W W - \delta_W K_W, \qquad (21)$$

$$\frac{\mathrm{d}K_G}{\mathrm{d}t} = \alpha_G G - \delta_G K_G, \qquad (22)$$

$$\frac{\mathrm{d}K_M}{\mathrm{d}t} = \alpha_M M - \delta_M K_M, \qquad (23)$$

where $\delta_W$, $\delta_G$, and $\delta_M$ are parameters to be fit to the data. Note that we have assumed that the time that it takes to halve a density of Ki67+ expressing cells is assumed to not depend on the layer.

The Ki67+ ODE system corresponding to the linear model is equivalent to Eqn.s (21)–(23). The Ki67+ ODE system corresponding to the two-population model is given by:

$$\frac{\mathrm{d}K_W}{\mathrm{d}t} = \alpha_1 A_W + \alpha_3 B_W + k_{w,1} A_W E_W(t) + k_{w,4} E_W(t) B_W - \delta K_W, \qquad (24)$$

$$\frac{\mathrm{d}K_G}{\mathrm{d}t} = \alpha_1 A_G + \alpha_3 B_G + k_{g,1} A_G E_G(t) + k_{g,4} E_G(t) B_G - \delta K_G, \qquad (25)$$

$$\frac{\mathrm{d}K_M}{\mathrm{d}t} = \alpha_1 A_M + \alpha_3 B_M + k_{m,1} A_W E_W(t) + k_{m,4} E_M(t) B_M - \delta K_M, \qquad (26)$$

where $A_W(t)$, $B_W(t)$, $A_G(t)$, $B_G(t)$, $A_M(t)$, and $A_M(t)$ denote the density of A or B cells in the white matter, granular, or molecular layer, respectively.

Simulation details

The systems of ODEs were solved in Python (version 3.10.12), using an implicit first-order backward differentiation formula from the SciPy library[96]. For the fitting process, data corresponding to cell densities are normalized by the average initial white matter density (for the cerebellum data) or by the average initial dentate gyrus density (for the hippocampus data). As such, the initial white matter and dentate gyrus densities are set equal to one cell $\cdot$ mm$^{-3}$. Cell densities within other layers (except for the molecular layer within the cerebellum, which was set equal to zero because that region has zero initial volume) were treated as additional model parameters, and thus were sampled and fit according to the data. The volume within each layer is also normalized by its respective average initial value at day P2 (or, for the molecular layer, at day P4), so that the constants $V_W^0$, $V_G^0$, $V_M^1$, $V_D^0$ and $V_C^0$ in Eqn.s (1)–(5) are equal to one mm$^{-3}$.

Parameter estimation through MCMC sampling

The experimental datasets are composed of direct measurements for the density of microglia, density of Ki67+ microglia, and volumes within the white matter, granular, and molecular layers for the cerebellum and the dentate gyrus and cornu ammonis layers for the hippocampus. Data are reported at seven non-equally spaced time points from day P2 (the initial date of the experiment) to P28. Three measurements are provided for each time point in the cerebellum dataset, while four measurements at each time point are provided for the hippocampus dataset. For the parameter estimation process, we summarize experimental measurements in an observation vector $\boldsymbol{y} = (y_1, y_2, \ldots)$where each component, $y_i$, denotes a total cell density, Ki67+ cell density, or volume measurement within a given layer. For instance, for the cerebellum, $\boldsymbol{y}$ consists of measurements of twenty-seven measurements corresponding to the total and proliferating cell densities, and volumes, for each of the layers ($\boldsymbol{y} \in \mathbb{R}^{\mathbf{9x3}}$).

We relate experimental measurements, $\boldsymbol{y}_D(t)$, to model predictions, $\boldsymbol{y}(t)$, via an additive independent Gaussian error model:

$$y_i^D(t_j) = y_i(t_j) + \varepsilon_i, \quad \varepsilon_i \sim N(0, \sigma_i^2), \qquad (27)$$

where the standard deviation of the model error with dataset $i$, $\sigma_i$, is estimated simultaneously with the model parameters.

The error model in Eqn. (27), under the assumption that residuals are independent, gives the explicit log-likelihood of the model parameters, denoted by $\theta$:

$$l_D(\theta) = -\frac{1}{2}\sum_i \sum_j \left(\log(2\pi\sigma_i^2) + \frac{\left(y_i^D(t_j) - y_i(t_j)\right)^2}{2\sigma_i^2}\right), \qquad (28)$$

where $t_j$ denotes the experimental observation times.

We take a Bayesian approach to model calibration, in which uncertainty associated with model parameters ($\theta$) is quantified in a posterior distribution $P(\theta|\boldsymbol{y}^D)$. This posterior distribution can be calculated from Bayes' theorem

$$P(\theta|\boldsymbol{y}^D) \propto P(\boldsymbol{y}^D|\theta)\pi(\theta),$$

where $P(\boldsymbol{y}^D|\theta) = \exp\{l_D(\theta)\}$ is the likelihood of observing the measured data, and $\pi(\theta)$ is the prior distribution of the parameter vector $\theta$. In all cases, we choose uniform priors with lower bounds set equal to zero.

Posterior distributions are inferred using a Metropolis-Hastings MCMC (Markov chain Monte Carlo) sampler with adaptive proposal covariance. This is implemented in the parameter estimation toolbox PINTS[97]. In the MCMC algorithm, a Markov Chain starts at position $\theta$ and moves to a candidate position $\theta^*$ with acceptance probability $q = \min\{1, \mathrm{P}(\theta^*|\boldsymbol{y}_D)/P(\theta|\boldsymbol{y}_D)\}$. This process favours transitions to higher posterior probabilities but still allows moves to lower-probability regions to avoid local maxima. The resulting stationary posterior distributions are presented in the main text.

Plotting doubling times

Doubling times for the mathematical model were estimated by assuming that the population would grow exponentially if terms other than those related to proliferation were ignored. For the two-population model, increases in the total cell density are due to the A and B cell types. Summing the proliferative terms in the ODEs for these two populations and factoring out the sum $A(t) + B(t)$ in each layer leads to an equation that approximates the increase in the total cell density within each layer due to proliferation alone, without accounting for other mechanisms such as spreading or apoptosis. The doubling times of these equations for each cerebellum layer are approximately given by

$$T_W(t) = \frac{24\ln(2)}{\left(\alpha_1 + k_{w,1}E_W(t)\right)\frac{A_W}{A_W + B_W} + \left(\alpha_3 + k_{w,4}E_W(t)\right)\frac{B_W}{A_W + B_W}}, \qquad (32)$$

$$T_G(t) = \frac{24\ln(2)}{\left(\alpha_1 + k_{g,1}E_G(t)\right)\frac{A_G}{A_G + B_G} + \left(\alpha_3 + k_{g,4}E_G(t)\right)\frac{B_G}{A_G + B_G}}, \qquad (33)$$

$$T_M(t) = \frac{24\ln(2)}{\left(\alpha_1 + k_{m,1}E_M(t)\right)\frac{A_M}{A_M + B_M} + \left(\alpha_3 + k_{m,4}E_M(t)\right)\frac{B_M}{A_M + B_M}}, \qquad (34)$$

where all parameters and functions have the same interpretations as above. Doubling times have been multiplied by 24 to convert the time units to hours from days. Note that the doubling time depends on the relative fraction of A and B cell types and thus varies over time. Doubling times for the hippocampus layers are computed similarly.

<u>Model selection statistics</u>

We applied three different metrics to assess the quality of fits given by the MCMC procedures, and to compare the best fitting model between the logistic, linear, and two-population frameworks. The first, the Akaike information criterion (AIC), compares models using their log-likelihoods and penalizes models based on their potential to overfit the data[98]. The penalty term for the AIC is proportional to the number of parameters in the model. The AIC is calculated as

$$\mathrm{AIC} = -2l_D(\theta_{\mathrm{MLE}}) - 2\,N_p, \qquad (29)$$

where $N_p$ is equal to the total number of model parameters, $\theta_{\mathrm{MLE}}$ the set of parameter values giving the maximum log-likelihood (the maximum likelihood estimate, or MLE), and $l_D(\theta_{\mathrm{MLE}})$

its corresponding value. The AIC is on a log-scale, so there is no minimum value. Lower values (including negative numbers) indicate better fitting, more parsimonious models.

A second information-based metric is the Bayesian information criterion (BIC)[98,99]. Its name arises from its derivation using Bayesian techniques for selecting a model from the given data. The BIC gives the maximum value of this posterior distribution, and is given by

$$\mathrm{BIC} = -2l_D(\theta_{\mathrm{MLE}}) - 2\ln(n), \qquad (30)$$

where $n$ is equal to the total number of datapoints and all other parameters have the same interpretation as above. The BIC yields similar results as the AIC but performs better when the number of data points is small[98].

The final information-based metric that we consider is the Watanabe-Akaike information criterion (WAIC), also known as the “widely applicable” information criterion. Rather than calculate the log-likelihood of the entire data set, the WAIC instead computes the expected log predictive density, a measure of how likely the given model will predict a new observation[99]. This estimate is then penalized by a function that depends on the number of data points. The WAIC is given by

$$\begin{aligned}\mathrm{WAIC} \;=\; &-2\sum_i\sum_j \ln\left(\frac{1}{S}\sum_s p\big(y_i(t_j)\big|\theta_s\big)\right)\\ &+4\sum_i\sum_j\left(\ln\left(\frac{1}{S}\sum_s p\big(y_i(t_j)\big|\theta_s\big)\right)-\frac{1}{S}\sum_s \ln\Big(p\big(y_i(t_j)\big|\theta_s\big)\Big)\right), \qquad (31)\end{aligned}$$

where $S$ denotes the number of samples to generate the posterior distribution in the MCMC algorithm and $p\big(y_i(t_j)\big|\theta_s\big)$ the likelihood of obtaining the value $y_i$ at time $t_j$ from the mathematical model using the parameter set given by $\theta_s$. This is given by

$$p\big(y_i(t_j)\big|\theta_s\big) = \frac{1}{\sqrt{2\sigma_i^2}}\exp\left(\frac{\big(y_i^D(t_j)-y_i(t_j)\big)^2}{2\sigma_i^2}\right),$$

where $y_i^D(t_j)$ and $y_i(t_j)$ have the same interpretation as in Eqn. (28). As for the AIC and BIC, there is no lower limit to the WAIC due to the log scale used. Lower values (including negative ones) are more indicative of a better fitting and more parsimonious model.
